# Recovery of trait heritability from whole genome sequence data

**DOI:** 10.1101/588020

**Authors:** Pierrick Wainschtein, Deepti Jain, Zhili Zheng, TOPMed Anthropometry Working Group, NHLBI Trans-Omics for Precision Medicine (TOPMed) Consortium, L. Adrienne Cupples, Aladdin H. Shadyab, Barbara McKnight, Benjamin M. Shoemaker, Braxton D. Mitchell, Bruce M. Psaty, Charles Kooperberg, Ching-Ti Liu, Christine M. Albert, Dan Roden, Daniel I. Chasman, Dawood Darbar, Donald M Lloyd-Jones, Donna K. Arnett, Elizabeth A. Regan, Eric Boerwinkle, Jerome I. Rotter, Jeffrey R. O’Connell, Lisa R. Yanek, Mariza de Andrade, Matthew A. Allison, Merry-Lynn N. McDonald, Mina K Chung, Myriam Fornage, Nathalie Chami, Nicholas L. Smith, Patrick T. Ellinor, Ramachandran S. Vasan, Rasika A. Mathias, Ruth J.F. Loos, Stephen S. Rich, Steven A. Lubitz, Susan R. Heckbert, Susan Redline, Xiuqing Guo, Y.-D Ida Chen, Cecelia A. Laurie, Ryan D. Hernandez, Stephen T. McGarvey, Michael E. Goddard, Cathy C. Laurie, Kari E. North, Leslie A. Lange, Bruce S. Weir, Loic Yengo, Jian Yang, Peter M. Visscher

## Abstract

Heritability, the proportion of phenotypic variance explained by genetic factors, can be estimated from pedigree data ^1^, but such estimates are uninformative with respect to the underlying genetic architecture. Analyses of data from genome-wide association studies (GWAS) on unrelated individuals have shown that for human traits and disease, approximately one-third to two-thirds of heritability is captured by common SNPs ^2–5^. It is not known whether the remaining heritability is due to the imperfect tagging of causal variants by common SNPs, in particular if the causal variants are rare, or other reasons such as overestimation of heritability from pedigree data. Here we show that pedigree heritability for height and body mass index (BMI) appears to be largely recovered from whole-genome sequence (WGS) data on 25,465 unrelated individuals of European ancestry. We assigned 33.7 million genetic variants to groups based upon their minor allele frequencies (MAF) and linkage disequilibrium (LD) with variants nearby, and estimated and partitioned genetic variance accordingly. The estimated heritability was 0.68 (SE 0.10) for height and 0.30 (SE 0.10) for BMI, with a range of ~0.60 – 0.71 for height and ~0.25 – 0.35 for BMI, depending on quality control and analysis strategies. Low-MAF variants in low LD with neighbouring variants were enriched for heritability, to a greater extent for protein-altering variants, consistent with negative selection thereon. Cumulatively variants with 0.0001 < MAF < 0.1 explained 0.47 (SE 0.07) and 0.30 (SE 0.10) of heritability for height and BMI, respectively. Our results imply that rare variants, in particular those in regions of low LD, is a major source of the still missing heritability of complex traits and disease.

## Introduction

Natural selection shapes the joint distribution of effect size and allele frequency of genetic variants for complex traits in populations, including that of common disease in humans, and determines the amount of additive genetic variation in outbred populations^6^. Traditionally, additive genetic variation, and its ratio to total phenotypic variation (narrow-sense heritability) is estimated using resemblance between relatives, by equating the expected proportion of genotypes shared identical-by-descent with the observed phenotypic correlation between relatives^1,6^. Such methods are powerful but blind with respect to genetic architecture. In the last decade, experimental designs that use observed genotypes at many loci in the genome have facilitated the mapping of genetic variants associated with complex traits. In particular, genome-wide association studies (GWAS) in humans have discovered thousands of variants associated with complex traits and diseases^7^. GWAS to date have mainly relied on arrays of common SNPs that are in LD with underlying causal variants. Despite their success in mapping trait-associated variants and detecting evidence for negative selection^8,9^, the proportion of phenotypic variance captured by all common SNPs, i.e., the SNP-based heritability 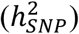, is significantly less than the estimates of pedigree heritability 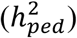^2,3^. Using SNP genotypes imputed from a fully sequenced reference panel recovers additional additive variance^5,10^, but there is still a gap between SNP-based and pedigree heritability estimates. The most plausible hypotheses for this discrepancy are that causal variants are not well tagged (or imputed) by common SNPs because they are rare and that pedigree heritability is over-estimated due to confounding of common environmental effects or non-additive genetic variation^3,11,12^.

Understanding the sources of the still missing heritability and achieving a better quantification of the genetic architecture of complex traits is important for experimental designs to map additional trait loci, for precision medicine and to understand the association between specific traits and fitness. Here we address the hypothesis that the still missing heritability is to a large extent due to rare variants not sufficiently tagged by common SNPs, by estimating additive genetic variance for height and body mass index (BMI) from whole genome sequence (WGS) data on a large sample of 25,465 unrelated individuals from the Trans-Omics for Precision Medicine (TOPMed) program.

## Results

### Heritability estimates of height and BMI using WGS data

#### Data overview and quality control

We used a dataset of 66,790 genomes (Supplementary Table 1) from which we selected a subset of 25,465 genomes with European ancestry by performing a two-step principal component analysis (PCA) on common and rare variants, using the 1000 Genomes^13^ and the Human Genome Diversity Panel^14^ as the reference panels (Online Methods; Supplementary Figure 1). We further removed outlier individuals based on their heterozygosity by grouping variants with similar minor allele frequency (MAF) and linkage disequilibrium (LD) characteristics (Online Methods, Supplementary Figure 2). After stringent quality control (QC), we retained 33.7M (out of 950M sequenced) variants, including 31.3M SNPs and 2.4M insertion-deletions (indels). We analysed variants observed at least 5 times in our dataset, which corresponds to a MAF threshold of 0.0001. The available phenotypes, height and BMI, were pre-adjusted for age and standardized to a mean of zero and a variance of 1 in each sex and cohort group (Online methods, Supplementary Figure 4). We also analysed both traits with a rank-based inverse normal transformation (height_RINT_ and BMI_RINT_) after adjusting for age and sex (Supplementary Notes, Supplementary Figure 6).

First, we used common variants known to be associated with height and BMI in European samples to test the consistency of predictive power of polygenic scores for each trait within each cohort, and the prediction results were consistent with those reported previously^15^ and those in the UK Biobank (UKB) (Online Methods, Supplementary Figure 7). Then, to verify that we could replicate prior estimates of 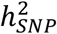 based on common SNPs, we selected ~992k HapMap 3 (HM3)^16^ SNPs from the sequence variants and estimated 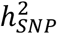 for height and BMI using the residual maximum likelihood analysis (GREML) approach implemented in GCTA^17^ We estimated an 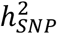 of 0.48 (SE 0.02) for height and 0.24 (SE 0.02) for BMI, again consistent with previous estimates^2,18^. To mimic a SNP-array plus imputation strategy, we imputed sequence variants in common with those available on three commonly used arrays to the Haplotype Reference Consortium (HRC) reference panels^19^ and estimated 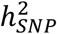 by stratifying the imputed variants according to MAF and LD using the GREML-LDMS approach^10^ implemented in GCTA (Online Methods, Supplementary Table 2 and Supplementary Table 3). We followed recommendations from Evans et al.^20^ for the LD annotation and therefore used SNP-specific LD metrics rather than segment-based metrics used in Yang et al.^10^ (Online Methods, Supplementary Figure 8, and Supplementary Notes). Estimates obtained using this imputation strategy are hereafter referred to as 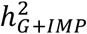. Estimates were in the range of 0.50-0.56 (SE 0.06-0.07) for height and 0.16-0.21 (SE 0.07) for BMI (Figure 1). When replacing the imputed SNPs with their sequenced genotypes, the estimates of 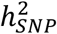 consistently increased (Supplementary Figure 10) with most of the differences coming from the variants with 0.0001 < MAF < 0.001 in the low-LD group, where imputation accuracy was the lowest. Overall, we largely replicate results from previous studies for common or imputed variants.

**Figure 1:**
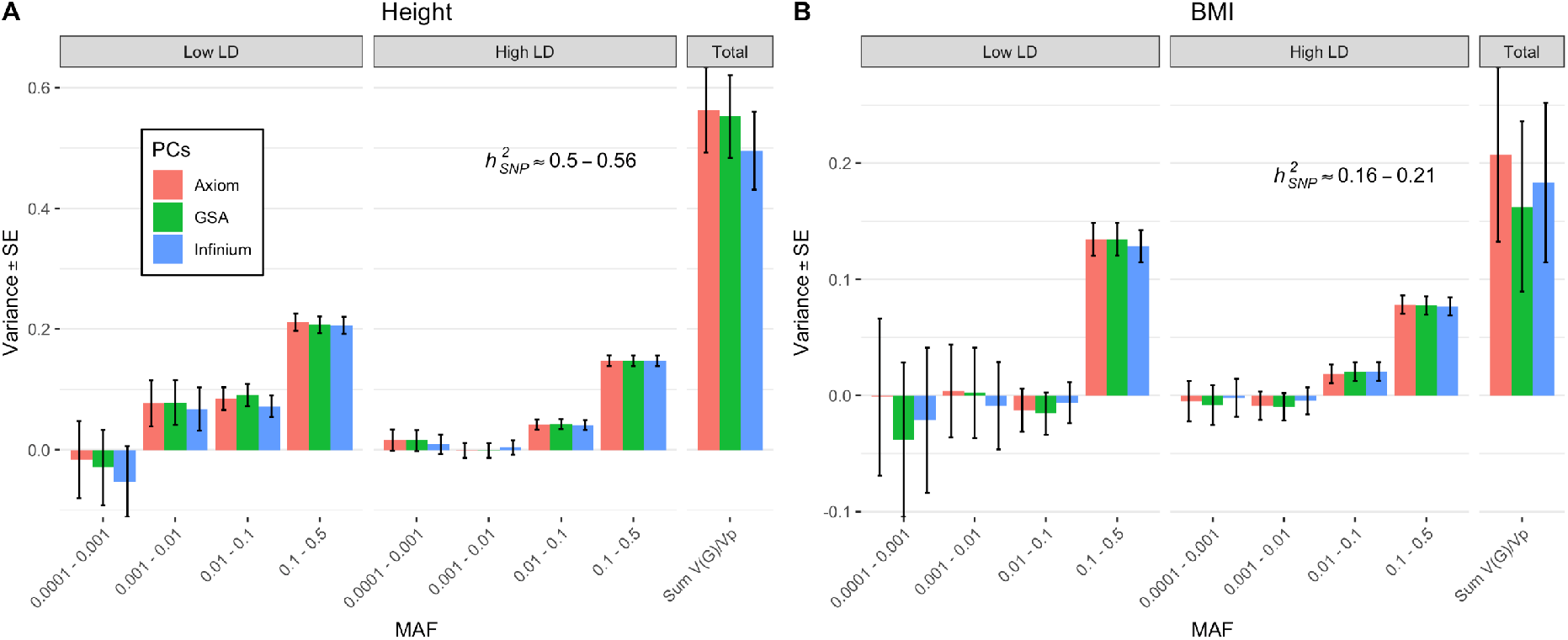
GREML-LDMS estimates with 8 bins (2 LD bins for each of the 4 MAF bins) correcting for 20 PCs (calculated from LD-pruned HM3 SNPs) after imputing SNPs from Illumina InfiniumCore24, GSA 24 and Affymetrix Axiom arrays using Haplotype Reference Consortium reference panels. (A) Estimates of 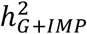 for height are between 0.50-0.56 (SE 0.06-0.07). (B) Estimates for BMI are between 0.16-0.21 (SE 0.07). The large SEs of the estimates for variants with MAF between 0.0001 to 0.001 can be explained by the large number of imputed variants in this MAF bin because the sampling variance of a SNP-based heritability estimate is proportional to the effective number of independent variants^35^. Between ~19.0M and ~20.0M variants in total are included in the analysis. The number of variants in each of the 4 MAF bins (twice the number in each LD bin) can be found in Supplementary Figure 9.

#### Estimation of trait heritability from WGS data

Having established that results from common or imputed variants were consistent with expectation, we then used all sequence variants with MAF > 0.0001 to estimate and partition additive genetic variance. We grouped variants according to MAF and LD (Supplementary Table 4, Supplementary Figure 11), using the GREML-LDMS partitioning method^10,20^ with a median-based LD grouping strategy (Online Methods). Estimates of heritability based on WGS data 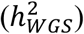 were consistently larger than 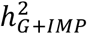. When correcting for the first 20 principal components (PCs) computed from HM3 SNPs, we found 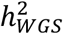 ~ 0.70 (SE 0.09) for height and ~0.29 (SE 0.09) for BMI (Supplementary Figure 12). The estimates for height are close to the pedigree estimates of 0.7-0.8 while this is not the case for BMI at 0.4-0.6, respectively^3,21^. We discuss below how much these results might be inflated by uncorrected population stratification.

The difference between 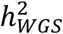 and 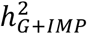 is predominantly explained by rare variants, in particular those in low LD with nearby variants. For the variants with MAF < 0.1, 0.31 of the phenotypic variance for height was accounted for by variants in the low-LD group but only 0.03 of the variance by variants in the high-LD group. For BMI, 0.05 of the phenotypic variance is accounted for by variants in the low-LD group and 0.03 from the ones in the high-LD group. Importantly, the large contribution of rare variants with low LD metrics could only be detected using WGS data as these variants are not present on SNP arrays and their imputation is not sufficiently accurate^22^. For both traits, our results confirm evidence for negative selection^8,9^ (Supplementary Figure 13).

### Impact of rare variant stratification

#### Linear regression of phenotype on PCs

We performed a number of additional analyses to test the robustness of estimates of 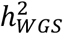. In particular, we conducted a series of analyses to attempt to quantify the contribution of any uncaptured population stratification to our estimates that is associated with rare variants. The standard correction method using PCs may not be effective in controlling for local population stratification in spatially structured populations, unless a large number of PCs are fitted in the analysis^23^. To estimate how much phenotypic variance is explained by PCs, we regressed the phenotype on a large number of PCs computed from common and rare variants (Online methods). This analysis showed that increasing the number of PCs from 20 (from common variants) to 160 (common and rare variants) modestly increased the regression R^2^ from 0.012 to 0.018 for height and from 0.001 to 0.004 for BMI (Supplementary Figure 14). These results imply that population structure, as quantified by linear regression on PCs from common and rare variants, in total may contribute only about 1.5% of phenotypic variance for height and nearly none for BMI. However, fitting them as fixed effects in a linear mixed model might not affect heritability estimates commensurately as shown below.

#### Using birthplace coordinates to capture potential population stratification in the UKB

We lack any direct information on spatial substructure in the TOPMed dataset, and therefore turned to the UKB where such information is available. We selected a sample of 35,867 unrelated individuals of European ancestry with both whole-exome sequence (WES) data and birthplace coordinates available (Online methods). We investigated how well the potential population stratification could be captured by the birth coordinates in the UKB. The estimates from a GREML-LDMS analysis fitting 14 MAF/LD bins (i.e., 7 MAF x 2 LD) from the WES data and an additional bin of HM3 common SNPs from imputed data were 0.62 (SE 0.04) for height and 0.33 (SE 0.04) for BMI. When including the birthplace coordinates as fixed covariates in the GREML-LDMS analysis, the estimates were essentially the same, 0.61 (SE 0.04) for height and 0.33 (SE 0.04) for BMI (Supplementary Figure 15), showing either no evidence of a strong effect of local stratification on the GREML estimates for rare WES variants in the UKB or that fitting birth coordinates does not fully capture population stratification. Estimates between TOPMed and UKB WES datasets were also similar when selecting variants present in both datasets after equalising the sample sizes (Online methods, Supplementary Figure 16), although we observed UKB WES-based estimates were consistently higher, possibly because the UKB sample is genetically more homogenous than our TOPMed sample.

#### Correcting for population stratification

We then quantified the number of PCs that are necessary to correct for population stratification within each MAF/LD bin using a novel approach. By further dividing SNP groupings based on their location on either the odd- or even-numbered chromosomes, we computed two sets of PCs for each LD/MAF bin (Online methods). Any variance explained by fitting a PC from one set of chromosomes on the other set would capture inter-chromosomal correlations, an indication of population stratification in the absence of relatedness. We did not have sufficient ancestry information to directly quantify how much this strategy can detect and correct for the effect of spatial substructure in the TOPMed dataset, and again used UKB data where such spatial information is available. We estimated the number of relevant PCs to account for population stratification by quantifying the inter-chromosomal correlations, (Supplementary Figure 17). By using birth coordinates and PCs of the UKB samples, we could visualise the stratification on a map (Supplementary Figure 18) and quantify it using Moran’s index of spatial autocorrelation (Supplementary Figure 19). From these analyses on the UKB data we concluded that our approach to discover which PCs are necessary to account for population stratification is appropriate. After performing the same analysis on TOPMed samples, we identified 48 PCs across the 8 MAF/LD bins that could fully account for population stratification between sets of chromosomes. We then used those PCs computed from independent variants in the GREML-LDMS analyses, which decreased estimates of 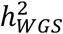 from 0.70 (SE 0.09) to 0.60 (SE 0.09) and from 0.29 (SE 0.10) to 0.23 (SE 0.10) for height and BMI respectively (Supplementary Figure 12), suggesting the presence of population stratification effects not captured by the 20 common variants PCs used in the analysis above. Importantly, biases in heritability estimates due to uncorrected population stratification do not match the actual proportion of trait variance that it explains. Finally, we repeated the GREML-LDMS analyses fitting 160 PCs (i.e., 20 PCs computed from each of the 8 MAF/LD bins) and observed very similar estimates to fitting only the 48 PCs identified from the inter-chromosomal correlations, indicating that population stratification has been accounted for when using the 48 PCs.

#### Influence of SNP quality and GRM estimators

As a further check, we compared GREML-LDMS estimates by selecting the complete set of genotyped variants in place of the high-quality variants identified with a classifier trained using a support vector machine algorithm (SVM) and additional hard filters (Online Methods). That increased the total number of variants used to compute GRMs from 33.7M to 44.2M variants. The estimate of 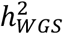 for height was 0.62 when fitting the 20 HM3 PCs and 0.63 when fitting the 160 PCs. When investigating the differences between these estimates and the ones from high quality variants, we noticed some large differences in diagonal and off-diagonal values of the low MAF GRMs, potentially due to sequencing errors or batch effects in these ultra-rare variants (Supplementary Figure 20, Supplementary Figure 21). We therefore concluded that the choice of stringent variant QC was justified.

To quantify the effect of differential SNP weighting within each GRM, we investigated the influence of calculating the GRM estimator *A_jk_* using the average ratio over loci (“average of ratio”)^17^ instead of the ratio of total SNP covariance and total SNP heterozygosity over loci (“ratio of averages”)^24^ on the GREML-LDMS estimates (Online Methods, Supplementary Figure 22). Note that the default GCTA method (average of ratios) assumes an inverse relationship between MAF and variant effect size whereas the ratio of averages assumes no relationship between MAF and variants effect sizes. For height and BMI, the average of ratio method showed slightly higher estimates (0.63, SE 0.10 for height and 0.29 (SE 0.11) for BMI, respectively) than the ratio of averages (0.61 (SE 0.09) and 0.25 (SE 0.10)), accounting for population stratification. These results show that our current estimates seem robust to the GRM estimator (see Discussion and the Supplementary Notes for more discussion), consistent with prior work^10^.

#### Investigating the influence of direct GRM QC on heritability estimates

We finally investigated how outlier values in the rare variants GRMs influence the estimates of 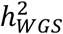. From the 28,755 unrelated individuals of European ancestry (before filtering individuals based on their heterozygosity values in each bin, Supplementary Figure 2), we filtered individuals based either on their off-diagonal values (removing pairs > 0.1), diagonal values (<0.7 and >1.3), or both (Online Methods, Supplementary Figure 23). For height, removing one of each pair of individuals with large off-diagonal values in any of the rare variants GRMs did not change the estimates regardless of the number of PCs fitted in the model (~0.58 – 0.71 (SE 0.08)). However, excluding individuals with extremely large diagonal values yielded a substantial increase in 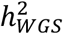 estimates (0.71 – 0.81 (SE 0.09)). Removing individuals based on both diagonals and off-diagonals values still showed this large increase in heritability estimates (0.71 – 0.82 (SE 0.09 – 0.10)). For BMI, filtering samples did not increase the estimates but revealed a larger contribution from rare variants in the low-LD bins; and a large overall increase was observed compared to estimates from QC based on sample heterozygosity.

#### Impact of LD annotation

We first used in-sample or out-sample LD/MAF from the UK10K dataset^25^ and observed consistent heritability estimates suggesting that our inference from TOPMed annotation is not biased by using MAF and LD stratification from another dataset (Supplementary Notes, Supplementary Figure 24). We then investigated the influence of LD on GREML-LDMS estimate by increasing the number of LD bins. Prior work suggest that 2 LD bins are sufficient to estimate and partition genetic variance^10^. However, when using WGS variants there is substantial variation in LD score within a MAF group (Supplementary Figure 25). We therefore investigated whether further partitioning according to LD could better capture genetic variance. We divided each MAF bin into 3 and 4 LD bins instead of 2, thereby increasing the total number of GRMs fitted in the model from 8 to 12 (when dividing each MAF bin into 3 LD tertile groups) or 16 (when dividing each MAF bin into 4 LD quartile groups). Compared to the 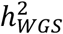 estimates based on 2 LD bins, the estimates based on 3 and 4 LD bins were consistently higher, ranging between 0.67 – 0.68 (SE 0.10) for height and 0.28 – 0.32 (SE 0.10) for BMI (Figure 2), with the 48 PCs fitted as fixed covariates. This increase is consistent with larger LD heterogeneity within MAF bins, which was not fully accounted for by 2 LD bins. While using 3 LD bins substantially increased estimates for both traits, we did not observe such increase when increasing the number of LD bins to 4 (Supplementary Figure 26). Comparing the Akaike Information Criterion (AIC) between different models, the 4 LD grouping strategy consistently performed the best for height and BMI, followed by the 3 LD grouping indicating that the 2 LD grouping strategy is not the most appropriate when considering rare variants (Supplementary Figure 27). These estimates are also closer to the estimates of 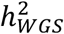 without outlying diagonal and off-diagonal elements (see above), indicating that increasing the number of LD bins might be an effective way to account for rare variants heterogeneity between individuals. However, additional partitioning comes at the cost of decreased precision of the estimate of heritability.

**Figure 2:**
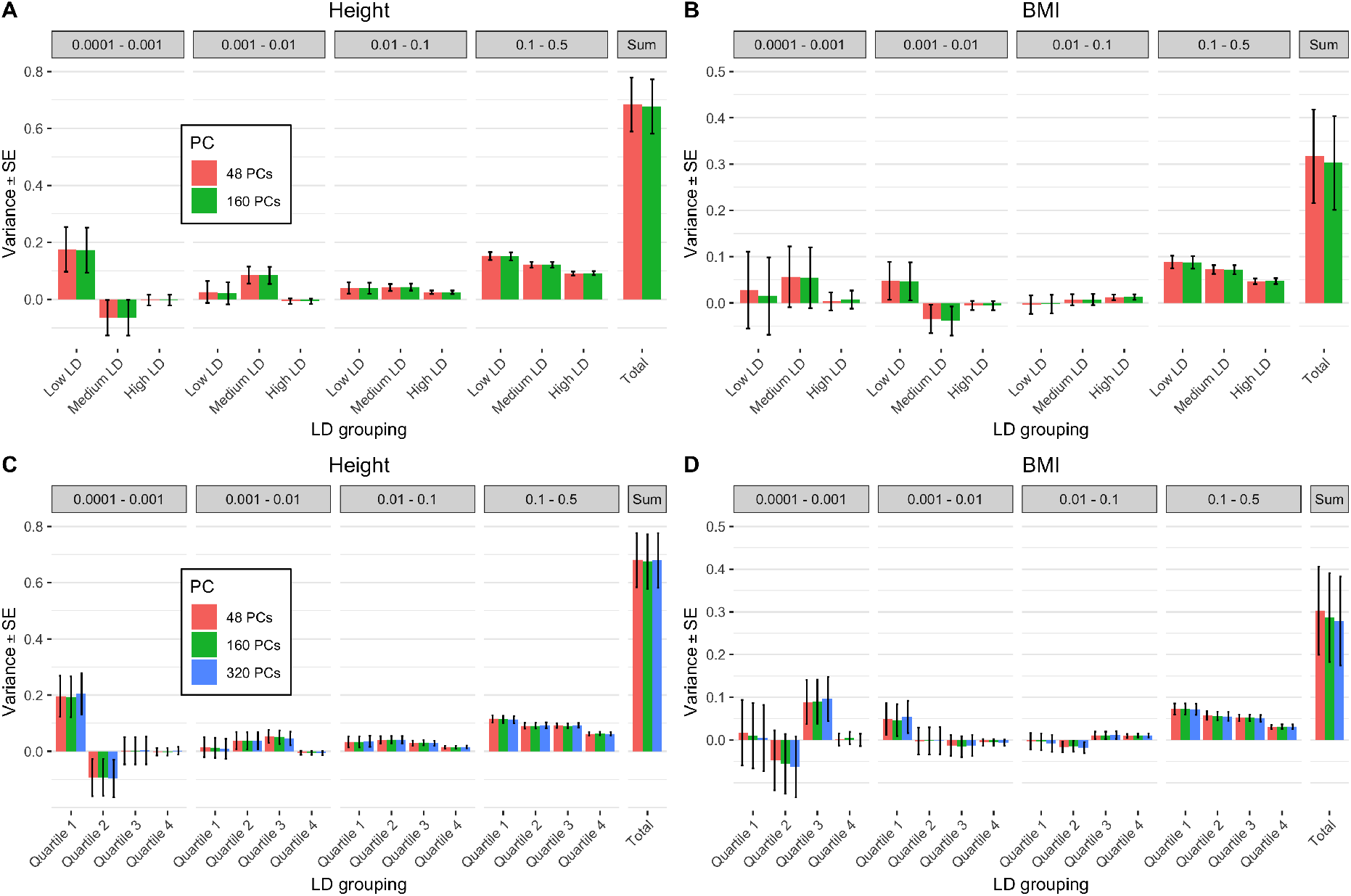
GREML-LDMS of height and BMI for N=25,465 samples using 3 or 4 LD groups for each MAF bin correcting for 48/160/320 PCs computed from WGS variants. Each variant was allocated in a tertile or a quartile according to its LD score. (A) Estimates using 3 LD bins for height: 0.68 (SE 0.09 – 0.10). (B) Estimates using 3 LD bins for BMI: 0.30 – 0.32 (SE 0.10). (C) Estimates using 4 LD bins for height: 0.67 – 0.68 (SE 0.10). (D) Estimates using 4 LD bins for BMI: 0.28 – 0.30 (SE 0.10).

### Functional annotation

Having estimated and partitioned additive genetic variance from WGS data and investigated the robustness of the estimates, we then explored whether the variance could further be partitioned by functional genomic annotations. To investigate the specific contribution of low-LD variants with MAF < 0.1 to heritability, we partitioned the low-MAF and low-LD variants bins further according to the putative effect of a variant on protein coding using SnpEff^26^. The protein-altering group comprises loss of function and non-synonymous variants whereas the remaining variant set comprises synonymous, regulatory or non-coding variants (Online methods, Supplementary Table 5). The proportion of protein-altering variants was different across the LD and MAF groups, with an increase in low MAF bins (Supplementary Figure 11), consistent with purifying selection on this class of variants^9^. Overall, we considered a total number of 11 groups (2 bins for variants with MAF > 0.1 and 3 MAF bins for variants with MAF < 0.1; each MAF bin is further split into 3 groups: low LD protein-altering, low LD non-protein-altering and high LD). When running a GREML-LDMS analysis with these 11 bins fitting the 48 PCs, the total estimates remained similar for height 0.61 (SE 0.09) and slightly increased for BMI 0.24 (SE 0.10), although these estimates are biased downward owing to the use of only 2 LD bins (further splitting by LD and variant effect would lead to fitting too many random effects in the analysis). Interestingly, the average variance explained per variant was larger for bins with protein-altering variants (low-LD) compared to bins with non-protein-altering variants (low-LD) or high-LD variants (Figure 3).

**Figure 3:**
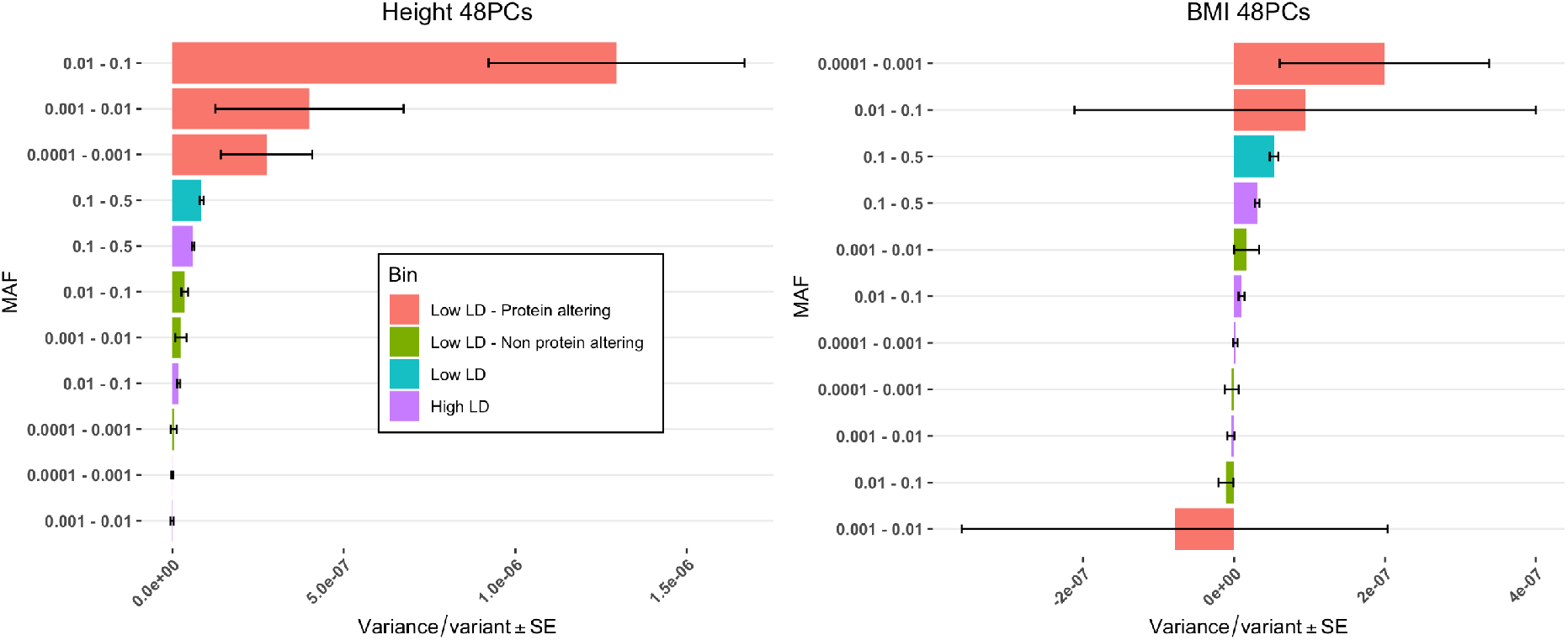
Variance explained per variant (the estimate of genetic variance divided by the number of variants in each bin) from GREML-LDMS with the low-LD and low-MAF (<0.1) variants partitioned into 2 distinct categories according to the SnpEff putative effect of the variant (protein-altering or non-protein-altering), correcting for 48 PCs from WGS variants. There is a total of 11 genetic components in this analysis. There is an apparent enrichment of heritability in the protein-altering groupings (low LD) over non-protein-altering (low LD) or high LD variants for height (A) as well as for BMI, although the standard errors for this trait are large (B).

## Discussion

### Estimates of heritability from WGS on height and BMI

We have used the largest sample to date with both WGS data and phenotypes to estimate the heritability of height and BMI captured by rare and common variants sequenced in a sample of 25,465 unrelated individuals from the TOPMed consortium. Our estimates largely but not fully recover the heritability estimated from pedigree data. We observed an additional variance detected over and above SNP arrays or imputation due to rare variants, in particular rare protein-altering variants in low LD with other genomic variants.

### Follow-up analysis performed to assess the robustness of our estimates

To assess the robustness of these results, we conducted several follow-up analyses. We estimated the variance explained by correcting for a large number of principal components (up to 160) which may minimize any bias due to population stratification. We used a LD and MAF reference from another European ancestry dataset with whole genome sequences (Supplementary Notes), and furthermore investigated the robustness of our estimates with alternative LD partitioning. All three analyses confirmed the validity of our estimates as additional variance is systematically detected from rare variants not previously tagged by imputation methods. We also estimated heritability for height and BMI using another GRM estimator and confirmed the robustness of our statistical framework. Moreover, comparing 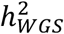 with 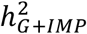 allowed us to demonstrate that most of the heritability due to very rare variants (0.0001 < MAF < 0.001) was missing when using imputed data but revealed by using WGS data. We evaluated the loss of information on variance component estimates due to imputation compared to a similar variant coverage of WGS data and found negligible differences in the estimates of genetic variance. We investigated further the enrichment in heritability for different types of variants (high or low impact on the protein) and showed that for low-LD variants with MAF < 0.1, non-synonymous and protein-altering variants are more enriched for trait heritability than synonymous or non-coding variants, as shown in previous studies^4,27^.

### Impact of population stratification on WGS estimates

By investigating the variance explained by principal components from one chromosome set onto the other, we identified inter-chromosomal correlations among rare variants, indicating residual stratification in the sample. Inter-chromosomal correlation for rare variants is to be expected as it has been shown that recent population growth resulted in an excess of rare variants in European populations^28,29^, and our analyses show that PCs derived from common variants only are insufficient to account for it. We used the UKB exome dataset to detect and quantify such a stratification in a sample independent of the TOPMed. Although we would expect a larger stratification in TOPMed than in the UKB samples, the similar inter-chromosomal correlation patterns show that we can capture it through fitting PCs as fixed effects in a GREML-LDMS analysis. The concordance of the GREML-LDMS estimates fitting either 48 PCs identified through the chromosome analysis or a larger number of PCs (e.g., 160, 20 PCs per bin) indicates that population stratification can be accounted for by fitting PCs. Population stratification could bias estimates of heritability if it is correlated with environmental effects. Regression of the phenotypes on PCs accounted for little variance explained even when fitting a large number of PCs calculated from rare variants. Similarly, there is no bias due to environmental effect associated with birthplace spatial coordinates using the UKB exome data. Nevertheless, the absence of any such effect in the UKB does not provide direct evidence of its absence in the TOPMed sample.

### Potential influences of extreme GRM values

One potential bias seems to be related to extreme values in the diagonal elements of the rare variants GRMs. When removing individuals with high diagonal values, we saw an increase in 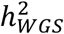 estimate to levels that recover most of the remaining missing heritability, with a large contribution from the low-LD rare variants. This was not observed when removing individuals based on their off-diagonal elements. Differences between individuals in their diagonal values can reflect inbreeding, population structure and sampling variance. For common variants, the contribution to the estimate of genetic variance from differences in diagonal values is dwarfed by that from the off-diagonals, but this is not the case for rare variants (Supplementary Figure 28). Large diagonal values would bias the estimates downwards if they are not correlated with increased genetic variance. Further work is needed to fully understand the causes and consequences of the variance of diagonal values in GRMs estimated from rare variants.

### Potential bias from assortative mating

Height is known to drive positive assortative mating in the population^30^, which implies that the genetic variance in the current population is larger than that in a randomly mating population^1^. Although the difference between the true heritabilities in randomly and assortatively mating populations is typically small, recent findings using SNP array data have shown that estimates from GREML can be biased upwardly for traits undergoing assortative mating, and that it approaches the random mating heritability only for very large sample sizes^31^. We do not know how much our estimates from WGS data on height are biased because of assortative mating.

### Other sources of variance

The experimental design to estimate trait heritability that is least biased by assumptions is to use sibling pairs and their realized identity-by-descent relationships estimated from marker data^32^. Recently, Kemper et al. reported an estimate of 0.81 (SE 0.10) for the heritability of height in the current population using this design^33^. Our GREML estimates for height accounting for population stratification are in the range of 0.60 to 0.70. Therefore, even if we consider the largest estimates then there is still a gap between those and the within-family estimate of heritability. One explanation for this discrepancy could be sampling variance, since our estimation errors are about 0.09 – 0.10 and that from Kemper et al. is even larger. On the other hand, while our research shows a large role of low-LD rare (0.0001 to 0.01) variants, variants with MAF<0.0001 (including singletons,^34^ which were ignored in our study), and structural variants, which are not well-captured by short-read WGS data, could contribute to trait variance and explain the apparent still-missing heritability. In addition, we only studied the autosome, and would expect the X chromosome to contribute a small amount of genetic variation. Larger sample size for both the sibling and WGS population designs are needed to resolve these outstanding questions.

### Quality control and analysis considerations

Our results lead to a number of important QC and analysis considerations when estimating and partitioning genetic variance from single-ancestry WGS data. Stringent quality control should be performed at multiple levels, including variant- and individual-level genotype QC by selecting high quality variants and rejecting individuals with outlying genome-wide heterozygosity; population stratification QC, by selecting individuals from the targeted ancestry using multiple rounds of global and local ancestry projections; and common variant trait association QC, by validating that prediction accuracy and SNP-based heritability are consistent with prior reports on the same trait, ideally in the same ancestry. Individual and pairwise sample QC could also be performed by estimating identity-by-descent segments and on properties of genomic relationship matrices. Finally, for the actual analysis it is important to fit a sufficient number of principal components estimated from both common and rare variants, and to stratify variants in as many LD, MAF and functional annotation strata as the sample size allows.

### Limitations of our analyses

Our estimates of heritability from WGS data have standard error of about 9-10%. Since standard errors are approximately inversely proportional to sample size^35^, doubling the sample size to 50,000 would narrow errors to ~5% and would allow further and more precise partitioning of genetic variance. Until now the question of the contribution from rare variants to the missing heritability could only be investigated through imputing genotypes from WGS reference panels that was subject to imperfect tagging. Our results quantify this contribution and allows for recovery of some of the remaining missing heritability. It would be interesting to further partition the genome (by variant type, predicted variant deleteriousness^36^ and more LD/MAF groupings), but standard errors of the estimates would be too large given our current sample size. Similarly, with a larger sample size, contribution to the heritability from assortative mating could be quantified^37^ The contribution of rare variants to narrow-sense heritability, larger than expected under a neutral model, also reinforces previous observations that height- and BMI-associated variants have been under negative selection^8–10^, although population expansion could also lead to an increase in heritability from rare variants^38^. Once again, a larger sample size would allow us to draw stronger conclusions on the selective pressure on the genetic variants associated with the two traits. Additionally, the TOPMed samples come from multiple cohorts across the US are potentially subject to multiple and diverse non-genetic effects. Some cohorts of the TOPMed program are ascertained towards diseases correlated with high BMI values. Having case-control cohorts as part of the larger analysis might affect the robustness of the estimates. Finally, our sample for analysis was restricted to a single ancestry. Since genetic architecture and heritability are per definition population specific, future analyses using data from other ancestries will reveal how generalisable our results are.

### Implications

Our results have important implications for the still missing heritability of many traits and diseases^3^. Indeed, the ratio of SNP to pedigree heritability for diseases is lower than for height and BMI, leading to potentially more discovery from rare variants contributing to diseases using WGS data. Our results are also important for polygenic scores as using WGS data could, in principle, lead to predictors with larger prediction accuracy for many polygenic diseases. With the cost for sequencing still much higher than array genotyping, large scale WGS data acquisition is currently limited to national initiatives such as the TOPMed and other programs but is bound to expand in the next decade. Large cohorts of genotyping array data will still prove useful for gene discovery or predictions of common diseases and should complement WGS data for a broader understanding of genetic architecture. In the future, WGS programs for specific diseases on large cohorts could lead to a large increase of low MAF variants identified. Sample sizes required to detect such variants from genome-wide association studies using sequence data are of the same order of magnitude as current well-powered GWAS on common variants, i.e., hundreds of thousands to millions of individuals.

## URLs

GCTA, http://cnsgenomics.com/software/gcta/#Overview. Plink, https://www.cog-genomics.org/plink2. UK10K, https://www.uk10k.org/. NHLBI, www.nhlbiwgs.org. TOPMed methods, https://www.nhlbiwgs.org/topmed-whole-genome-sequencing-project-freeze-5b-phases-1-and-2.

## Online Methods

### Data collection

In this study, we used Whole-Genome Sequence (WGS) data from the Trans-Omics for Precision Medicine (TOPMed) Program. The TOPMed program collects WGS data from different studies and centers, in the United States and elsewhere, in partnership with the National Heart, Lung, and Blood Institute (NHLBI, see URLs). The “freeze 8” version of the data includes 140,306 samples containing ~920M SNPs and indels in the called variants files (as BCF, binary variant call format). These variants have been called on genome assembly GRCh38 as human genome reference (see URLs for methods). Participant consent was obtained for each of the 18 studies (Supplementary Table 1, Supplementary Table 6) containing Europeans samples in the freeze 8 as well as the associated phenotypes for height and BMI.

### Quality control

We selected freeze 8 samples with height and BMI phenotypes available (N=66,790). After removing individuals under 18 years old, we had 64,930 adults left. For each sex within each of the 18 different studies included in this analysis (each cohort), we regressed the height and BMI according to their age and kept the residuals. Moreover, to remove differences in mean and standard deviation between sexes and among cohorts, we standardized the residuals by the standard deviation of each sex and cohort. The standardized residuals on height and BMI of each gender group of each cohort followed a distribution N(0,1) (Supplementary Figure 4). We also applied a rank-based inverse normal transformation on the height and BMI (Height_RINT_ and BMI_RINT_) after adjustment for age and sex. On the genotypes, we performed a multi-step quality control. We first selected the samples with phenotypes available (N=66,790) and retained only the high-quality variants that passed a SVM classifier. The SVM classifier was trained using variants present on genotyping arrays labelled as positive controls, and variants with many Mendelian inconsistencies labelled as negative controls. We then excluded variants with genotypes missingness rate > 0.05, Hardy-Weinberg equilibrium test *P* value <1 × 10^-6^, or with a minor allele frequency <0.0001 using PLINK v1.9 (see URLs^39^). To select samples of European ancestry, we performed a similar QC on the 2,504 samples of the 1000 Genomes Project (with a MAF threshold of 0.004, to account for the difference in sample size between datasets). On the 1000 Genomes genotypes, we used different parameters to select two sets of independent variants for common (M= 1,495,743, MAF range of 0.01 to 0.5, window size of 50kb and R^2^ threshold of 0.1) and rare variants (M= 1,512,042, MAF range of 0.004 to 0.01, window size of 100kb and R^2^ threshold of 0.05). We computed 20 principal components on the 1000 Genomes samples using common and rare variants and then used the variants loadings to project TOPMed samples on the PCs (with respectively 579,015 and 1,268,148 variants matching) (Supplementary Figure 1).

TOPMed samples were classified as of European ancestry if their Euclidean distance, based on the first 4 PCs computed from common variants weighted by their respective eigenvalues, to the cluster of the 1000 Genomes samples of European ancestry was lower than 3 SD of the within-cluster distance (N=37,212 samples left). We then performed similar filtering on the first 4 PCs computed rare variants, which removed extra 274 samples (N=36,938). To further remove ancestral outliers not captured by PCs, we used RFMix to infer local ancestry from 639,958 autosomal SNPs on 938 individuals (grouped in 7 super-populations) from the Human Genome Diversity Panel and removed 2,900 individuals with the inferred local ancestry more than 3 standard deviations away from the mean of European, East-Asian, African and North-American populations (Supplementary Figure 3).

For the remaining 34,038 individuals, we built a genetic relatedness matrix using variants on HM3 reference panels with a MAF > 0.01 and removed one of each pair of individuals with estimated genetic relatedness > 0.05, resulting in 28,755 unrelated individuals.

The last QC step was to remove individuals showing an excess of heterozygosity after we noticed individuals showing short IBD segments from different ancestries but having a high impact on the off-diagonal elements of the GRM for the rare variant bins (see Supplementary Notes and Supplementary Figures 29 to 35). As described above, we stratified the variants by MAF and LD into 8 bins and computed the sample heterozygosity using the variants in each bin. To obtain a uniform distribution of heterozygosity across MAF range, we performed 4 rounds of filtering, removing samples 3 standard deviations away from the mean heterozygosity of the samples within each MAF and LD bin (Supplementary Figure 5).

At the end of all the quality control steps, we retained 25,465 unrelated individuals of European ancestry and 33.7 million variants (MAF and LD distributions of the variants are shown in Supplementary Figure 11). We also computed a 2^nd^ dataset with the same samples but using all the genotyped variants with a MAF > 0.0001 (not only the SVM-filtered high-quality ones). We maintained SNP QC for each step of the QC process and also recomputed the LD score with the final samples.

### Polygenic scores from common variants

To perform joint genotype-phenotype QC, we constructed a polygenic score (PGS) for each trait using trait-associated variants identified in the UKB. We performed GWAS of height and BMI in N=400,831 European ancestry participants of the UKB using fastGWA^40^. We then selected 1362 and 452 independent (LD r^2^<0.01) genome-wide significant SNPs (P<5×10^-8^) to calculate PGS of height and BMI, respectively, of which 1360 and 449 SNPs were matched to our TOPMed data. For each phenotype, we regressed the standardized phenotype on the mean-centred PGS and computed the regression slope and the variance explained by the predictor. To make a direct comparison with a non-TOPMed dataset we performed the same analysis using a hold-out sample of 14,587 UKB participants independent of our GWAS discovery set. The regression of the predicted trait on its PGS gave a slope of ~0.90 for height and ~0.83 for BMI, with a corresponding variance explained of 0.25 and 0.04, respectively.

### Statistical framework of the GREML analysis

The GREML analysis is based on the idea to fit the effects of all the variants as random effects in a mixed linear model (MLM),

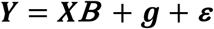

where ***XB*** is a vector of fixed effects such as age, sex and in our case the principal components of each subset of variants, ***g*** is an n x 1 vector of the total genetic effects captured by all the sequenced variants of all the individuals (with n being the sample size) and ***ε*** is a vector of residuals effects. From MLM, ***g*** follows a distribution 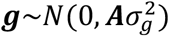 where ***A*** is a GRM interpreted as the genetic relationship between individuals. We estimate 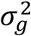 using the REML algorithm^17^

The genetic relationship between individuals *j* and *k* (*A_jk_*) was estimated by the following equation:

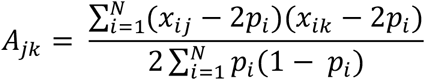

where *x_ij_* (*x_ij_* takes value in 0,1 or 2) is the minor allele count for SNP *i* in individual *j*, *N* is the number of variants, and *p* is the sample minor allele frequency.

We refer to this method of estimating pairwise relationships as the ratio of total SNP covariance and the total SNP heterozygosity over loci, also called the ratio of averages^24,41^. By using the sample allele frequencies, *A_jk_* does not represent a measure of kinship between two individuals, although the GRM should be highly correlated with the kinship matrix if we were to have full and accurate pedigree data on the entire sample^24^.We calculated multiple GRMs based on subsets of SNPs (stratified by MAF, LD, annotations, etc) and fit them as random effects according to a more general model:

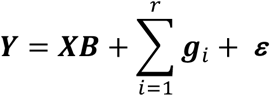

where the phenotypic variance 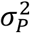 is the sum of the residual variance and the variance of each of the *i^th^* genetic factor (each with a corresponding GRM).

To compare the methods to calculate the genetic relationship between individuals *j* and *k* we also used an estimator where the ratio is the SNP covariance divided by SNP heterozygosity (the average of ratios) from the following equation:

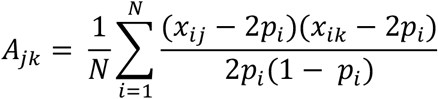

It has been shown previously that this estimator is accurate among unrelated samples, but does not perform as well as the ratio of averages estimator when there is relatedness among samples^24^, as we have when calculating GRMs from very rare variants.

### Proportion of genetic variation captured by imputation

Previous REML estimates based on rare variants were conducted by imputing SNP chip data on a reference panel such as 1000 Genomes^10^. To check the consistency of our dataset with previous estimates, we mimicked SNP chip data by selecting SNPs in our dataset present on Illumina InfiniumCore24 v1.2, GSA 24 V3.0 and Affymetrix UKB Axiom arrays. We downloaded the list of SNPs of these arrays, the UCSC to Ensembl reference and the

GRCh37 reference. With those, we selected subsets of the SNPs in common between the TOPMed dataset and each of the SNP arrays. After this merging and cleaning process, we retained the majority of the SNPs present on each array (Supplementary Table 2). We then imputed our dataset on the Michigan imputation server^19^. Imputation was performed in multiple stages. Prior to imputation, each SNP coordinate was lifted from hg38 to hg19. Then each chromosome was phased against Europeans from HRC.r1.1 ^19^ reference using EAGLE2 v2.4 ^42^. In a second stage, data were imputed using Minimac 4. On the imputed datasets, we removed variants with imputation info score < 0.3, missingness rate > 0.05, Hardy-Weinberg equilibrium test *P*-value <1 × 10^-6^, or MAF < 0.0001 and individuals with missingness rate > 0.05. After imputation and filtering, we had between ~19 and 20M SNPs left in each imputed dataset (Supplementary Table 3 and Supplementary Figure 9). In each imputed dataset, we stratified SNPs into 4 MAF bins (0.0001<MAF<0. 001, 0.001<MAF<0.01, 0.01<MAF<0.1, 0.1<MAF<0.5). For each of the 22 autosomes, we calculated the LD score of each variant with the others on a sliding window of 10Mb using GCTA software^17^ We performed two types of LD binning, selecting variants based on their individual LD scores or on their segment-based LD scores (segment length = 200Kb) (Supplementary Figure 8). Each of the 4 MAF bins was divided into 2 more bins, one for variants with LD scores above the median value of the variants in the bin (high LD bin) and one for variants with LD score below median (low LD bin) (Supplementary Table 4). We then used GCTA to perform a GREML-LDMS analysis with the first 20 PCs calculated using HM3 SNPs from the WGS dataset fitted as fixed effects and the variants in the 8 MAF and LD bins as 8 random-effect components (Figure 1).

To assess the influence of imputation errors on variance estimates, we selected, for the dataset imputed from the SNPs on the Axiom array, the SNPs that were available in both the imputed UKB data and the TOPMed WGS data. We had 17.9M SNPs in both the TOPMed and the imputed datasets. In each of these pairwise datasets (a set from the TOPMed WGS and another set from the imputed UKB), we partitioned SNPs in 8 bins, according to their MAF and LD scores, similarly to the analysis above. We then ran a GREML-LDMS analysis on height and BMI in each dataset with either 20 principal components calculated from HM3 SNPs or 160 PCs (20 PCs computed from each of the 8 MAF/LD bins) fitted as fixed covariates.

### GREML estimates from WGS data

Before estimating the proportion of phenotypic variance due to additive genetic factors from WGS data we initially wanted to check for consistency with previous studies and performed a single-component GREML analysis (GREML-SC approach) in GCTA using HM3 SNPs with 20 PCs. We calculated the PCs from LD-pruned SNPs and fitted them as fixed effects in the GREML-SC analysis. Note that we chose the GREML-LDMS analysis to estimate heritability of height and BMI when using the entire dataset for the following reasons. It has previously been shown^10^ that a GREML-SC approach can give a biased estimate of *h*^2^ if causal variants have a different MAF spectrum from the variants used in the GREML analysis. Moreover, if two variants in the same GRM have different LD properties, this can also lead to biased estimates of heritability. We performed a GREML-LDMS analysis using 4 MAF bin (0.0001<MAF<0. 001, 0.001<MAF<0.01, 0.01<MAF<0.1 and 0.1<MAF<0.5) and 2 LD bins. Similar to what was done using data imputed from array SNPs, we defined 8 bins by splitting each of the 4 MAF bin into two LD bins (Supplementary Figure 12). Within each of these 8 bins, we calculated 20 PCs that we fitted together as fixed covariates to correct for population stratification (160 PCs in total). To investigate how low-quality variants would bias estimates, we also conducted an analysis including the variants that did not pass a support vector machine (SVM) classifier or additional filters on excess of heterozygosity and Mendelian discordancy.

To investigate the robustness of the assumptions on the relationship between MAF and effect size, we ran a GREML-LDMS analysis on the previously defined 8 MAF and LD bins using either the ratio of averages over loci or the average over loci of ratios methods to compute the GRMs (Supplementary Figure 22). To investigate the relationship between LD and effect size, we also divided each MAF bin into 3 (low, medium and high LD) or 4 LD bins (quartiles) with a similar number of variants and calculated the GRMs using the ratio of averages method. We ran a GREML-LDMS analysis using the 12 or 16 GRMs (Figure 2, Supplementary Figure 26). For each model, we calculated the corresponding AIC using the log-likelihood and the number of fixed and random effects fitted in the model (Supplementary Figure 27).

### Enrichment analysis using the variant effect consequence

Using SnpEff annotations^26^ and the LD and MAF bins defined from the GREML-LDMS analysis on the WGS data mentioned above, we further separated the low-LD variants in each of the 0.0001<MAF<0.001, 0.001<MAF<0.01 and 0.01<MAF<0.1 bins into 2 bins according to their predicted variant effects. The SnpEff variant effect annotations were divided into 4 categories according to their predicted effects on gene expression and protein translation. The 4 categories are based on the Sequence Ontology terms used in functional annotations (Supplementary Table 5). Putative effects on proteins can be “High” (protein truncating variants, frameshift variants, stop gained, and stop lost etc), “Moderate” (mostly non-synonymous variants), “Low” (mostly synonymous variants) or “Modifier” (mostly intronic and upstream or downstream regulatory variants). We merged variants having “High” and “Moderate” impacts in a “Protein-altering” bin and variants having “Low” and “Modifier” impacts in a “Non-protein-altering” bin. We then ran a GREML-LDMS analysis with 11 GRMs, fitting the 48 PCs shown to well capture the effect of population stratification (Figure 3) as fixed covariates. To compute the variance explained per SNP, we divided the estimate of variance explained for each bin by the number of variants in the bin. The standard error was obtained by dividing the standard error of the estimated variance explained for the bin by the number of variants in the bin.

### Investigation of the influence of spatial coordinates on GREML estimates using whole-exome sequence (WES) data from the UKB

We used the spatial coordinates in the UKB to evaluate the influence of local spatial stratification on GREML estimates. Based on a GRM derived from HM3 SNPs in the UKB, we selected a set of 35,867 unrelated individuals of European ancestry with WES data available. These individuals also had phenotypical data, including age, sex, height, BMI, assessment center and north / east birth coordinates. For each sex, we regressed height and BMI against age and then standardized the residuals to N(0,1). We also performed an RINT for both height and BMI. We imputed the missing north and east birth coordinates (UKB data field 129 and 130) using the average birth coordinates of the samples from the same assessment centers and scaled each coordinate to fall into 0 to 1 range. Quality control on the genotype data were excluding variants with genotypes missingness rate >0.05, Hardy-Weinberg equilibrium test *P* value <1 × 10^-6^, or with a minor allele count > 3 (similar MAF than the TOPMed dataset) and excluding individuals with sample missingness rate >0.05. We had 2,075,174 variants left in the dataset. We removed HM3 SNPs duplicated with the imputed dataset from SNP array and calculated 14 GRMs based on individual LD and MAF properties of the variants. We ran a GREML-LDMS fitting 14 GRMs from exome variants and a GRM from HM3 SNPs imputed from SNP array. We fitted as fixed covariates 20 PCs computed from HM3 variants. To evaluate the effect of local spatial stratification, we also fitted the east and north birth coordinates on top of the PCs.

We compared our estimates with TOPMed by selecting variants found in both the TOPMed WGS and UKB WES datasets. We kept ~678k variants observed in both datasets, filtered out the ones showing a deviation from Hardy-Weinberg with a Fisher’s exact test p-value < 0.05 (Bonferroni corrected) and then grouped them into ~130K common ones (MAF > 0.01) and ~548k rare ones (0.0001 < MAF < 0.01). We calculated 4 GRMs (MAF- and LD-stratified) for each of the TOPMed and UKB Exome datasets and ran a GREML-LDMS analysis fitting 20 PCs calculated from their respective HM3 SNPs.

### Rare variants population stratification

To evaluate the effect of population stratification, we separated the UKB Exome dataset into whether a variant was on odd or even chromosomes. For each of these two sets of variants, we stratified them by MAF and LD into 14 bins, pruned them for LD in each bin using different criteria (window size and LD r^2^ threshold of 50 and 0.1 respectively for common variants and of 2000 and 0.01 respectively, for rare variants), and used the pruned variants to compute 150 PCs in each bin. We then evaluated the population stratification by looking at the adjusted R^2^ of fitting each PC from a set of chromosomes against all the PCs from the other chromosomes in each MAF/LD bin. For each PC, by taking the mean of the two R^2^ computed on a chromosome set on the other and vice versa, we obtained an average R^2^ of inter-chromosomal correlations for each MAF/LD bin (Supplementary Figure 17). For each individual, we computed across all MAF/LD bins, 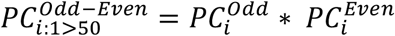 as the product of the PCs of odd and even chromosomes. After centering and scaling, we applied the RINT transformation to smoothen outliers and plotted this PC interaction term for each individual according to their birth coordinates (Supplementary Figure 18). With this PC interaction term, we computed Moran’s I as a measure of spatial autocorrelation (Supplementary Figure 19). We repeated the same procedure (computing PCs using LD-pruned SNPs from odd and even chromosomes) for the TOPMed individuals (Supplementary Figure 17). For the adjusted R^2^ that measures inter-chromosomal correlations, we used a segmented regression to find the optimal number of PCs to be fitted in the GREML-LDMS analysis to account for population stratification.

### Influence of outlier samples on heritability estimates

To investigate the influence of outlier samples/pairs on the heritability estimates, we investigated ways to filter outlier individuals other than the QC step on heterozygosity. We also noticed a strong relationship between the proportion of African ancestry and the diagonal values of the GRM constructed from variants in the MAF range of 0.0001 to 0.001 in the high-LD bins (Supplementary Notes). To investigate the influence of extreme values across the GRMs, we removed extreme values from the GRMs computed on unrelated individuals of European ancestry (N=28,755). We removed samples based either on the GRM diagonals, off-diagonals or both. For the filtering based on diagonals, we removed samples with diagonal values smaller than 0.7 or larger than 1.3 across any of the 8 WGS GRMs. This step removed 3,426 individuals, with the extreme values mostly coming from the rare variants GRMs in the high-LD groups. For the filtering based on off-diagonals, we selected a value of 0.1 across all GRMs as a cut-off to remove one of each pair of individuals with large off-diagonal values. It is of note that the pairs with large off-diagonal values were all found in the rare variants GRMs in the high-LD bins, which is partly because we have pruned individuals for relatedness based on the GRM derived from HM3 SNPs. This process removed additional 2,061 individuals. Finally, we also filtered samples based on both their diagonal and off-diagonals (pair > 0.1) GRM values (4,526 samples removed in total).

## Acknowledgements

This research was supported by the Australian Research Council (DP160102400, FT180100186, FL180100072 and DE200100425), the Australian National Health and Medical Research Council (1113400 and 1078037), the US National Institutes of Health (R01MH100141), the Sylvia & Charles Viertel Charitable Foundation, and the Westlake Education Foundation. This study makes use of data from UK10K project (a full list of acknowledgements to this dataset can be found below).

Whole genome sequencing (WGS) for the Trans-Omics in Precision Medicine (TOPMed) program was supported by the National Heart, Lung and Blood Institute (NHLBI). WGS for “NHLBI TOPMed: Genetics of Cardiometabolic Health in the Amish” (phs000956.v3.p1.c999) was performed at the Broad Institute of MIT and Harvard (HHSN268201500014C). WGS for TOPMed “NHLBI TOPMed: Trans-Omics for Precision Medicine Whole Genome Sequencing Project: ARIC” (phs001211.v1.p1.c999) was performed at the Broad Institute of MIT and at the Baylor Human Genome Sequencing Center (3R01HL092577-06S1, HHSN268201500015C, 3U54HG003273-12S). WGS for TOPMed “NHLBI TOPMed: Mount Sinai BioMe Biobank (BioMe)” (phs001644.v1.p1.c999) was performed at the McDonnell Genome Institute and at the Baylor Human Genome Sequencing Center (HHSN268201600037I, HHSN268201600033I).WGS for TOPMed “NHLBI TOPMed: Coronary Artery Risk Development in Young Adults (CARDIA)” (phs001612.v1.p1.c999) was performed at the Baylor Human Genome Sequencing Center and at the Keck Molecular Genomics Core Facility (HHSN268201600038I, HHSN268201600033I). WGS for “NHLBI TOPMed: The Cleveland Clinic Atrial Fibrillation Study of the CV/Arrhythmia Biobank” (phs001189.v1.p1.c999) was performed at the Broad Institute of MIT and Harvard (3R01HL092577-06S1). WGS for “NHLBI TOPMed: The Cleveland Family Study (WGS)” (phs000954.v2.p1.c999) was performed at the University of Washington Northwest Genomics Center (3R01HL098433-05S1). WGS for “NHLBI TOPMed: Cardiovascular Health Study” (phs001368.v1.p1.c999) was performed at the Baylor Human Genome Sequencing Center (HHSN268201500015C, 75N92021D00006). WGS for “NHLBI TOPMed: Genetic Epidemiology of COPD (COPDGene) in the TOPMed Program” (phs000951.v2.p2.c999) was performed at the Broad Institute of MIT and Harvard and the University of Washington Northwest Genomics Center (HHSN268201500014C). WGS for “NHLBI TOPMed: Whole Genome Sequencing and Related Phenotypes in the Framingham Heart Study” (phs000974.v3.p2.c999) was performed at the Broad Institute of MIT and Harvard (3R01HL092577-06S1). WGS for “NHLBI TOPMed: GeneSTAR (Genetic Study of Atherosclerosis Risk)” (phs001218.v1.p1.c999) was performed at the Broad Institute of MIT and Harvard, at Macrogen Corp (HHSN268201500014C) and at Illumina (HL112064).WGS for “NHLBI TOPMed: Genetics of Lipid Lowering Drugs and Diet Network (GOLDN)” (phs001359.v1.p1.c999) was performed at the University of Washington Northwest Genomics Center (3R01HL104135-04S1). WGS for “NHLBI TOPMed: Heart and Vascular Health Study (HVH)” (phs000993.v2.p2.c999) was performed at the Broad Institute of MIT and Harvard and the Baylor Human Genome Sequencing Center (3R01HL092577-06S1, 3U54HG003273-12S2). WGS for “NHLBI TOPMed: Whole Genome Sequencing of Venous Thromboembolism (WGS of VTE)” (phs001402.v1.p1.c999) was performed at the Baylor Human Genome Sequencing Center (HHSN268201500015C, 3U54HG003273-12S2). WGS for “NHLBI TOPMed: MESA and MESA Family AA-CAC” (phs001416.v1.p1.c999) was performed at the Broad Institute of MIT and Harvard (3U54HG003067-13S1, HHSN268201500014C). WGS for “NHLBI TOPMed: MGH Atrial Fibrillation Study” (phs001062.v3.p2.c999) was performed at the Broad Institute of MIT and Harvard (3R01HL092577-06S1). WGS for “NHLBI TOPMed: Partners Healthcare Biorepository (Partners)” (phs001024.v1.p1.c999) was performed at the Broad Institute of MIT and Harvard (3R01HL092577-06S1). WGS for “NHLBI TOPMed: The Vanderbilt AF Ablation Registry” (phs000997.v3.p2.c999) was performed at the Broad Institute of MIT and Harvard (3R01HL092577-06S1). WGS for “NHLBI TOPMed: The Vanderbilt Atrial Fibrillation Registry” (phs001032.v3.p2.c999) was performed at the Broad Institute of MIT and Harvard (3R01HL092577-06S1). WGS for “NHLBI TOPMed: The Women’s Genome Health Study” (phs001040.v3.p1.c999) was performed at the Broad Institute of MIT and Harvard (3R01HL092577-06S1). WGS for “NHLBI TOPMed: Women’s Health Initiative (WHI)” (phs001237.v1.p1.c999) was performed at the Broad Institute of MIT and Harvard (HHSN268201500014C). Centralized read mapping and genotype calling, along with variant quality metrics and filtering were provided by the TOPMed Informatics Research Center (3R01HL-117626-02S1). Phenotype harmonization, data management, sample-identity QC, and general study coordination, were provided by the TOPMed Data Coordinating Center (3R01HL-120393-02S1). We gratefully acknowledge the studies and participants who provided biological samples and data for TOPMed.

The full TOPMed Consortium authorship list can be found at: https://www.nhlbiwgs.org/topmed-banner-authorship

## COHORT SPECIFIC ACKNOWLEDGEMENTS

### Amish Research Program

This research has been supported in part by NIH grants U01 HL072515, U01 HL072515, R01 AG18728, and P30 DK072488.

### Atherosclerosis Risk in Communities Study

The Atherosclerosis Risk in Communities study has been funded in whole or in part with Federal funds from the National Heart, Lung, and Blood Institute, National Institutes of Health, Department of Health and Human Services, under Contract nos. (HHSN268201700001I, HHSN268201700002I, HHSN268201700003I, HHSN268201700005I, HHSN268201700004I). The authors thank the staff and participants of the ARIC study for their important contributions.

### Coronary Artery Risk Development in Young Adults

Molecular data for the Trans-Omics in Precision Medicine (TOPMed) program was supported by the National Heart, Lung and Blood Institute (NHLBI). Whole Genome Sequencing for the NHLBI TOPMed: CARDIA Study (phs001612) was performed at the Baylor College of Medicine Human genome Sequencing Center (contract HHSN268201600033I). Core support including centralized genomic read mapping and genotype calling, along with variant quality metrics and filtering were provided by the TOPMed Informatics Research Center (3R01HL-117626-02S1; contract HHSN268201800002I). Core support including phenotype harmonization, data management, sample-identity QC, and general program coordination were provided by the TOPMed Data Coordinating Center (R01HL-120393; U01HL-120393; contract HHSN268201800001I). We gratefully acknowledge the studies and participants who provided biological samples and data for TOPMed.

The Coronary Artery Risk Development in Young Adults Study (CARDIA) is conducted and supported by the National Heart, Lung, and Blood Institute (NHLBI) in collaboration with the University of Alabama at Birmingham (HHSN268201800005I & HHSN268201800007I), Northwestern University (HHSN268201800003I), University of Minnesota (HHSN268201800006I), and Kaiser Foundation Research Institute (HHSN268201800004I).

### Cardiovascular Health Study

This research was supported by contracts HHSN268201200036C, HHSN268200800007C, HHSN268201800001C, N01HC55222, N01HC85079, N01HC85080, N01HC85081, N01HC85082, N01HC85083, N01HC85086, and grants U01HL080295, U01HL130114, and R01HL059367 from the National Heart, Lung, and Blood Institute (NHLBI), with additional contribution from the National Institute of Neurological Disorders and Stroke (NINDS). Additional support was provided by R01AG023629 from the National Institute on Aging (NIA). A full list of principal CHS investigators and institutions can be found at CHS-NHLBI.org. The content is solely the responsibility of the authors and does not necessarily represent the official views of the National Institutes of Health.

### Cleveland Family Study (CFS)

CFS was supported by National Institutes of Health grants National Institutes of Health grants R01-HL046380-15, HL113338, and KL2-RR024990-05, and R35HL135818 [5-R01-HL046380, and 5-KL2-RR024990-05.

### Genetic Epidemiology of COPD Study (COPDGene)

The COPDGene project described was supported by Award Number U01 HL089897 and Award Number U01 HL089856 from the National Heart, Lung, and Blood Institute. The content is solely the responsibility of the authors and does not necessarily represent the official views of the National Heart, Lung, and Blood Institute or the National Institutes of Health. The COPDGene project is also supported by the COPD Foundation through contributions made to an Industry Advisory Board comprised of AstraZeneca, Boehringer Ingelheim, GlaxoSmithKline, Novartis, Pfizer, Siemens and Sunovion. A full listing of COPDGene investigators can be found at: http://www.copdgene.org/directory

### Framingham Heart Study

The Framingham Heart Study (FHS) acknowledges the support of contracts NO1-HC-25195 and HHSN268201500001I from the National Heart, Lung and Blood Institute and grant supplement R01 HL092577-06S1 for this research. We also acknowledge the dedication of the FHS study participants without whom this research would not be possible. Dr. Vasan is supported in part by the Evans Medical Foundation and the Jay and Louis Coffman Endowment from the Department of Medicine, Boston University School of Medicine.

### Genetics of Lipid Lowering Drugs and Diet Network (GOLDN)

GOLDN biospecimens, baseline phenotype data, and intervention phenotype data were collected with funding from National Heart, Lung and Blood Institute (NHLBI) grant U01 HL072524. Whole-genome sequencing in GOLDN was funded by NHLBI grant R01 HL104135 and supplement R01 HL104135-04S1.

### Massachusetts General Atrial Fibrillation Study

This work was supported by grants from the National Institutes of Health to Dr. Ellinor (1RO1HL092577, R01HL128914, K24HL105780). Dr. Ellinor is also supported by the American Heart Association (18SFRN34110082) and by the Fondation Leducq (14CVD01).

### Mayo Clinic Venous Thromboembolism Study (Mayo_VTE)

Funded, in part, by grants from the National Institutes of Health, National Heart, Lung and Blood Institute (HL66216 and HL83141). the National Human Genome Research Institute (HG04735, HG06379), and research support provided by Mayo Foundation.

### Multi-Ethnic Study of Atherosclerosis (MESA)

The MESA project is supported by the National Heart, Lung, and Blood Institute (NHLBI) in collaboration with MESA investigators. Support for MESA is provided by contracts 75N92020D00001, HHSN268201500003I, N01-HC-95159, 75N92020D00005, N01-HC-95160, 75N92020D00002, N01-HC-95161, 75N92020D00003, N01-HC-95162, 75N92020D00006, N01-HC-95163, 75N92020D00004, N01-HC-95164, 75N92020D00007, N01-HC-95165, N01-HC-95166, N01-HC-95167, N01-HC-95168, N01-HC-95169, UL1-TR-000040, UL1-TR-001079, and UL1-TR-001420. Also supported in part by the National Center for Advancing Translational Sciences, CTSI grant UL1TR001881, and the National Institute of Diabetes and Digestive and Kidney Disease Diabetes Research Center (DRC) grant DK063491 to the Southern California Diabetes Endocrinology Research Center.

### The Partners Healthcare Biorepository (Partners)

Dr. Lubitz is supported by NIH grant 1R01HL139731 and American Heart Association 18SFRN34250007.

### The Johns Hopkins Genetic Study of Atherosclerosis Risk (GeneSTAR)

was supported by grants from the National Institutes of Health through the National Heart, Lung, and Blood Institute (HL49762, HL071025, U01HL72518, HL087698, HL092165, HL099747, K23HL105897, HL112064) and the National Institute of Nursing Research (NR0224103), by a grant from the National Center for Research Resources (M01-RR000052) to the Johns Hopkins General Clinical Research Center, and by a grant from the National Center for Research Resources and the National Center for Advancing Translational Sciences (UL1 RR 025005) to the Johns Hopkins Institute for Clinical and Translational Research.

### The Mount Sinai BioMe Biobank (BioME)

is supported by The Andrea and Charles Bronfman. We thank the participants in the BioMe Biobank for their invaluable contribution to biomedical research.

Ruth Loos is supported by the NIH (X01HL134588; R01DK110113; R01HL142302; R01DK107786; R01DK124097).

### The Women’s Health Initiative (WHI)

The WHI program is funded by the National Heart, Lung, and Blood Institute, National Institutes of Health, U.S. Department of Health and Human Services through contracts HHSN268201600018C, HHSN268201600001C, HHSN268201600002C, HHSN268201600003C, and HHSN268201600004C. <u>Investigators: *Program Office*</u>: (National Heart, Lung, and Blood Institute, Bethesda, Maryland) Jacques Rossouw, Shari Ludlam, Joan McGowan, Leslie Ford, and Nancy Geller. <u>*Clinical Coordinating Center*</u>: (Fred Hutchinson Cancer Research Center, Seattle, WA) Garnet Anderson, Ross Prentice, Andrea LaCroix, and Charles Kooperberg. <u>*Investigators and Academic Centers*</u>: (Brigham and Women’s Hospital, Harvard Medical School, Boston, MA) JoAnn E. Manson; (MedStar Health Research Institute/Howard University, Washington, DC) Barbara V. Howard; (Stanford Prevention Research Center, Stanford, CA) Marcia L. Stefanick; (The Ohio State University, Columbus, OH) Rebecca Jackson; (University of Arizona, Tucson/Phoenix, AZ) Cynthia A. Thomson; (University at Buffalo, Buffalo, NY) Jean Wactawski-Wende; (University of Florida, Gainesville/Jacksonville, FL) Marian Limacher; (University of Iowa, Iowa City/Davenport, IA) Jennifer Robinson; (University of Pittsburgh, Pittsburgh, PA) Lewis Kuller; (Wake Forest University School of Medicine, Winston-Salem, NC) Sally Shumaker; (University of Nevada, Reno, NV) Robert Brunner. <u>*Women’s Health Initiative Memory Study*</u>: (Wake Forest University School of Medicine, Winston-Salem, NC) Mark Espeland

<u>UK10K</u> (EGA accessions: EGAS00001000108 and EGAS00001000090): was funded by the Wellcome Trust award WT091310. Twins UK (TUK): TUK was funded by the Wellcome Trust and ENGAGE project grant agreement HEALTH-F4-2007-201413. The study also receives support from the Department of Health via the National Institute for Health Research (NIHR)-funded BioResource, Clinical Research Facility and Biomedical Research Centre based at Guy’s and St. Thomas’ NHS Foundation Trust in partnership with King’s College London. Dr Spector is an NIHR senior Investigator and ERC Senior Researcher. Funding for the project was also provided by the British Heart Foundation grant PG/12/38/29615 (Dr Jamshidi). A full list of the investigators who contributed to the UK10K sequencing is available from https://www.uk10k.org/

### UK Biobank

This research has been conducted using the UK Biobank Resource under project 12505.

## References

1 Fisher, R. A. XV.—The Correlation between Relatives on the Supposition of Mendelian Inheritance. Transactions of the Royal Society of Edinburgh 52, 399–433, doi:10.1017/s0080456800012163 (1918).

2 Yang, J. et al. Common SNPs explain a large proportion of the heritability for human height. Nat Genet 42, 565–569, doi:10.1038/ng.608 (2010).

3 Visscher, P. M., Brown, M. A., McCarthy, M. I. & Yang, J. Five years of GWAS discovery. Am J Hum Genet 90, 7–24, doi:10.1016/j.ajhg.2011.11.029 (2012).

4 Finucane, H. K. et al. Partitioning heritability by functional annotation using genomewide association summary statistics. Nat Genet 47, 1228–1235, doi:10.1038/ng.3404 (2015).

5 Speed, D. et al. Reevaluation of SNP heritability in complex human traits. Nat Genet 49, 986–992, doi:10.1038/ng.3865 (2017).

6 Lynch, M. & Walsh, B. Genetics and analysis of quantitative traits. (Sinauer, 1998).

7 MacArthur, J. et al. The new NHGRI-EBI Catalog of published genome-wide association studies (GWAS Catalog). Nucleic Acids Res 45, D896–D901, doi:10.1093/nar/gkw1133 (2017).

8 Gazal, S. et al. Linkage disequilibrium-dependent architecture of human complex traits shows action of negative selection. Nat Genet 49, 1421–1427, doi:10.1038/ng.3954 (2017).

9 Zeng, J. et al. Signatures of negative selection in the genetic architecture of human complex traits. Nat Genet 50, 746–753, doi:10.1038/s41588-018-0101-4 (2018).

10 Yang, J. et al. Genetic variance estimation with imputed variants finds negligible missing heritability for human height and body mass index. Nat Genet 47, 1114–1120, doi:10.1038/ng.3390 (2015).

11 Zuk, O., Hechter, E., Sunyaev, S. R. & Lander, E. S. The mystery of missing heritability: Genetic interactions create phantom heritability. Proc Natl Acad Sci U S A 109, 1193–1198, doi:10.1073/pnas.1119675109 (2012).

12 Young, A. I. et al. Relatedness disequilibrium regression estimates heritability without environmental bias. Nat Genet 50, 1304–1310, doi:10.1038/s41588-018-0178-9 (2018).

13 Genomes Project, C. et al. A global reference for human genetic variation. Nature 526, 68–74, doi:10.1038/nature15393 (2015).

14 Bergstrom, A. et al. Insights into human genetic variation and population history from 929 diverse genomes. Science 367, doi:10.1126/science.aay5012 (2020).

15 Yengo, L. et al. Meta-analysis of genome-wide association studies for height and body mass index in approximately 700000 individuals of European ancestry. Hum Mol Genet, doi:10.1093/hmg/ddy271 (2018).

16 International HapMap, C. et al. Integrating common and rare genetic variation in diverse human populations. Nature 467, 52–58, doi:10.1038/nature09298 (2010).

17 Yang, J., Lee, S. H., Goddard, M. E. & Visscher, P. M. GCTA: a tool for genome-wide complex trait analysis. Am J Hum Genet 88, 76–82, doi:10.1016/j.ajhg.2010.11.011 (2011).

18 Yang, J. et al. Genome partitioning of genetic variation for complex traits using common SNPs. Nat Genet 43, 519–525, doi:10.1038/ng.823 (2011).

19 McCarthy, S. et al. A reference panel of 64,976 haplotypes for genotype imputation. Nat Genet 48, 1279–1283, doi:10.1038/ng.3643 (2016).

20 Evans, L. M. et al. Comparison of methods that use whole genome data to estimate the heritability and genetic architecture of complex traits. Nat Genet 50, 737–745, doi:10.1038/s41588-018-0108-x (2018).

21 Elks, C. E. et al. Variability in the heritability of body mass index: a systematic review and meta-regression. Frontiers in endocrinology 3, 29, doi:10.3389/fendo.2012.00029 (2012).

22 Mitt, M. et al. Improved imputation accuracy of rare and low-frequency variants using population-specific high-coverage WGS-based imputation reference panel. Eur J Hum Genet 25, 869–876, doi:10.1038/ejhg.2017.51 (2017).

23 Mathieson, I. & McVean, G. Differential confounding of rare and common variants in spatially structured populations. Nat Genet 44, 243–246, doi:10.1038/ng.1074 (2012).

24 Goudet, J., Kay, T. & Weir, B. S. How to estimate kinship. Mol Ecol 27, 4121–4135, doi:10.1111/mec.14833 (2018).

25 UK10K Consortium. The UK10K project identifies rare variants in health and disease. Nature 526, 82–90, doi:10.1038/nature14962 (2015).

26 Cingolani, P. et al. A program for annotating and predicting the effects of single nucleotide polymorphisms, SnpEff: SNPs in the genome of Drosophila melanogaster strain w1118; iso-2; iso-3. Fly (Austin) 6, 80–92, doi:10.4161/fly.19695 (2012).

27 Gusev, A. et al. Partitioning heritability of regulatory and cell-type-specific variants across 11 common diseases. Am J Hum Genet 95, 535–552, doi:10.1016/j.ajhg.2014.10.004 (2014).

28 Keinan, A. & Clark, A. G. Recent explosive human population growth has resulted in an excess of rare genetic variants. Science 336, 740–743, doi:10.1126/science.1217283 (2012).

29 Genome of the Netherlands, C. Whole-genome sequence variation, population structure and demographic history of the Dutch population. Nat Genet 46, 818–825, doi:10.1038/ng.3021 (2014).

30 Stulp, G., Simons, M. J., Grasman, S. & Pollet, T. V. Assortative mating for human height: A meta-analysis. Am J Hum Biol 29, doi:10.1002/ajhb.22917 (2017).

31 Border, R. et al. Assortative Mating Biases Marker-based Heritability Estimators. BioRvix, doi:10.1101/2021.03.18.436091 (2021).

32 Visscher, P. M. et al. Assumption-free estimation of heritability from genome-wide identity-by-descent sharing between full siblings. PLoS Genet 2, e41, doi:10.1371/journal.pgen.0020041 (2006).

33 Kemper, K. E. et al. Phenotypic covariance across the entire spectrum of relatedness for 86 billion pairs of individuals. Nat Commun 12, 1050, doi:10.1038/s41467-021-21283-4 (2021).

34 Hernandez, R. D. et al. Ultra-rare variants drive substantial cis-heritability of human gene expression. BioRvix, doi:10.1101/219238 (2019).

35 Visscher, P. M. et al. Statistical power to detect genetic (co)variance of complex traits using SNP data in unrelated samples. PLoS Genet 10, e1004269, doi:10.1371/journal.pgen.1004269 (2014).

36 Shihab, H. A. et al. An integrative approach to predicting the functional effects of noncoding and coding sequence variation. Bioinformatics 31, 1536–1543, doi:10.1093/bioinformatics/btv009 (2015).

37 Yengo, L. et al. Imprint of assortative mating on the human genome. Nature Human Behaviour 2, 948–954, doi:10.1038/s41562-018-0476-3 (2018).

38 Uricchio, L. H., Zaitlen, N. A., Ye, C. J., Witte, J. S. & Hernandez, R. D. Selection and explosive growth alter genetic architecture and hamper the detection of causal rare variants. Genome Res 26, 863–873, doi:10.1101/gr.202440.115 (2016).

39 Chang, C. C. et al. Second-generation PLINK: rising to the challenge of larger and richer datasets. Gigascience 4, 7, doi:10.1186/s13742-015-0047-8 (2015).

40 Jiang, L. et al. A resource-efficient tool for mixed model association analysis of large-scale data. Nat Genet 51, 1749–1755, doi:10.1038/s41588-019-0530-8 (2019).

41 VanRaden, P. M. Efficient methods to compute genomic predictions. J Dairy Sci 91, 4414–4423, doi:10.3168/jds.2007-0980 (2008).

42 Loh, P. R. et al. Reference-based phasing using the Haplotype Reference Consortium panel. Nat Genet 48, 1443–1448, doi:10.1038/ng.3679 (2016).

43 Hill, W. G. & Weir, B. S. Variation in actual relationship as a consequence of Mendelian sampling and linkage. Genet Res (Camb) 93, 47–64, doi:10.1017/S0016672310000480 (2011).

44 Hill, W. G. & White, I. M. Identification of pedigree relationship from genome sharing. G3 (Bethesda) 3, 1553–1571, doi:10.1534/g3.113.007500 (2013).

45 Manichaikul, A. et al. Robust relationship inference in genome-wide association studies. Bioinformatics 26, 2867–2873, doi:10.1093/bioinformatics/btq559 (2010).

46 Browning, S. R. & Browning, B. L. High-resolution detection of identity by descent in unrelated individuals. Am J Hum Genet 86, 526–539, doi:10.1016/j.ajhg.2010.02.021 (2010).

47 Palamara, P. F., Lencz, T., Darvasi, A. & Pe’er, I. Length distributions of identity by descent reveal fine-scale demographic history. Am J Hum Genet 91, 809–822, doi:10.1016/j.ajhg.2012.08.030 (2012).

48 Nait Saada, J. et al. Identity-by-descent detection across 487,409 British samples reveals fine scale population structure and ultra-rare variant associations. Nat Commun 11, 6130, doi:10.1038/s41467-020-19588-x (2020).

49 Hinrichs, A. S. et al. The UCSC Genome Browser Database: update 2006. Nucleic Acids Res 34, D590–598, doi:10.1093/nar/gkj144 (2006).

