## Supplementary materials for "Recovery of trait heritability from whole genome sequence data"

### Supplementary note 1: the effect of rare variants population stratification on genomic relationship matrices

Here, we describe various analyses aiming at better understanding the impact of rare variants stratification on estimates of SNP-based heritability from WGS data. The following notes expand in greater details some of the analyses briefly described in the Online Methods section.

#### Extreme values in rare-variants GRMs

After performing an initial QC to select unrelated adults (of age > 18) samples of European ancestry (N = 28,755) based on the first 4 PCs for common and rare variants and 1000 Genomes reference populations[13], we performed QC on variants using several filters (excluding variants with genotypes missingness rate > 0.05, Hardy-Weinberg equilibrium test $P$ value $<1 \times 10^{-6}$, MAF < 0.0001 and high quality variants from a SVM classifier). On the 36.9M variants left after sample and genotype QC, defined 8 variants bins based on their MAF (0.0001 – 0.001, 0.001 – 0.01, 0.01 – 0.1, 0.1 – 0.5) and individual LD score value (median-based). We then computed GRM using these 8 variants groupings. We observed extreme diagonal and off-diagonals values in GRMs (Online methods) calculated from variants with 0.0001 < MAF < 0.01. Prior to performing any further sample/variant filtering, we first investigated the influence of the GRM estimator on these extreme values. We compared two GRM estimators implemented in GCTA: the default GRM estimator[17], which calculate average of per-SNP relatedness estimates; and the VanRaden estimator,[41] which estimates relatedness as the ratio of SNP covariance and SNP heterozygosity. We found small differences between the diagonal elements of these two matrices (Supplementary Figure 29) and larger differences in off-diagonal elements of the GRM (Supplementary Figure 30). The average of ratios estimator shows off-diagonal values consistently higher for extreme values. As a whole, we observe, for variants with 0.0001 < MAF < 0.01, high relatedness for some pairs (with relatedness values consistent with a first-degree relationship) and large GRM diagonal values that are inconsistent with the expected sampling variance of rare variants GRMs with a larger number of effective markers[10]. The GRM estimator alone does not explain why a set of unrelated (estimated common variants relatedness < 0.05) samples using HM3 common SNPs exhibits very high relatedness from rare variants GRMs.

#### Influence of IBD segments on relatedness

The relatedness threshold of 0.05 using HM3 common SNPs, while being widely used in previous studies[10], might not be sufficient to remove residual relatedness. For example, cousins-once-removed have an expected relatedness value of 0.0625 (SD 0.017)[43]. A relatedness threshold of 0.05 could then include such pairs who would share segments IBD as shown through simulations in previous studies[44]. We first selected samples with a more stringent threshold on HM3 SNPs of 0.025. That removed an additional 2282 samples. When investigating the rare variants GRMs we could still observe extreme values in the diagonals (~4) and off-diagonals (~1) elements, despite the more stringent threshold on common variants relatedness.

In the absence of sample relatedness for common variants and genome-wide population stratification, large values in rare variants GRMs could be driven by shared IBD segments. To identify the shared IBD segments, we selected 987,393 LD-pruned variants from the full WGS data (window size of 50kb and LD $r^2$ threshold of 0.1) with MAF > 0.01 and identified pairs shared IBD segments > 1MB on each of the 22 chromosomes with KING software[45].

Across all chromosomes, 56.6M segments were shared IBD with 3.26M unique pairs sharing one or more segments IBD (Supplementary Figure 31). For each pair of individuals (sharing at least 1 segment IBD), we compared the cumulative length of IBD segments with its corresponding GRM off-diagonal values. We found that some pairs sharing the longest IBD segments accumulatively did not contribute disproportionately to the GRM off-diagonal elements for high-LD variants with 0.0001 < MAF < 0.01. We also noticed that some pairs had an extreme relatedness value (large off-diagonals), yet did not share a large proportion of their genome IBD. These pairs were enriched with individuals exhibiting a high level of MAF/LD bin-specific heterozygosity (Supplementary Figure 32). These results suggest that high off-diagonal values are caused by high mean heterozygosity due to variants that are disproportionally shared but not in long IBD segments.

We confirmed these findings by investigating, for a pair of individuals showing an extreme off-diagonal value in the GRM for high-LD variants with 0.0001 < MAF < 0.001, the distribution of the heterozygous rare and oppositive homozygous common variants for the chromosome enriched for rare variants shared by the pair (Supplementary Figure 33). For this specific pair, we identified a region with many clumped rare variants and no oppositive homozygotes common variants, suggesting a region shared IBD.

We then defined 3 groups of samples (Supplementary Figure 34) selecting either the first 100 pairs with the highest off-diagonals (for high-LD variants with 0.0001 < MAF < 0.001), the first 100 pairs sharing the longest proportion of their genome IBD, and 100 pairs selected at random (within 0.01 SD from the median of IBD length shared and off-diagonal values) as control. For each group, we used a local ancestry reference to infer the genome-wide proportion ancestry of the group of samples (Online methods). There was no difference in mean ancestry across sample groups for each population with the exception that samples with high GRM off-diagonals had a significantly higher proportion of African ancestry, indicating that these pairs might be sharing few shorts IBD segments from a different ancestry, thereby strongly impacting their relatedness. We further investigated the source of heterozygosity by comparing the number of heterozygous variants from either the IBD segments or the entire genome, for the very rare high-LD variants (Supplementary Figure 34). The group showing high relatedness had a very high number of heterozygotes variants (as expected by their high mean heterozygosity rate) with most of these variants on their shared IBD segments. The group sharing a higher proportion of genome IBD had a substantially lower number of heterozygotes variants. Finally, the control group showed a small number of heterozygous variants overall and more spread across the entire genome.

In summary, we show that short IBD segments from a different ancestry have a strong impact on the estimated relatedness values for some pairs in the rare variants GRMs. Including a step in the QC pipeline to identify and remove such pairs allows the correction for extreme relatedness which could bias the variance components estimates. While directly investigating the origin of these segments specifically (through segment-based PCA, for example) is computationally demanding, we used the sample heterozygosity to identify the pairs sharing such segments and remove them. We used multiple rounds of heterozygosity QC to remove the extreme samples (Online methods). Shared IBD segments are expected in an outbred population[46-48], but we tried to control for the large effects it can have when considering very rare variants.

### Residual population stratification

Finally, we further our investigation to the influence of ancestry proportion on the diagonal elements of the GRMs. Our analysis is limited to individuals already selected to be from European ancestry by a PCA filtering on both common and rare variants then further filtered to stay within 3 SD from main reference populations using a local ancestry inference (Online methods). When investigating the relationship between the diagonal elements and the proportion of each ancestry for each MAF/LD grouping, we could not notice a meaningful relationship with the exception of the impact of African ancestry proportion positively correlated ($R^2 = 0.69$) with diagonal values for the high-LD variants with $0.0001 < \text{MAF} < 0.001$ (Supplementary Figure 35). The larger genetic diversity for African populations has been documented by previous studies[13] and is reflected in the diagonal values for rare variants GRMs.

The samples in our study have, after QC steps, a maximum of 1.5% of their genome that is estimated to be derived from African ancestry. Lowering this threshold would remove a large number of samples. Moreover, the sample identified as having African ancestry IBD segments still show an average proportion of African ancestry genome-wide of around 0.75%, far below our current threshold. Thus, while it is important to consider removing sample showing an excess of any ancestry other than the population under study (here, people of European ancestry), this criterion alone cannot be used to control for subtle population stratification and needs to be used in conjunction with more specific metrics (such as the sample MAF/LD bin heterozygosity rate).

### Quality control for WGS data

Much is yet to be learned when using WGS data with very rare variants for genetic analysis. While the set of 28,755 unrelated European ancestry samples did not present any specific issue when considering only common variants, including very rare variants in our analysis necessitated additional QC steps. Ensuring a homogeneous ancestry with PCA and local ancestry inference helps removing any cryptic population stratification that could influence the diagonals elements of the rare variants GRMs. Moreover, it is important to identify pairs sharing short IBD segments coming from a different ancestry and having a large influence on GRM elements for rare variants. Removing samples based on their MAF/LD bin-specific heterozygosity helps to correct for potentially biased estimates of variance components from rare variant GRMs. While this does not fully address the issue of samples showing large GRM element values, it is an efficient way to correct for the bias identified while trying to maximise the sample size. A WGS data set from a more homogeneous population might not exhibit such patterns among rare variants.

### Supplementary note 2: Testing the robustness of $h^2_{WGS}$ estimates

The following section extends the analysis performed to ensure the robustness of our heritability estimates for height and BMI. We used a set of N=25,465 unrelated samples of European ancestry without excess of heterozygosity. Variants were removed using several quality filters (genotypes missingness rate, Hardy-Weinberg equilibrium test *P* value, MAF, quality classifier) and grouped into 8 bins according to their MAF (0.0001 – 0.001, 0.001 – 0.01, 0.01 – 0.1, 0.1 – 0.5) and individual LD score value (median-based). The phenotypes were pre-adjusted for age and standardized to a mean of zero and a variance of 1 in each sex and cohort group. See Online Methods for more details on the dataset and QC performed.

#### *Influence of LD definition on variance estimates*

We observed minor differences in heritability estimates when replicating previous studies using only genotypes from SNP mimicking arrays followed by imputation ($h^2_{G+IMP}$). Estimates of $h^2_{G+IMP}$ for height and BMI were in the range of 0.50-0.56 (SE 0.06-0.07) and 0.16-0.21 (SE 0.07) respectively. Previous estimates[10] were 0.56 (SE 0.02) for height and 0.27 (SE 0.02) for BMI, with a set of 44,126 unrelated samples and ~17.6M imputed variants on 1000 Genomes panel with no filtering on imputation quality score, a segment-based LD definition to define LD bins and no LD pruning on the variants used to compute PCs. Moreover, our height variance estimate using segment-based LD definition using Axiom array SNPs prior to imputation was at 0.51 (SE 0.04) compared to 0.56 (SE 0.07) using individual SNP LD score. While a difference in heritability estimates between LD definition was expected from previous study[20], the difference between current and prior LD-based estimates for height could be explained by the PCs from LD pruned variants capturing better potential population stratification, a different imputation panel, a more stringent imputation quality threshold or differences in base population heritabilities. Differences in BMI could also be explained by the aforementioned reasons and the different LD definitions.

#### *Comparing LD / MAF structure using UK10K data*

We also tested if there was any bias due to a specific LD or MAF structure in the TOPMed dataset by using a different sequenced dataset for SNP stratification. We used the UK10K data set [25] and analysed the TOPMed data using the MAF and LD stratification from either TOPMed or UK10K data. We converted the UK10K WGS data [25] to GRCh38 coordinates using LiftMap, a wrapper Python script for LiftOver [49]. There are 3,781 individuals in the UK10K dataset. As with TOPMed, we performed a quality control of the genotypes using PLINK with the following filtering thresholds: individuals with missingness rate < 0.05 and variants with missingness rate < 0.05, Hardy-Weinberg equilibrium test *P* value $<1 \times 10^{-6}$, or minor allele frequency < 0.0001, and retained 3,781 individuals and 42.68M variants. From these variants, we selected 20.6M variants in common with those in the TOPMed dataset. Using the UK10K genotypes of the 20.6M variants, we defined 4 MAF bins (0.0001<MAF<0. 001, 0.001<MAF<0.01, 0.01<MAF<0.1 and 0.1<MAF<0.5) and further split the variants in each MAF bin into 2 LD bins according to their LD scores (calculated using a window size of 10Mb in either direction). We also defined 8 MAF- and LD-stratified variant bins using the TOPMed genotypes of the 20.6M variants. We estimated and partitioned additive genetic variance in the TOPMed based on these 20.6M variants, using either the MAF and LD annotation from the UK10K or TOPMed, fitting 160 PCs computed from LD-pruned WGS variants. The estimates were highly consistent between the two analyses (Supplementary Figure 24). For height, the estimates were 0.54 (SE 0.08) when using the TOPMed annotation and 0.55 (SE 0.07) using SNP stratification from the UK10K, and the corresponding estimates for BMI were 0.27 (SE 0.07 – 0.08) in both cases. The similarity between the estimates from the two references for MAF and LD stratification suggests that our inference from the TOPMed annotation is not biased by using MAF and LD stratification from another data set.

#### *Effect of rank-inverse normal transformation*

Finally, we analysed both height and BMI (both adjusted for age and standardized within each sex and cohort group) with a rank-based inverse normal transformation (RINT). We used the set of N=25,465 unrelated Europeans samples and 33.7M high quality variants. We used GREML-LDMS fitting 8 MAF/LD bins and 48 PCs capturing population stratification. The estimate for the RINT-transformed phenotypes were similar when compared to those from the untransformed trait for height (0.63 (SE 0.09)) but lower for BMI (0.20 (SE 0.10)) (Supplementary Figure 6). Since BMI naturally has a skewed distribution (Supplementary Figure 4), results may be sensitive to the scale of analysis. Past estimates of heritability from pedigree and GWAS designs are not consistent in the scale, with some using the actual scale of measurement and other performing a pre-analysis logarithm or RINT transformation of the data. We performed analyses on the actual and RINT scale and found that the estimates from the RINT-transformed data appeared more sensitive to the model, although the RINT estimates seems to be more consistent in analyses considering a reduced number of rare variants (such as down-sampling or analysing exome data). These differences could be due to sampling variation and would need to be investigated further with a larger sample size.

### Supplementary material

Supplementary Table 1: List of studies included in analyses with corresponding sample sizes comprised of participants with both genotypic and phenotypic information available before quality control

| **Study name and code** | **Sample size** |
|---|---|
| Genetics of Cardiometabolic Health in the Amish (Amish) | 1111 |
| Atherosclerosis Risk in Communities (ARIC) | 8128 |
| Mount Sinai BioMe Biobank (BioMe) | 11193 |
| Coronary Artery Risk Development in Young Adults (CARDIA) | 3087 |
| Cleveland Clinic Atrial Fibrillation (CCAF) Study | 363 |
| Cleveland Family Study (CFS) | 1259 |
| Cardiovascular Health Study (CHS) | 3537 |
| Genetic Epidemiology of COPD (COPDGene) | 10283 |
| The Framingham Heart Study (FHS) | 4166 |
| Genetic Studies of Atherosclerosis Risk (GeneSTAR) | 1763 |
| Genetics of Lipid Lowering Drugs and Diet Network (GOLDN) | 945 |
| Heart and Vascular Health Study (HVH) | 693 |
| Whole Genome Sequencing of Venous Thromboembolism (Mayo_VTE) | 1345 |
| Multi-Ethnic Study of Atherosclerosis (MESA) | 5351 |
| Massachusetts General Hospital Atrial Fibrillation (MGH_AF) | 989 |
| Partners Healthcare Biorepository (Partners) | 128 |
| The Vanderbilt AF Ablation Registry (VAFAR) | 173 |
| The Vanderbilt Atrial Fibrillation Registry (VU_AF) | 1128 |
| Women's Genome Health Study (WGHS) | 117 |
| Women’s Health Initiative (WHI) | 11031 |
| Total | 66790 |

*Supplementary Table 2: Number of SNPs on each of the Illumina HumanCore24, GSA 24 and Affymetrix Axiom arrays before merging with TOPMed dataset (probes), after merging and after preparing the files for imputation*

| Array name | Number of probes on the array | Number of SNPs after merging with TOPMed | Final number of SNPs after preparing files for imputation |
|---|---|---|---|
| Affymetrix Axiom | 784,849 | 742,749 | 735,920 |
| Illumina HumanCore24 | 263,947 | 257,901 | 257,392 |
| Illumina GSA | 648,327 | 523,370 | 520,235 |

*Supplementary Table 3: Imputation statistics after imputing SNPs on Illumina InfiniumCore24, GSA 24 and Affymetrix Axiom arrays using the HRC reference panel on Sanger imputation servers, with SNP number with imputation info score above 0.3 after QC.*

| Array | Imputation info scores | | | | Number of SNPs with $r^2 > 0.3$ after QC |
|---|---|---|---|---|---|
| | Min. | Median | Mean | Max. | |
| Axiom | 0.3 | 0.829 | 0.782 | 1 | 20,393,950 |
| Infinium | 0.3 | 0. 750 | 0. 727 | 1 | 18,993,608 |
| GSA | 0.3 | 0.791 | 0.755 | 1 | 19,822,722 |

*Supplementary Table 4: Number of variants and LD properties (based on individual variant LD score) of each of the four groupings of the TOPMed dataset according to the allele frequency of the variant. The number of SNPs decrease as the MAF increases as rare variants makes up for most of the variants in the dataset.*

| **MAF bin** | Number of SNPs | LD score properties | | |
|---|---|---|---|---|
| | | Mean | Median | SD |
| 0.0001 – 0.001 | 19,583,648 | 27.1 | 13.6 | 46.3 |
| 0.001 – 0.01 | 5,283,043 | 40.7 | 22.5 | 76.7 |
| 0.01 – 0.1 | 3,935,389 | 122.6 | 71.5 | 260.8 |
| 0.1 – 0.5 | 4,902,760 | 193.6 | 136.3 | 241.7 |

*Supplementary Table 5: Putative impacts and enriched GREML-LDMS analysis bin of variant effects as predicted by SnpEff v4.1 annotation software.*

| Enriched GREML-LDMS bin analysis | Putative impact | Sequence Ontology term |
|---|---|---|
| Protein-altering variants | HIGH | chromosome_number_variation |
| | | exon_loss_variant |
| | | frameshift_variant |
| | | rare_amino_acid_variant |
| | | splice_acceptor/donor_variant |
| | | start_lost |
| | | stop_gained/lost |
| | | transcript_ablation |
| | MODERATE | 3_or_5_prime_UTR_truncation & exon_loss |
| | | coding_sequence_variant |
| | | conservative_inframe_insertion/deletion |
| | | disruptive_inframe_insertion/deletion |
| | | missense_variant |
| | | regulatory_region_ablation |
| | | splice_region_variant |
| | | TFBS_ablation |
| Non-protein-altering variants | LOW | 5_prime_UTR_premature_start_codon_gain_variant |
| | | initiator_codon_variant |
| | | splice_region_variant |
| | | start/stop_retained_variant |
| | | synonymous_variant |
| | MODIFIER | 3_or_5_prime_UTR_variant |
| | | coding_sequence_variant |
| | | conserved_intergenic_variant |
| | | conserved_intron_variant |
| | | downstream_gene_variant |
| | | exon_variant |
| | | feature_elongation/truncation |
| | | gene_variant |
| | | intragenic/intergenic_region |
| | | intron_variant |
| | | mature_miRNA_variant |
| | | miRNA |
| | | NMD_transcript_variant |
| | | non_coding_transcript_exon_variant |
| | | non_coding_transcript_variant |
| | | regulatory_region_amplification |
| | | regulatory_region_variant |
| | | TF_binding_site_variant |
| | | TFBS_amplification |
| | | transcript_amplification |
| | | transcript_variant |
| | | upstream_gene_variant |

*Supplementary Table 6: Additional cohort specific information on age, height, BMI, total sample size and proportion of male/females in the study.*

| Study code | Age | | Height | | BMI | | Sample size | Sex proportion | |
|---|---|---|---|---|---|---|---|---|---|
| | Mean | SD | Mean | SD | Mean | SD | | M | F |
| Amish | 50.2 | 16.8 | 165.7 | 9.0 | 26.9 | 4.6 | 1025 | 0.50 | 0.50 |
| ARIC | 54.8 | 5.7 | 168.8 | 9.5 | 27.3 | 5.0 | 3471 | 0.48 | 0.52 |
| CCAF | 54.7 | 9.3 | 179.6 | 10.0 | 30.5 | 6.4 | 328 | 0.80 | 0.20 |
| CFS | 44.1 | 14.8 | 169.8 | 9.7 | 32.2 | 8.3 | 684 | 0.47 | 0.53 |
| CHS | 73.1 | 5.3 | 164.7 | 9.4 | 27.4 | 4.2 | 69 | 0.41 | 0.59 |
| COPDGene | 59.5 | 9.1 | 170.2 | 9.5 | 28.9 | 6.3 | 8640 | 0.54 | 0.46 |
| FHS | 38.5 | 9.8 | 168.7 | 9.5 | 25.9 | 4.9 | 3714 | 0.46 | 0.54 |
| GeneSTAR | 46.5 | 12.3 | 169.4 | 9.7 | 30.3 | 7.0 | 3070 | 0.40 | 0.60 |
| GOLDN | 48.1 | 16.5 | 171.1 | 9.8 | 28.3 | 5.7 | 902 | 0.47 | 0.53 |
| HVH | 55.4 | 5.9 | 177.0 | 10.1 | 33.8 | 9.4 | 66 | 0.65 | 0.35 |
| Mayo_VTE | 55.6 | 16.4 | 171.5 | 10.7 | 31.1 | 7.6 | 1167 | 0.48 | 0.52 |
| MESA | 60.7 | 9.7 | 167.2 | 10.0 | 28.5 | 5.5 | 4814 | 0.48 | 0.52 |
| MGH_AF | 54.2 | 10.4 | 178.5 | 9.9 | 28.4 | 5.3 | 718 | 0.79 | 0.21 |
| VAFAR | 58.2 | 8.6 | 177.8 | 10.3 | 32.7 | 6.6 | 154 | 0.69 | 0.31 |
| VU_AF | 53.0 | 11.1 | 177.6 | 10.2 | 31.2 | 6.9 | 983 | 0.72 | 0.28 |
| WHI | 66.7 | 6.8 | 161.3 | 6.2 | 28.7 | 5.9 | 9950 | 0.00 | 1.00 |

*Supplementary Table 7: Summary of the estimates of heritability for height and BMI from all main analyses performed.*

| Dataset | Experiment info | SNPs | Sample size | GRM Algorithm | N MAF bins | N LD bins | Random effects (GRMs) | Fixed effects (PCs) | Estimates | |
|---|---|---|---|---|---|---|---|---|---|---|
| | | | | | | | | | Height | BMI |
| TOPMed | GREML-SC (HM3 SNPs) | SVM HM3 | 25465 | Alg1 | 1 | 1 | | 20 | 0.48 (0.02) | 0.24 (0.02) |
| | GREML-MS (WGS SNPs) | High quality SVM WGS SNPs | | | 4 | 1 | 4 | 20 HM3 SNPs | 0.48 (0.05) | 0.24 (0.05) |
| | | | | | 4 | 1 | 4 | 160 WGS | 0.45 (0.05) | 0.23 (0.05) |
| | GREML-LDMS (WGS SNPs) (median based) | | | | 4 | 2 | 8 | 20 HM3 | 0.70 (0.09) | 0.29 (0.09) |
| | | | | | 4 | 2 | 8 | 48 WGS | 0.61 (0.09) | 0.25 (0.10) |
| | | | | | 4 | 2 | 8 | 160 WGS | 0.60 (0.09) | 0.23 (0.10) |
| | GREML-LDMS - WGS SNPs – 3 LD bins (tertile based) | | | | 4 | 3 | 12 | 20 HM3 | 0.78 (0.09) | 0.31 (0.10) |
| | | | | | 4 | 3 | 12 | 48 WGS | 0.68 (0.09) | 0.32 (0.10) |
| | | | | | 4 | 3 | 12 | 160 WGS | 0.68 (0.10) | 0.30 (0.10) |
| | GREML-LDMS - WGS SNPs – 4 LD bins (quantile based) | | | | 4 | 4 | 16 | 48 WGS | 0.68 (0.10) | 0.30 (0.10) |
| | | | | | 4 | 4 | 16 | 160 WGS | 0.67 (0.10) | 0.29 (0.10) |
| | | | | | 4 | 4 | 16 | 320 WGS (16*20) | 0.68 (0.10) | 0.28 (0.10) |
| | Enrichment analysis (splitting low MAF and low LD bins into protein-altering and non-protein-altering) | | | | 4 | 2(+1) | 11 | 20 HM3 | 0.70 (0.09) | 0.29 (0.09) |
| | | | | | 4 | 2(+1) | 11 | 48 WGS | 0.61 (0.09) | 0.24 (0.10) |
| | Enrichment and removing extreme diagonal samples | | 22100 | | 4 | 2(+1) | 11 | 20 HM3 | 0.79 (0.10) | 0.26 (0.10) |
| | | | | | 4 | 2(+1) | 11 | 48 WGS | 0.73 (0.10) | 0.21 (0.10) |
| | GREML-LDMS – Different GRM estimator: Average of ratios | | 25465 | Alg 0 | 4 | 2 | 8 | 20 HM3 | 0.74 (0.10) | 0.29 (0.11) |
| | | | | | 4 | 2 | 8 | 160 WGS | 0.63 (0.10) | NA |
| | GREML-LDMS – No High quality filter on variants | All variants WGS | | Alg 1 | 4 | 2 | 8 | 20 HM3 | 0.62 (0.06) | NA |
| | | | | | 4 | 2 | 8 | 160 WGS | 0.62 (0.06) | NA |

| | | | | | | | | | | |
|---|---|---|---|---|---|---|---|---|---|---|
| | GREML-LDMS – TOPMed LD/MAF reference | Intersection UK10K / TOPMed | | | 4 | 2 | 8 | 20 HM3 | 0.60 (0.08) | 0.32 (0.08) |
| | GREML-LDMS – UK10K LD/MAF reference | | | | 4 | 2 | 8 | 20 HM3 | 0.60 (0.06) | 0.30 (0.07) |
| | GREML-LDMS – Imputation from Axiom | Imputed SNPs Rsq > 0.3 | | | 4 | 2 | 8 | 20 HM3 | 0.56 (0.07) | 0.21 (0.07) |
| | GREML-LDMS – Imputation from GSA | | | | 4 | 2 | 8 | 20 HM3 | 0.55 (0.07) | 0.16 (0.07) |
| | GREML-LDMS – Imputation from Infinium | | | | 4 | 2 | 8 | 20 HM3 | 0.50 (0.06) | 0.18 (0.07) |
| | GREML-LDMS – Imputation from Axiom -LD score segment for LD reference | | | | 4 | 2 | 8 | 20 HM3 | 0.51 (0.04) | NA |
| | GREML-LDMS – Comparison Axiom imputed – WGS – Axiom genotypes | Intersection Axiom imputed / TOPMed WGS | | | 4 | 2 | 8 | 20 HM3 | 0.55 (0.07) | 0.18 (0.07) |
| | | | | | 4 | 2 | 8 | 160 WGS | 0.50 (0.07) | 0.13 (0.07) |
| | GREML-LDMS – Comparison Axiom imputed – WGS – TOPMed genotypes | | | | 4 | 2 | 8 | 20 HM3 | 0.62 (0.07) | 0.25 (0.07) |
| | | | | | 4 | 2 | 8 | 160 WGS | 0.56 (0.07) | 0.22 (0.07) |
| | GREML – MS – UKB Exome | Intersection UKB WES TOPMed WGS | | lg1 | 2 | 1 | 2 | 20 HM3 | 0.35 (0.02) | 0.10 (0.02) |
| | GREML – MS – TOPMed | | | | 2 | 1 | 2 | 20 HM3 | 0.30 (0.02) | 0.07 (0.02) |
| | GREML – LDMS – UKB Exome | | | | 2 | 2 | 4 | 20 HM3 | 0.37 (0.02) | 0.13 (0.02) |
| | GREML – LDMS – TOPMed | | | | 2 | 2 | 4 | 20 HM3 | 0.31 (0.02) | 0.09 (0.02) |
| | GREML -LDMS – No heterozygosity QC step | SVM WGS | 28754 | | 4 | 2 | 8 | 20 HM3 | 0.70 (0.08) | 0.36 (0.08) |
| | | | | | 4 | 2 | 8 | 160 WGS | 0.58 (0.08) | 0.31 (0.08) |
| | GREML -LDMS – No heterozygosity QC step – 3 LD bins analysis | | | | 4 | 3 | 12 | 20 HM3 | 0.76 (0.08) | 0.40 (0.09) |
| | | | | | 4 | 3 | 12 | 160 WGS | 0.64 (0.08) | 0.35 (0.09) |
| | GREML -LDMS – No heterozygosity QC step but removal of samples with diagonal values > 1.3 and < 0.7 | | 25333 | | 4 | 2 | 8 | 20 HM3 | 0.81 (0.09) | 0.37 (0.09) |
| | | | | | 4 | 2 | 8 | 160 WGS | 0.71 (0.09) | 0.31 (0.10) |
| | GREML -LDMS – No heterozygosity QC step but removed pairs with off-diagonal values across all WGS GRMs > 0.1 | | 26698 | | 4 | 2 | 8 | 20 HM3 | 0.71 (0.08) | 0.35 (0.09) |
| | | | | | 4 | 2 | 8 | 160 WGS | 0.59 (0.09) | 0.29 (0.09) |
| | | | 24233 | | 4 | 2 | 8 | 20 HM3 | 0.82 (0.09) | 0.38 (0.10) |

| | | | | | | | | | | |
|---|---|---|---|---|---|---|---|---|---|---|
| | GREML -LDMS – No heterozygosity QC step but removed diagonals and off-diagonals extreme values across all GRMs | | | | 4 | 2 | 8 | 160 WGS | 0.71 (0.10) | 0.31 (0.10) |
| UK Biobank Whole Exome | GREML-LDMS + HM3 GRM | WES + HM3 | 35867 | Alg0 | 7 | 2 | 14 + 1 (HM3) | 20 HM3 | 0.62 (0.04) | 0.33 (0.04) |
| | GREML-LDMS + HM3 GRM – fitting birth coordinates | | | | 7 | 2 | 14 + 1 (HM3) | 20 HM3 | 0.61 (0.04) | 0.33 (0.04) |
| | GREML-LDMS + HM3 GRM | | | | 7 | 2 | 14 + 1 (HM3) | 280 WES (20 per bin) | 0.59 (0.04) | 0.31 (0.04) |
| | GREML-LDMS + HM3 GRM – fitting birth coordinates | | | | 7 | 2 | 14 + 1 (HM3) | 280 WES (20 per bin) | 0.56 (0.04) | 0.25 (0.05) |
| | GREML-LDMS + HM3 GRM – fitting sequencing center | | | | 7 | 2 | 14 + 1 (HM3) | 280 WES (20 per bin) | 0.57 (0.04) | 0.28 (0.04) |

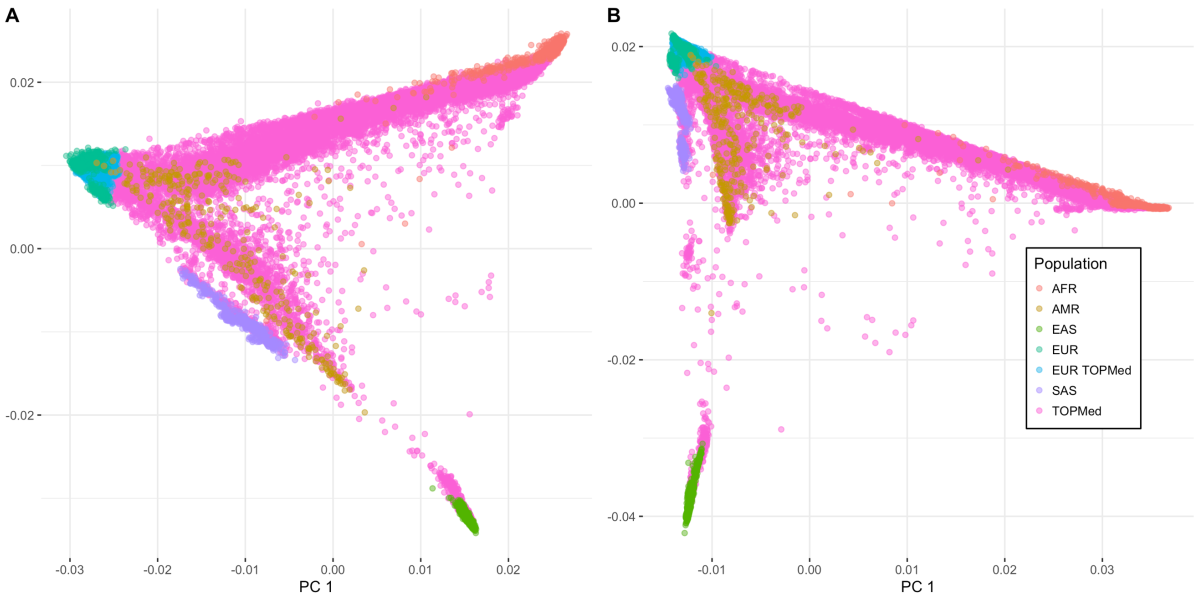


*Supplementary Figure 1: Principal Component Analysis plot of individuals compared to 1000 Genomes populations. (A) PCA of TOPMed samples and 1000G populations using 580k common SNPs. (B) PCA of TOPMed samples and 1000G populations using ~1.3M rare SNPs.*

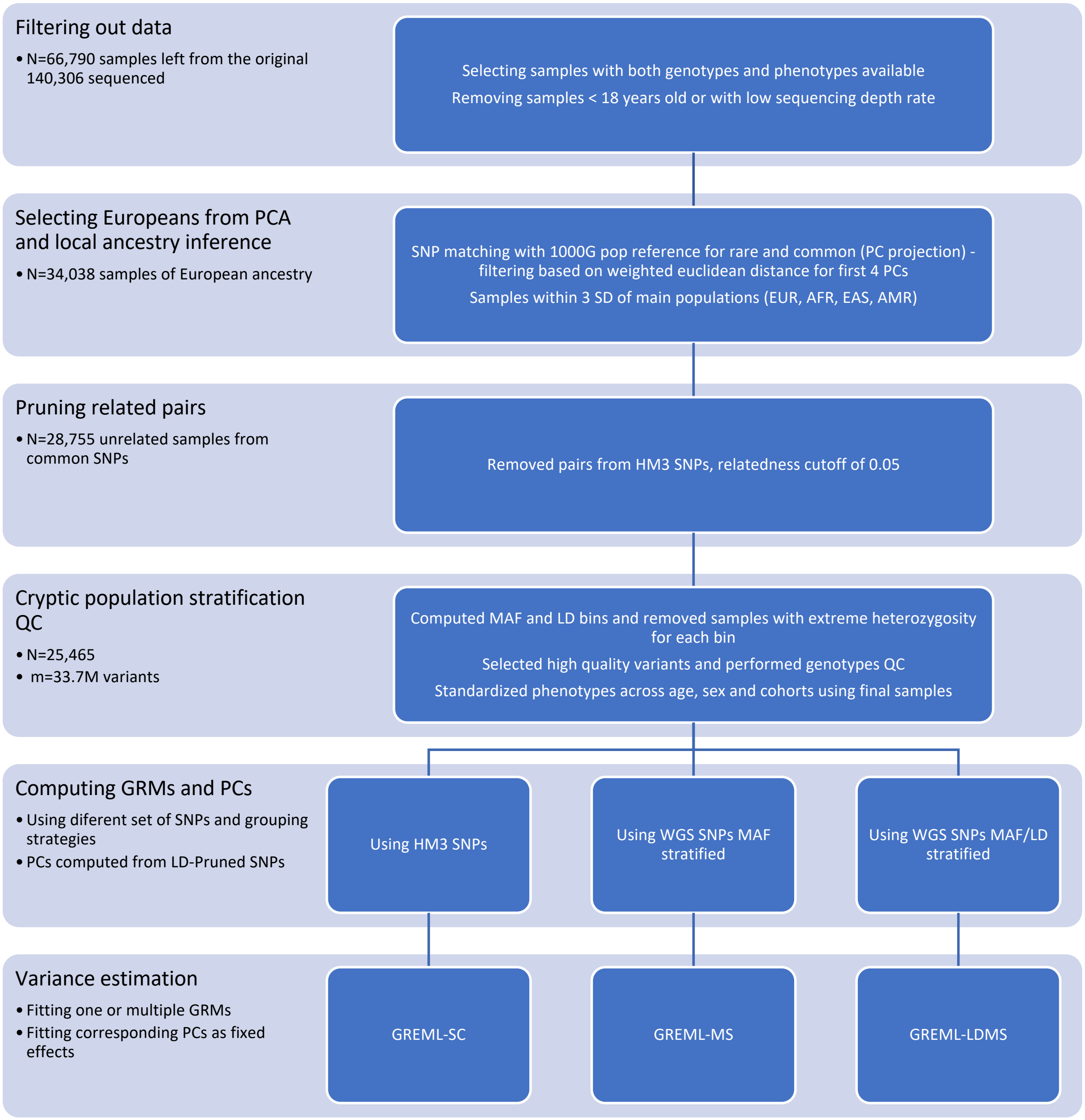


*Supplementary Figure 2: Quality control and analysis pipeline for TOPMed WGS data to estimate trait variance.*

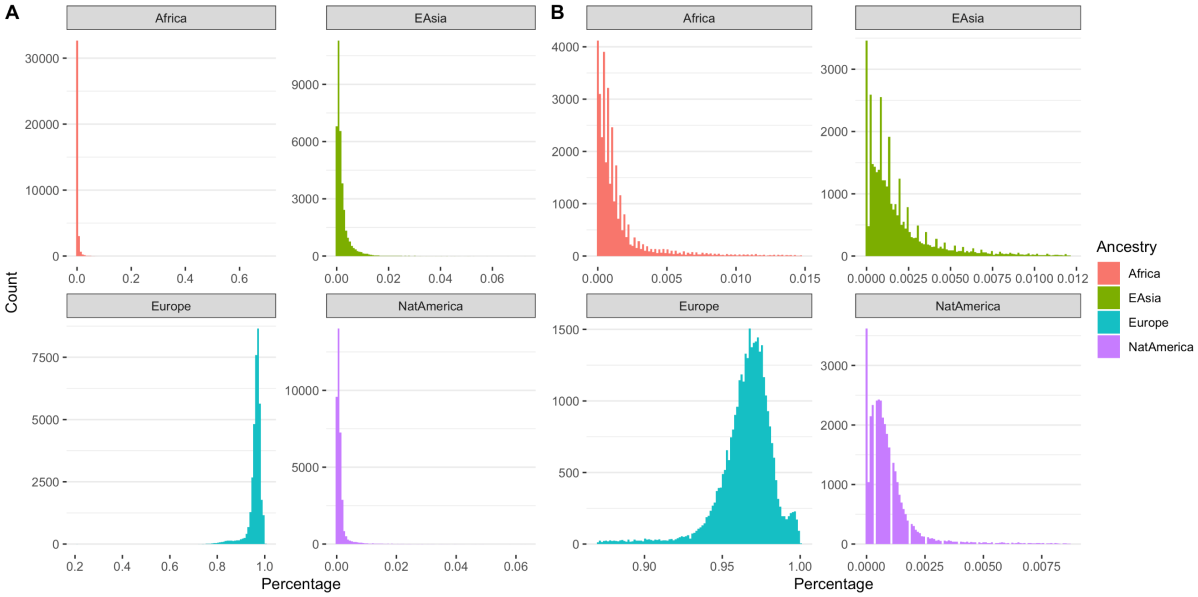


*Supplementary Figure 3: Proportion of ancestry according to RFMix reference before and after filtering samples further away from 3 standard deviations of each reference population. (A) Before filtering samples (N=36,938 ). (B) After filtering samples (N=34,038).*

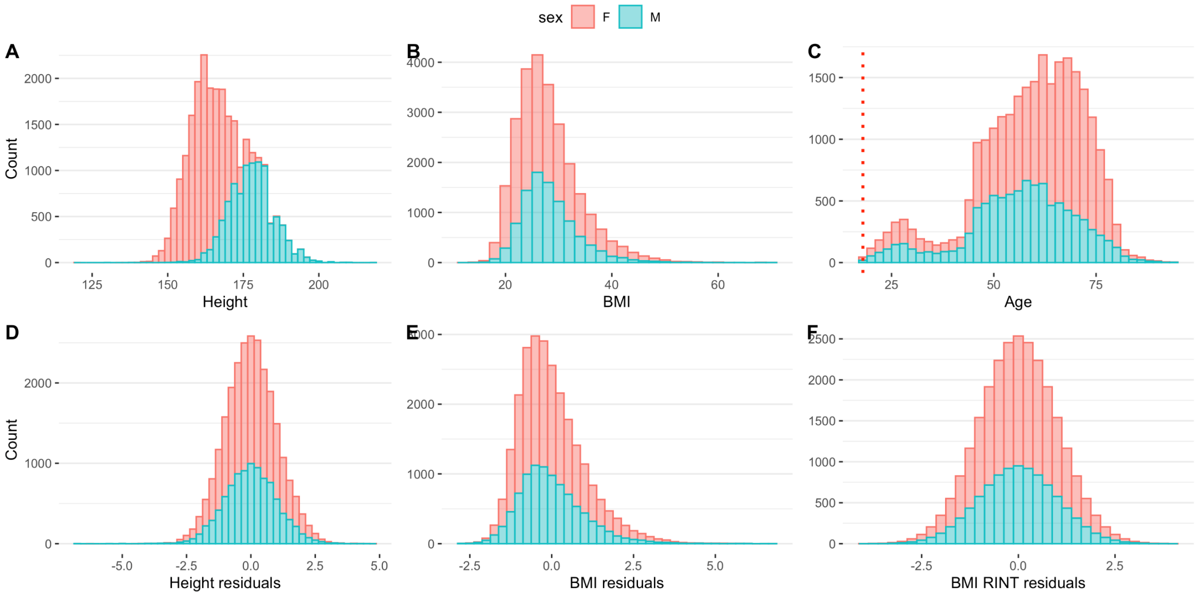


*Supplementary Figure 4: Distribution of N=25,465 phenotypes (unrelated Europeans) in the dataset before and after the QC process. (A) Distribution of height before standardization. (B) Distribution of BMI before standardization. (C) Distribution of age in the dataset with indivduals <18 years old removed. (D) Distribution of the residuals for height after QC standardization. (E) Distribution of the residuals for BMI after QC standardization. (F) Distribution of the residuals for BMI with a rank inverse normal transformation after QC standardization. The skewness in distribution of the BMI is removed by the transformation.*

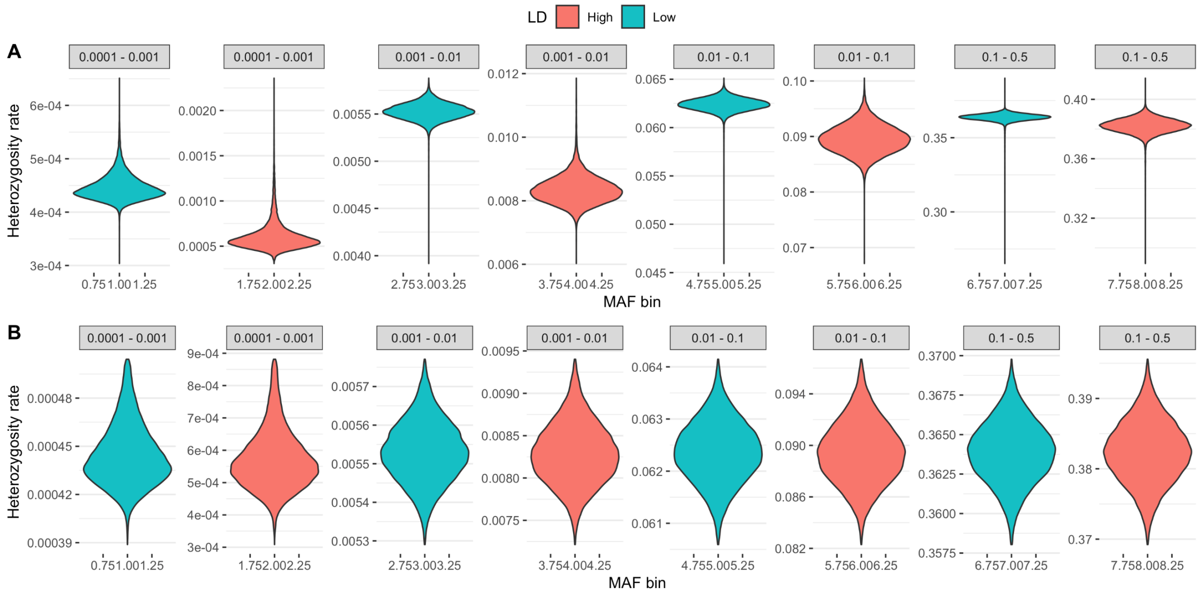


*Supplementary Figure 5: Distribution of sample heterozygosity for each MAF and LD grouping. (A) Distribution before filtering (N=28,755). (B) Distribution after 4 rounds of filtering out samples further away than 3 standard deviations for each distribution mean (N=25,465).*

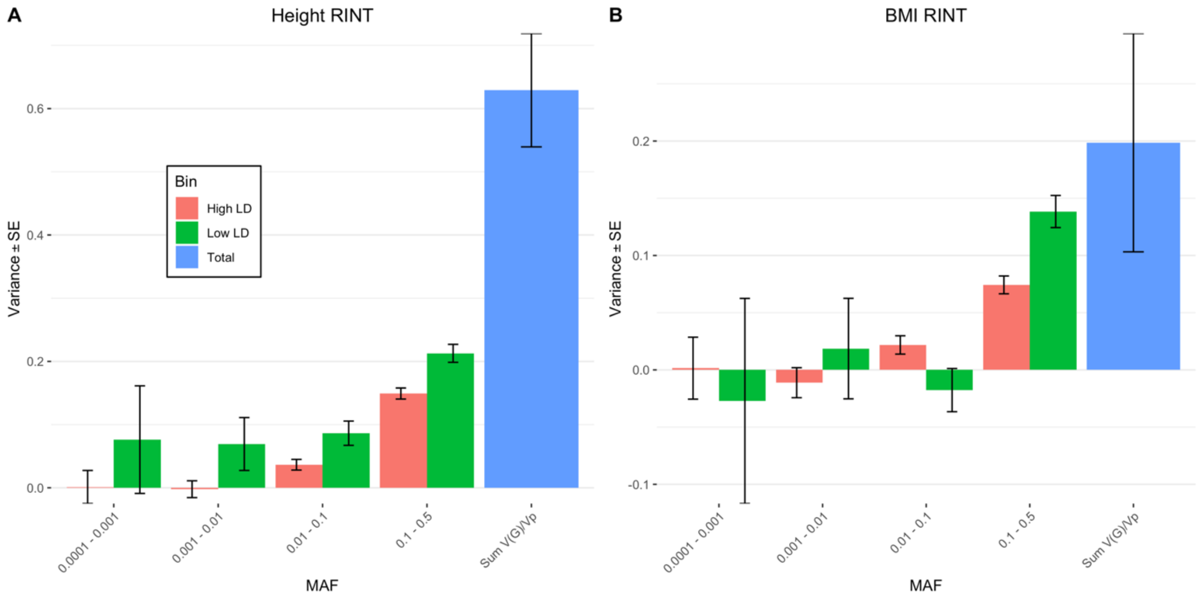


*Supplementary Figure 6: GREML-LDMS estimates using WGS (~33.7M variants) with a rank inverse normal transformation (RINT) correcting by 48 PCs. (A) Estimates for $height_{RINT}$ ~0.63 (SE 0.09) are consistent with untransformed trait. (B) Total estimates for $BMI_{RINT}$ are lower than BMI ~0.20 ( 0.10).*

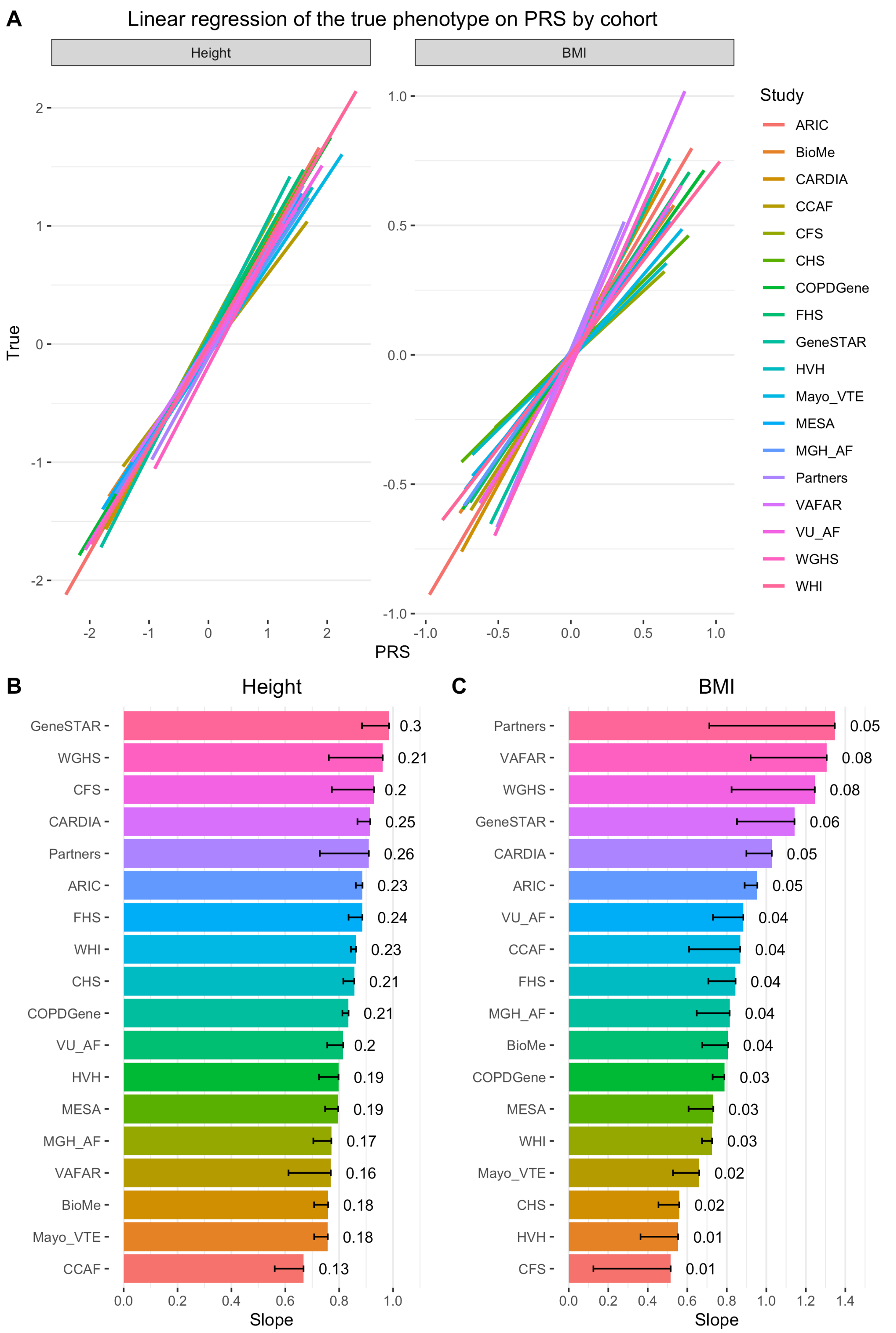


*Supplementary Figure 7: PGS in n=25,465 samples from 1360 and 449 independent SNPS associated with height and BMI respectively in the UKB matching TOPMed data set. (A). Fitted regression slopes for samples in each 18 cohorts. (B). Estimates of the individual slopes (x-axis) and the proportion of phenotypic variance explained (numbers displayed) for each TOPMed cohort.*

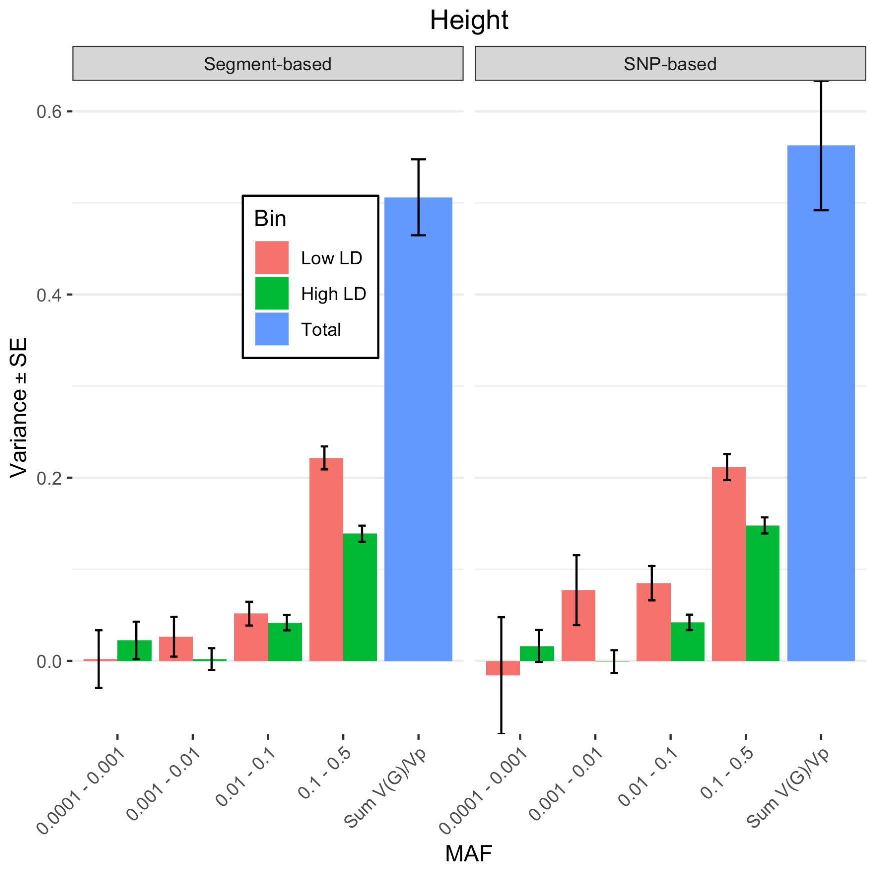


*Supplementary Figure 8: Estimates of SNP-based heritability ($h_{SNP}^2$) from a GREML-LDMS analysis based on 8 bins (4 MAF * 2 LD bins) for SNPs imputed on Axiom array (~20.4M SNPs), corrected for 20 HM3 SNPs PCs. LD stratification was done using either on segment-based LD value or individual-SNP LD value. Estimates went from $h_{SNP}^2$ ~0.51 (SE 0.04) using a segment-based LD value stratification to $h_{SNP}^2$ ~0.56 (SE 0.07) using an individual-SNP LD value stratification.*

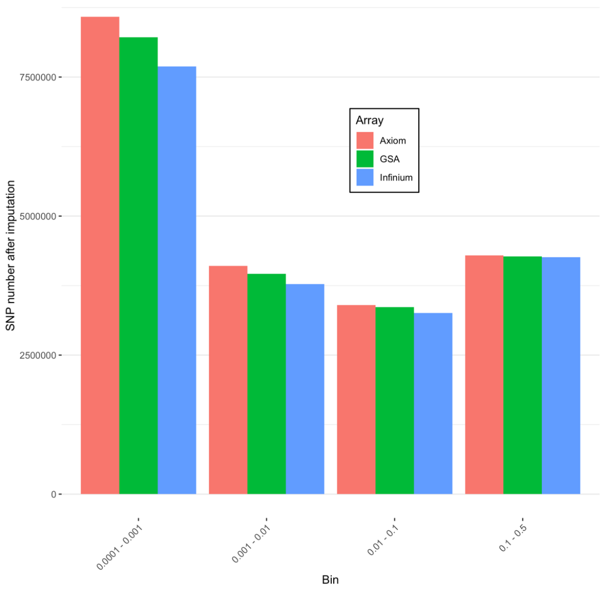


*Supplementary Figure 9: Number of SNPs in each MAF bin after imputation and QC process, for each of the Illumina InfiniumCore24, GSA-24 and Affymetrix Axiom arrays. Total number of SNPs is between ~19.0 and ~20.0 M SNPs.*

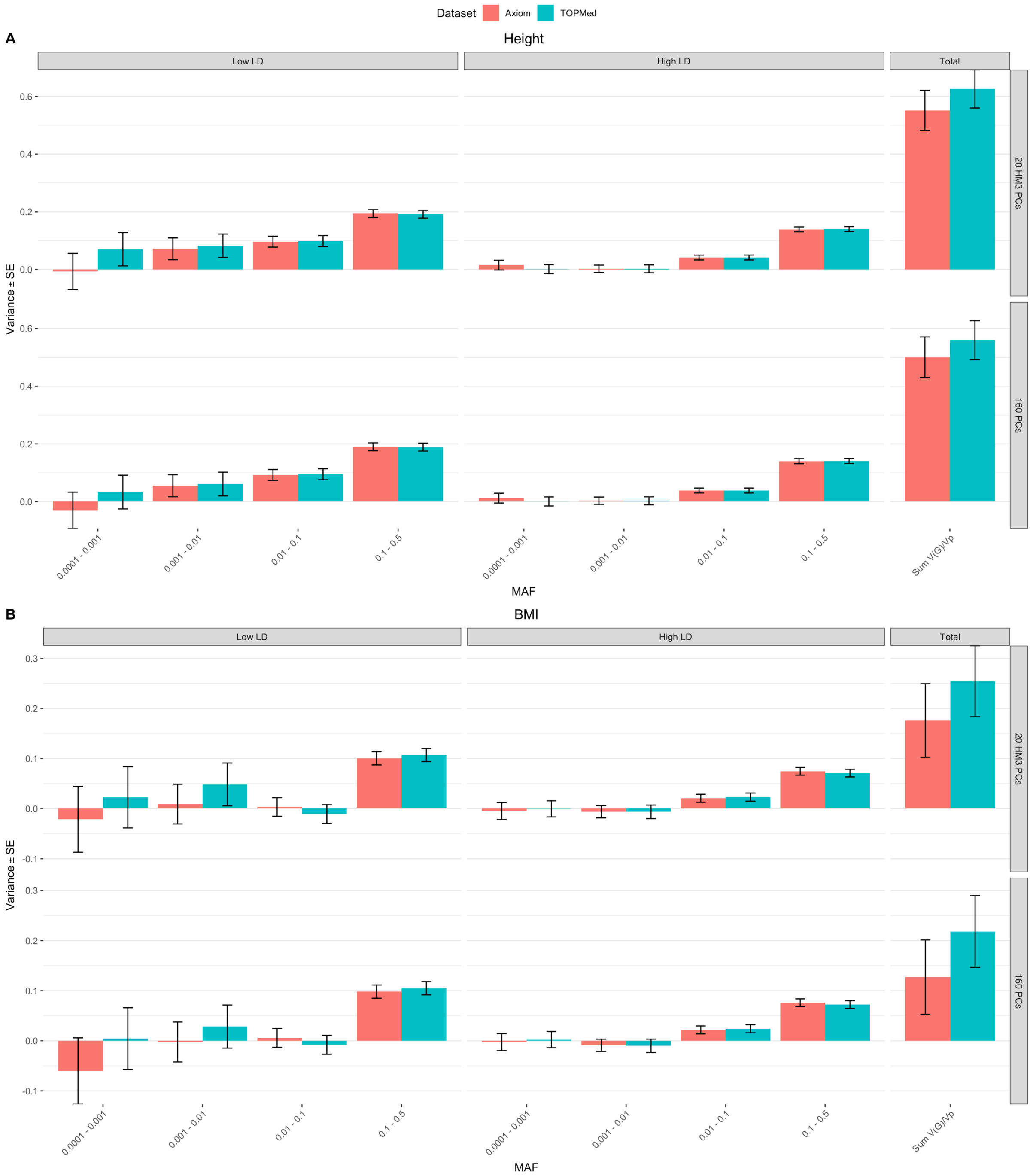


*Supplementary Figure 10: GREML-LDMS estimates stratified in 14 MAF and LD bins, using subset of SNPs in common between TOPMed dataset and imputed dataset from Axiom array SNPs (~17.9M SNPs in common) and correcting by either 20 PCs computed from HM3 independent SNPs or 160 PCs from each MAF/LD bin. Estimates are consistently higher for the TOPMed dataset, with most of the difference coming from the variants in the 0.0001 to 0.001 MAF range in low LD. This difference could be due to imputation errors as common variant bins seems consistent between the two datasets.*

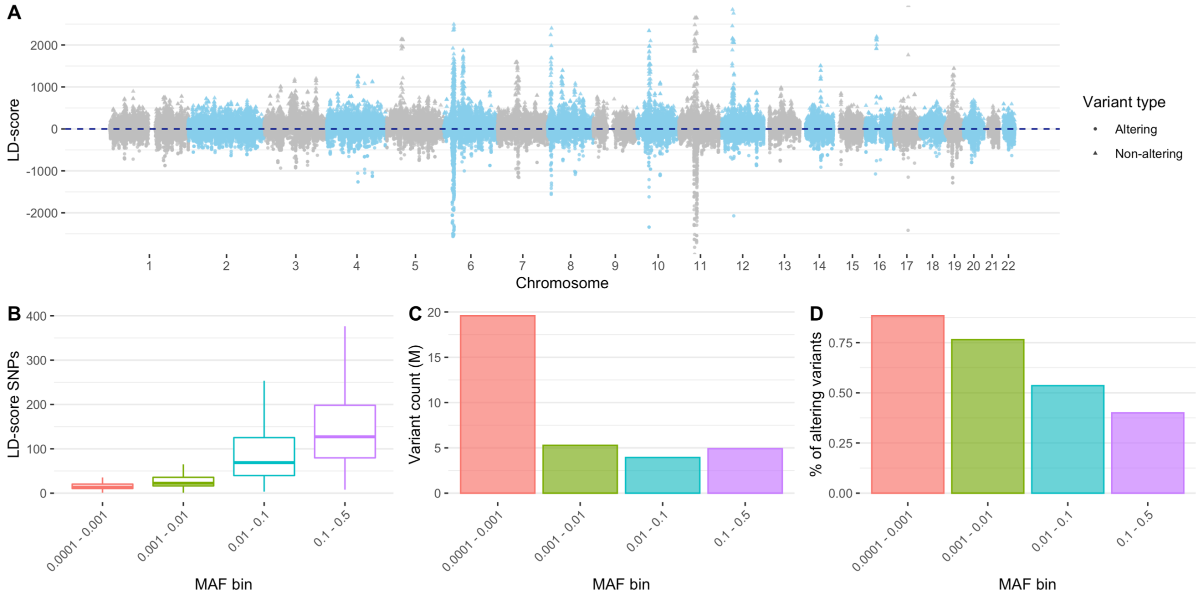

*Supplementary Figure 11: (A) LD score value across the genome for a subset of 350k protein-altering and non-protein-altering variants. LD value were calculated within each of the 4 MAF bin, LD values > 3000 are not shown (B) Boxplot of the distribution of individual SNPs LD values within each bin. (C) Number of SNPs for each of the four MAF bin. (D) Fraction of the high impacting variants as a percentage of the total number of variants in the bin.*

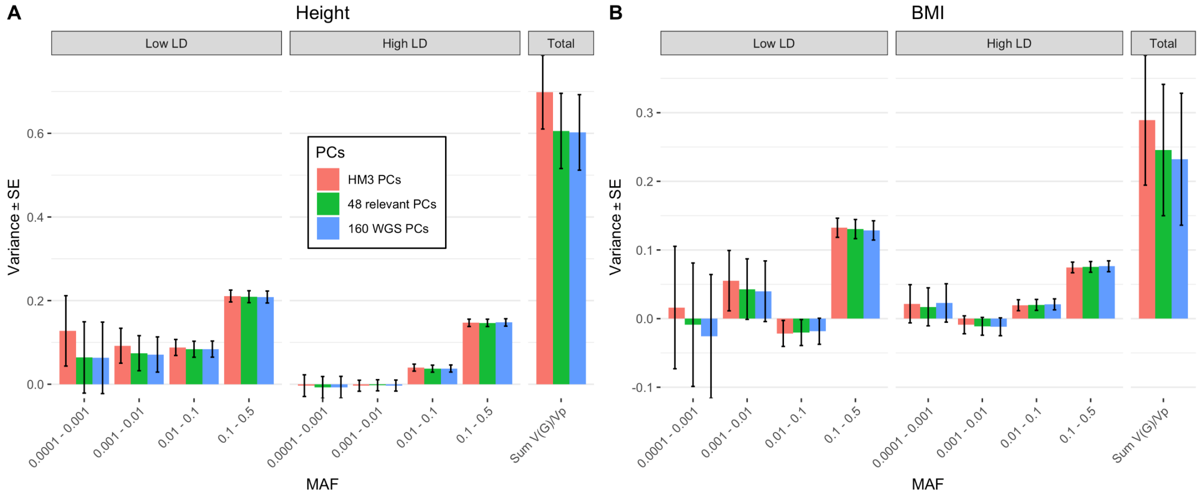

*Supplementary Figure 12: GREML-LDMS estimates from WGS data (~33.7M variants) stratified in 8 bins (4 MAF bins in 2 LD bins) with correction for 20 PCs (on HM3 SNPs), 48 PCs reflecting population stratification (Supplementary Figure 17) or 160 PCs (20*8 bins). (A) Estimates for height with $h^2_{WGS}$ at ~0.60 - ~0.70 (SE ~0.09). (B) Estimates for BMI with hWGS2 at ~0.25 - ~0.29 (SE 0.09 – 0.10). The number of variants in each of the 4 MAF bins (twice the number in each LD bin) is, from the lowest to highest MAF bins, 19.3M, 5.3M, 3.9M and 4.9M, respectively (Supplementary Table 4 and Supplementary Figure 11).*

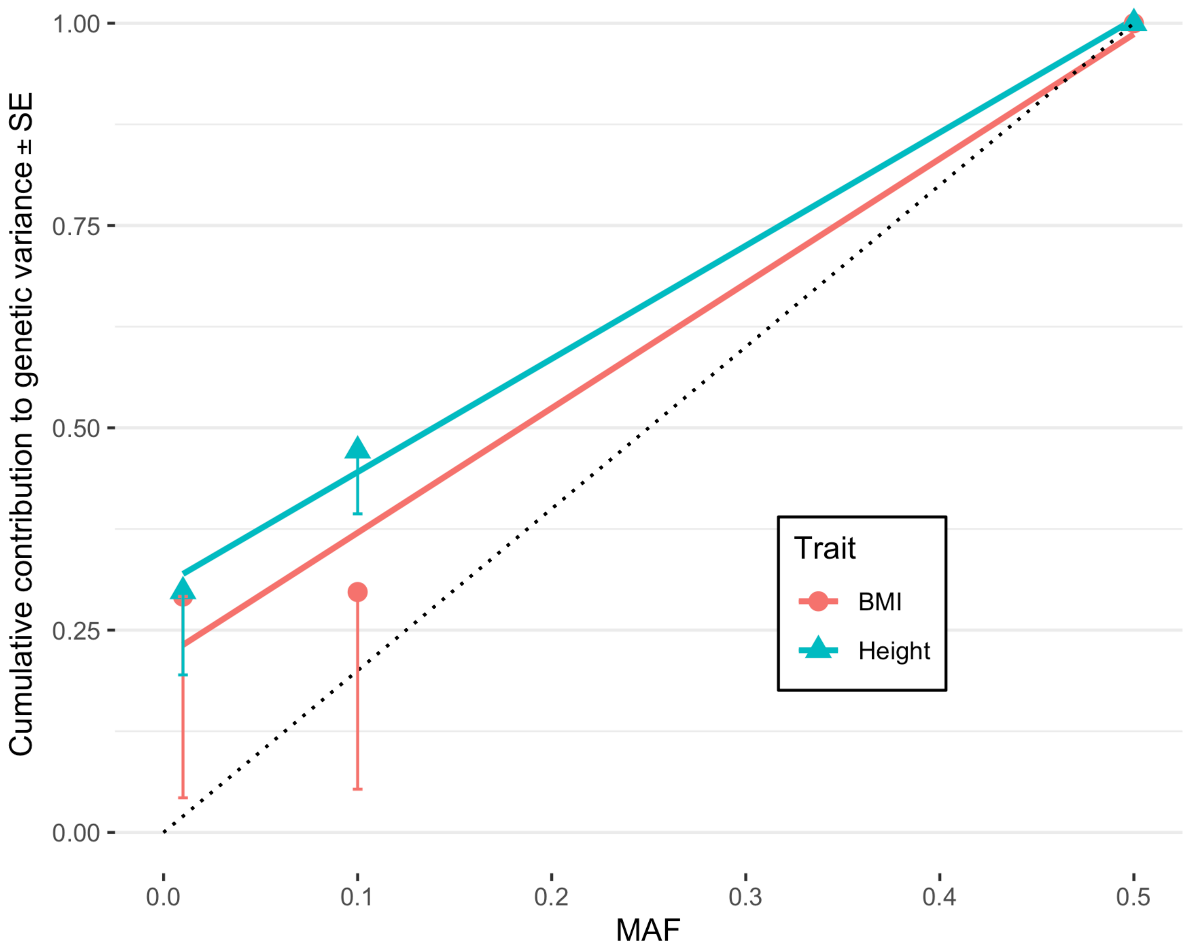


*Supplementary Figure 13: Estimate of the cumulative contribution of variants, for height and BMI, from GREML-LDMS analysis. The dotted line represents the expected contribution under a neutral evolutionary model. The deviation from this dotted line suggest that height and BMI are under negative selection. For each trait, a linear model has been fitted to better visualise the trait selection.*

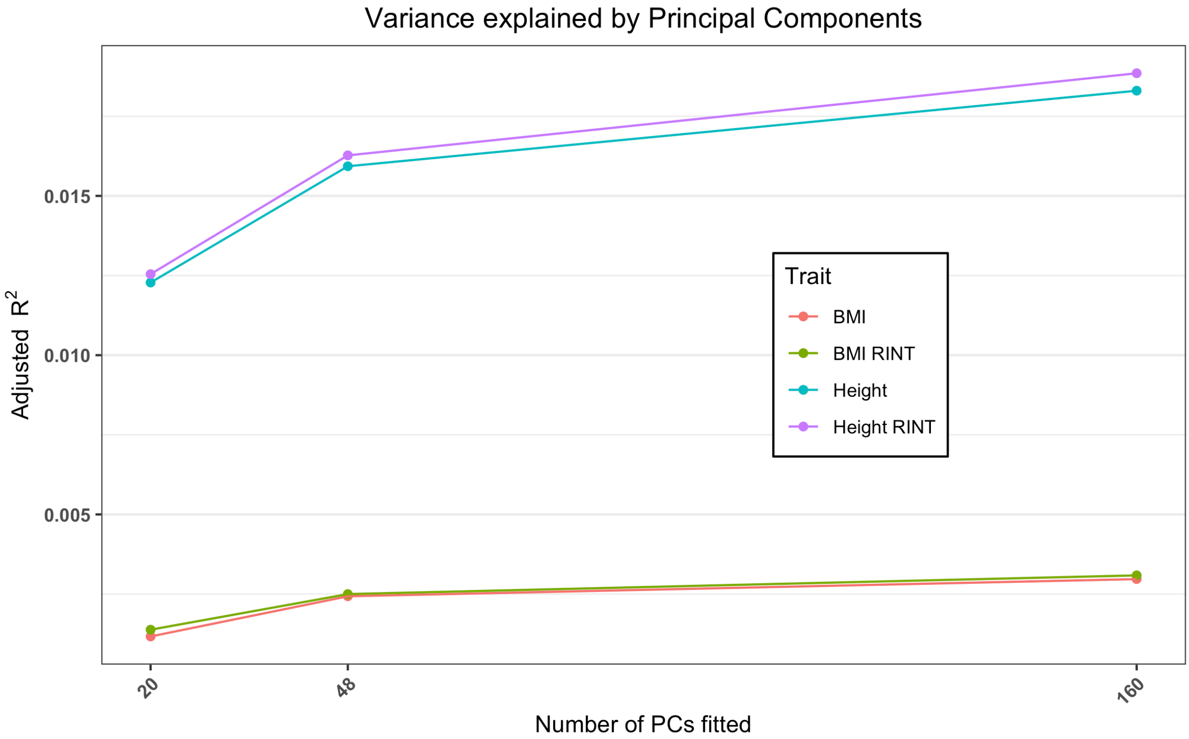

*Supplementary Figure 14: Adjusted $R^2$ of a linear model regressing the phenotype against 20 PCs from LD-pruned HM3 SNPs and 48 (calculated from 8 MAF/LD bins of LD-pruned SNPs) or 160 (calculated from 4 MAF bins * 2 LD bins, on independent SNPs) PCs calculated from TOPMed WGS dataset.*

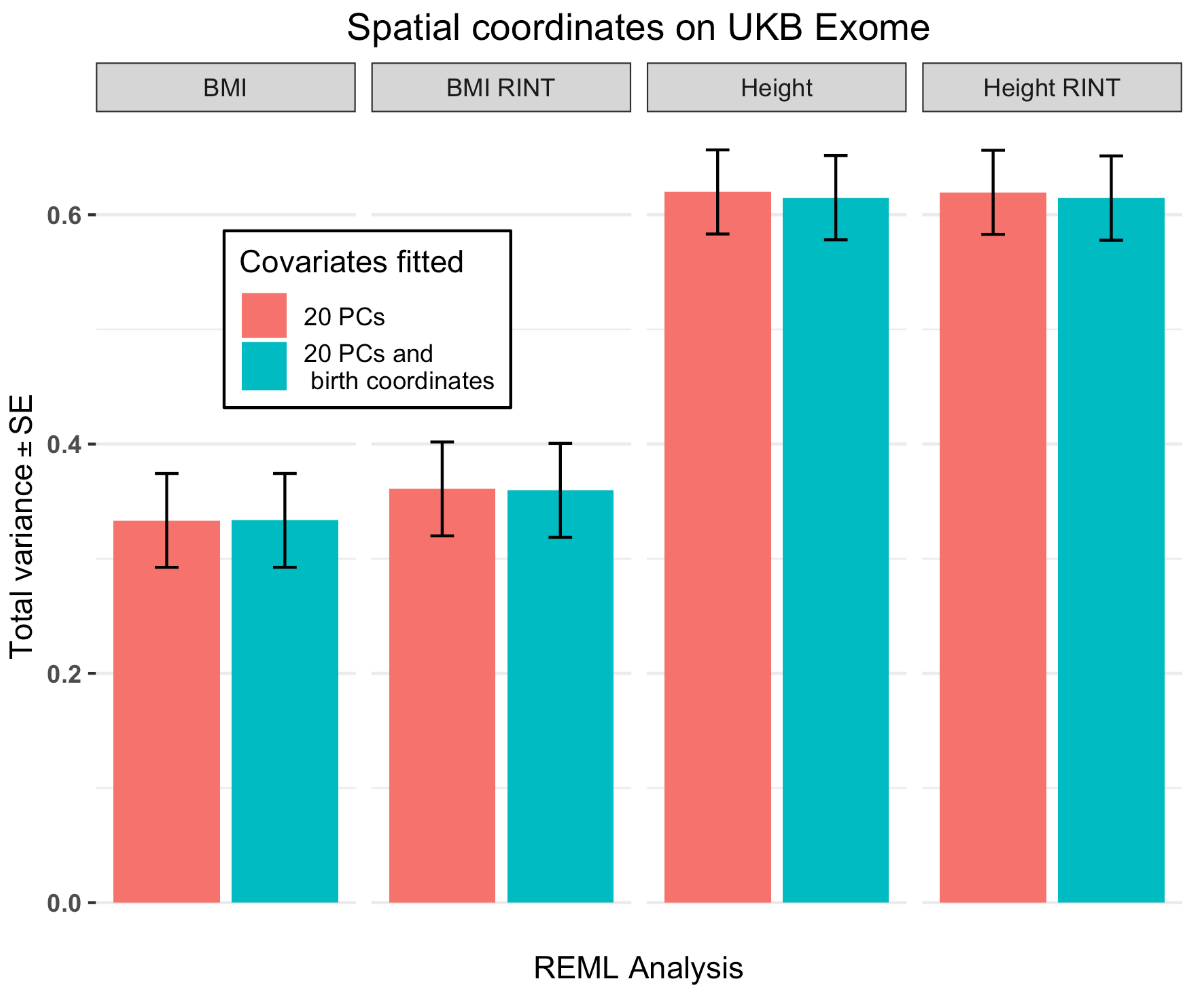


*Supplementary Figure 15: GREML-LDMS estimates fitting 14 bins from UKB Exome data and one bin of HM3 imputed common SNPs. Estimates fitting either 20 PCs calculated from HM3 SNPs or 20 PCs and the north and east birth coordinates scaled on a 0,1 range. Estimates are of ~0.61-0.62 (SE 0.04) for height and height$_{RINT}$, 0.33 (SE 0.04) for BMI and 0.36 (SE 0.04) for BMI$_{RINT}$.*

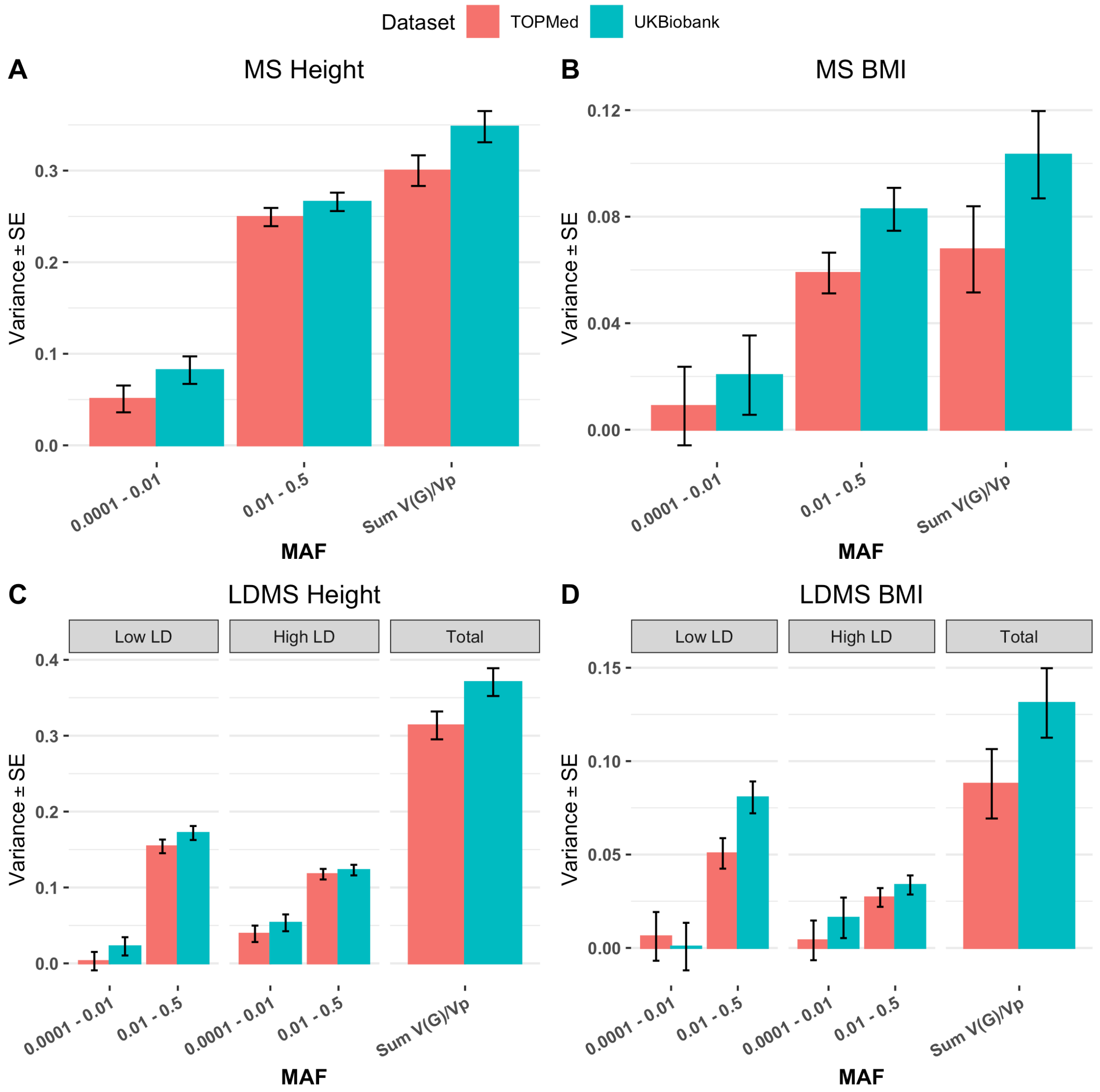


*Supplementary Figure 16: GREML estimates using exome SNPs present in both TOPMed and UKB Exome dataset. Only 2 MAF groupings are fitted in this mode with rare (0.0001 < MAF < 0.01) or common (0.1 < MAF < 0.5) variants. The sample size of UKB Exome dataset was downsampled to match TOPMed's sampe size (25,465 unrelated european individuals). Rare variants not following a normal distribution were removed (Online Methods). We further investigated the effect of LD stratification with the same set of variants in (C) and (D). For TOPMed and UKB Exome respectively. (A) Total estimates for height of 0.30 (SE 0.02) and 0.35 (SE 0.02). (B) Estimates for BMI of 0.07 (SE 0.02) and 0.10 (SE 0.02) for BMI. (C) GREML-LDMS estimates for height 0.31 (SE 0.02) and 0.37 (SE 0.02). (D) Estimates for BMI of 0.09 (SE 0.02) and 0.13 (SE 0.02).*

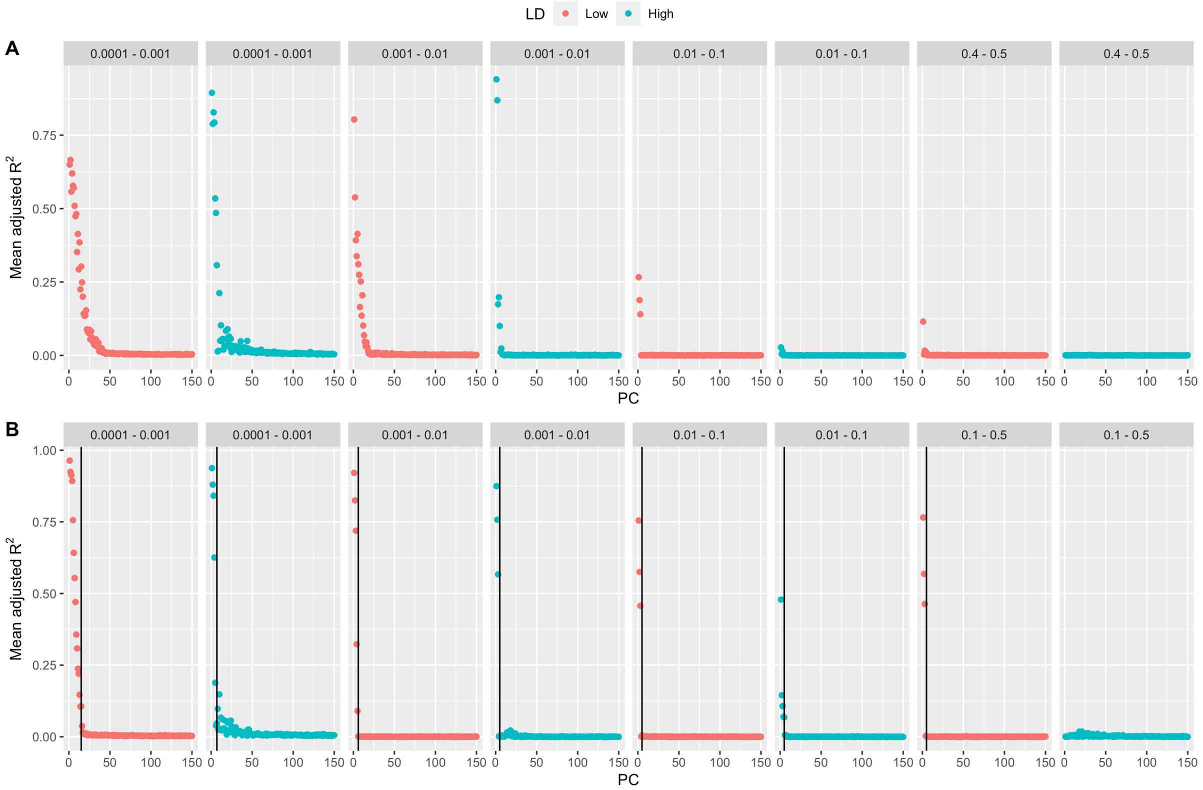


*Supplementary Figure 17: Adjusted $R^2$ from fitting 150 PCs computed from LD-pruned SNPs for different MAF and LD bins on odd and even chromosomes GRMs. The mean $R^2$ from fitting a PC from one set on all the other set PCs is shown. As we do not expect Inter-chromosomal correlations under random mating, observing a $R^2 > 0$ between sets of chromosomes most likely show some population stratification. (A) Mean $R^2$ for the UKB Exome samples. (B) Mean $R^2$for the TOPMed samples. Vertical lines show the threshold determining the best number of PCs to fit to account for population stratification. The thresholds for each MAF/LD bin was computed from a segmented regression model. From this analysis, the number of PCs to fit for each bin to correct for population stratification is respectively 15, 7, 6, 5, 5, 5, 5, 0 for a total of 48 PCs.*

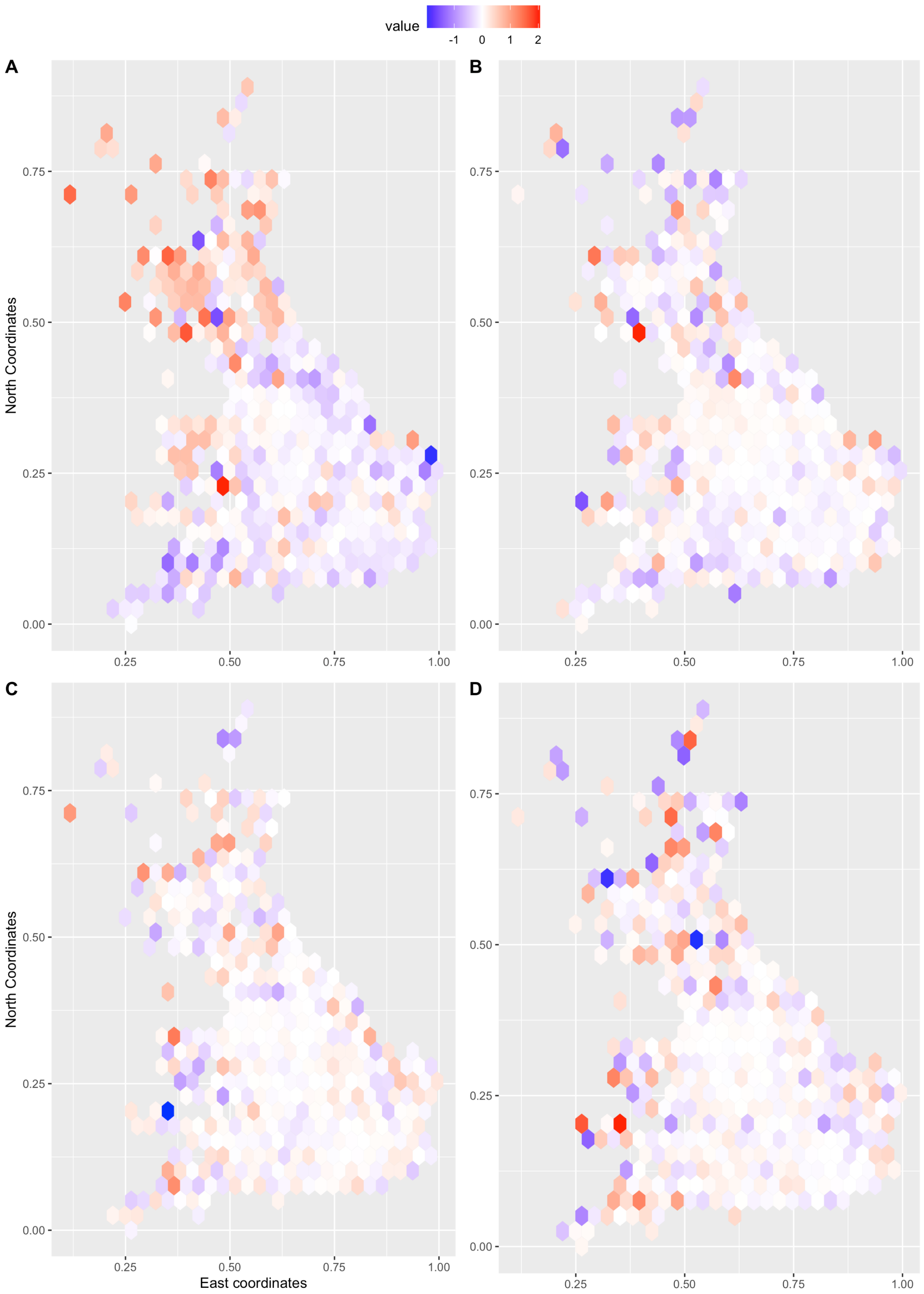


*Supplementary Figure 18: Product of $PC_{i=1,100} = PC^{i}_{Odd} * PC^{i}_{Even}$. for each principal component for each sample, further transformed by centering and applying a RINT transformation to smoothen the effect of outliers. For each individual, this product of odd-even chromosomes PCs was plotted according to their East and North birth coordinates.(A) PC5 on rare variants (0.0001 < MAF < 0.001) in low LD (. (B) PC100 on rare variants. (C) PC5 on common variants (0.4 < MAF<0.5) . (D) PC100 on common variants. We can see a larger stratification for (A) potentially indicating a north-south stratification.*

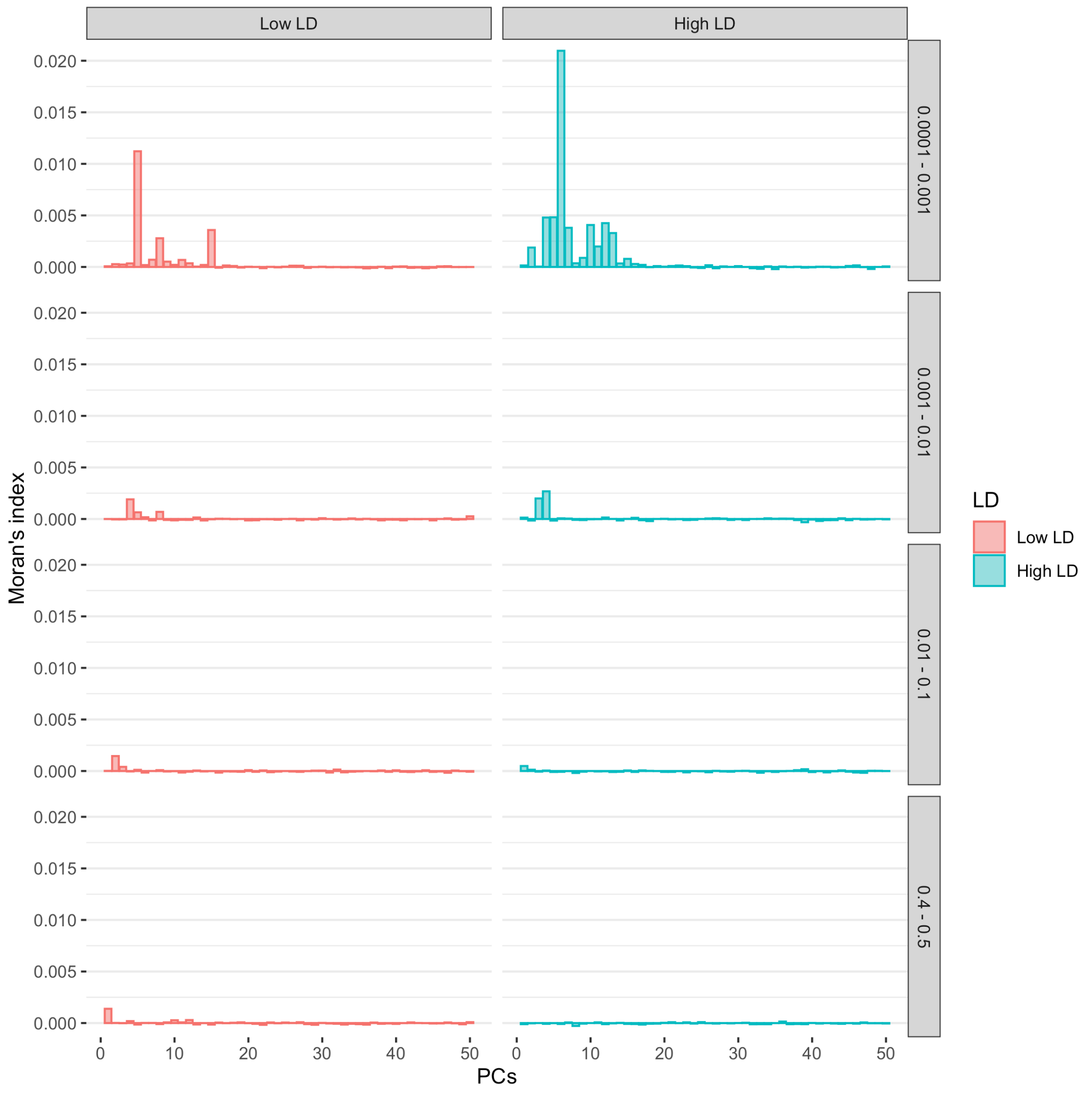


*Supplementary Figure 19: Moran's I auto-correlation index calculated of UK-Biobank Exome samples. This index was calculated across all the samples for each of the first 50s on 3 rare MAF bins and one common MAF bin partitioned according to SNP-based LD. The product of* $PC_{i=1,50} = PC^{i}_{Odd} * PC^{i}_{Even}$ *was used to compute Moran's I index. We can observe a larger spatial correlation for the first PCs of the rare variants bins.*

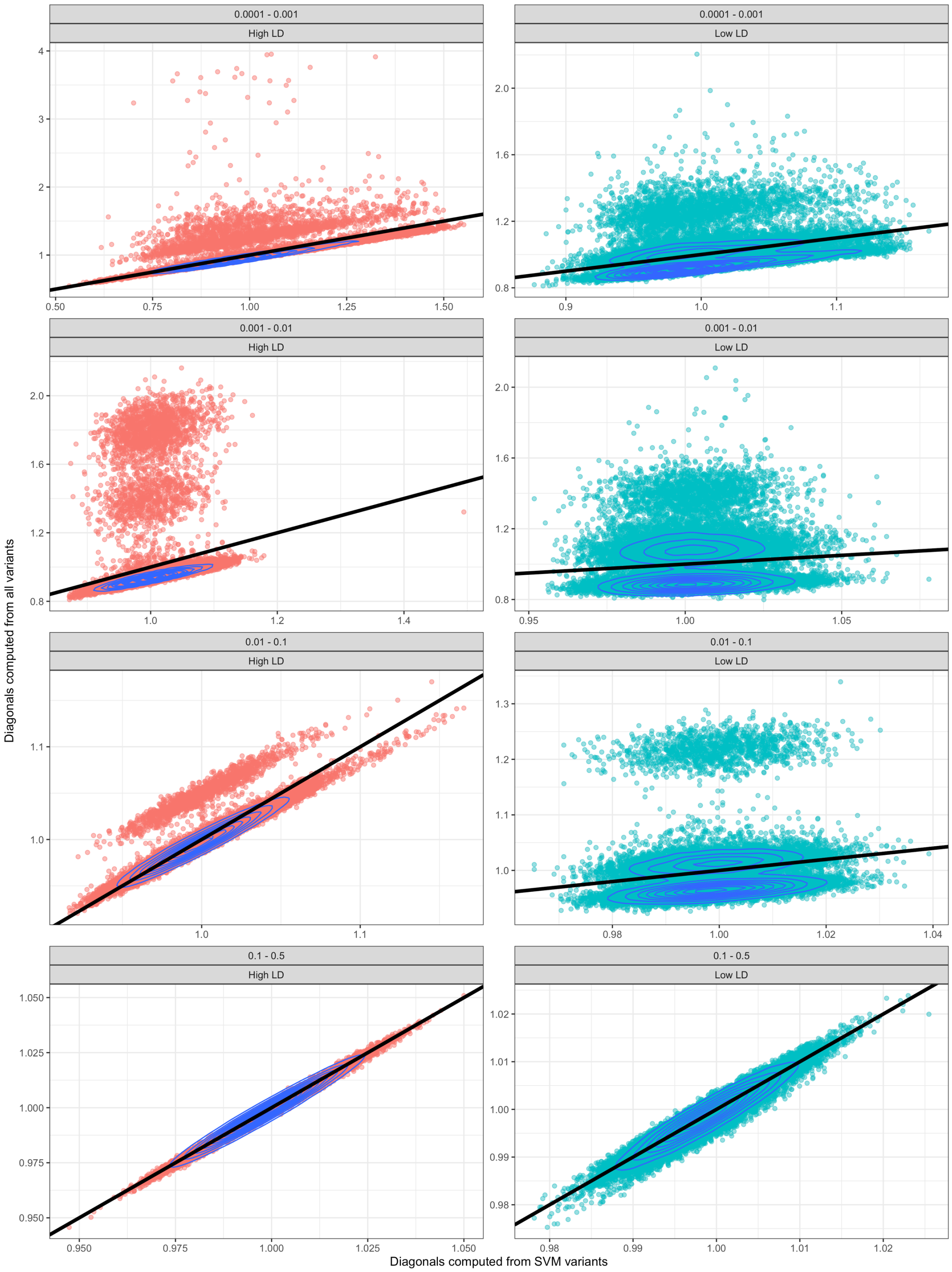


*Supplementary Figure 20: GRM diagonals comparison between GRM computed from the high quality variants (SVM) or all genotyped variants. GRMs were computed using the ratio of average method. There are N=25,465 samples per bin*

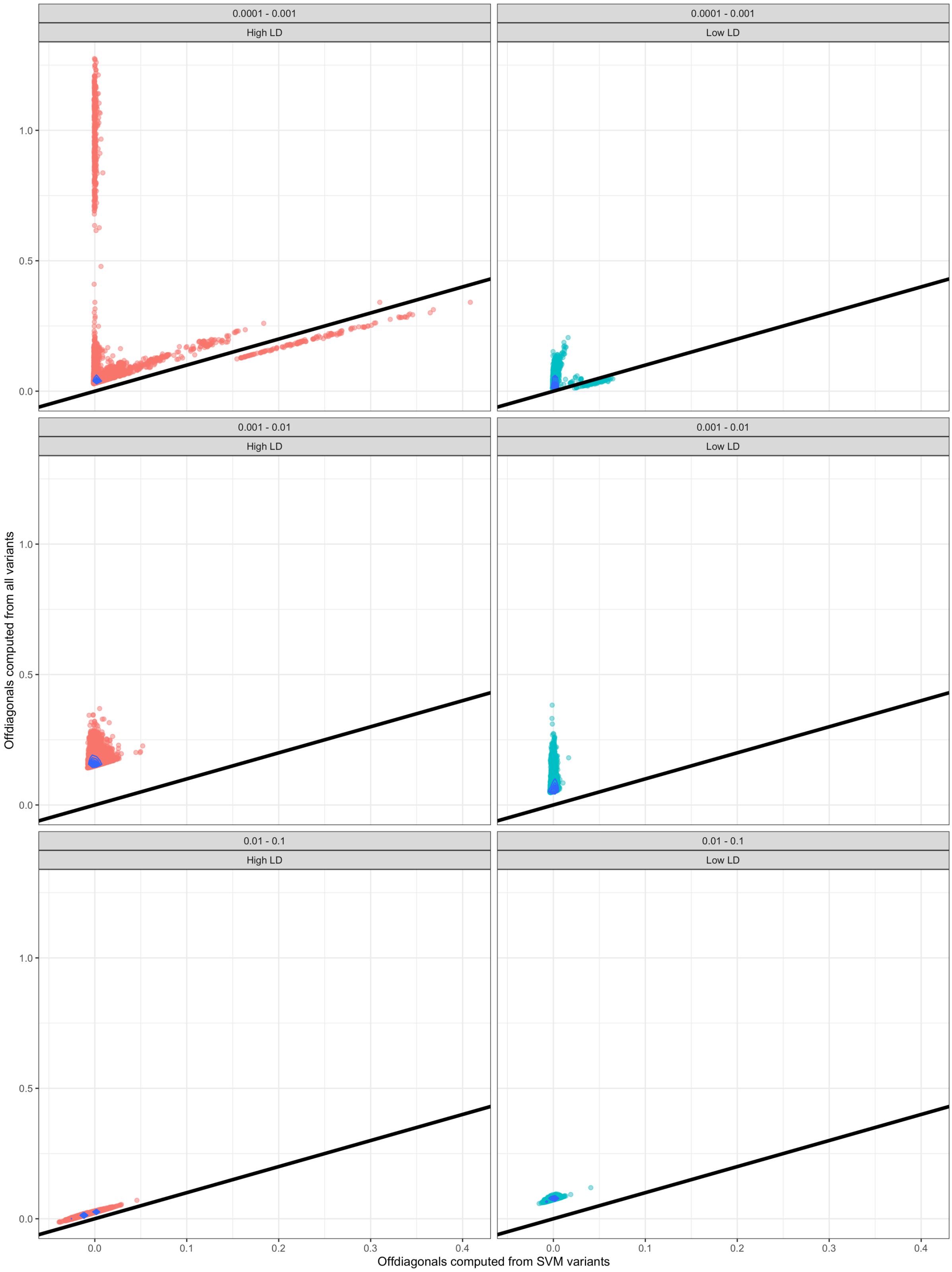


*Supplementary Figure 21: GRM off-diagonals comparison between GRM computed from the high quality variants (SVM) or all genotyped variants. GRM were computed using the ratio of average method. Only the largest 20,000 differences are shown for each bin with a minimum threshold of 1%. Common variants bins (0.1 < MAF < 0.5) did not show differences between the two methods larger than 1%.*

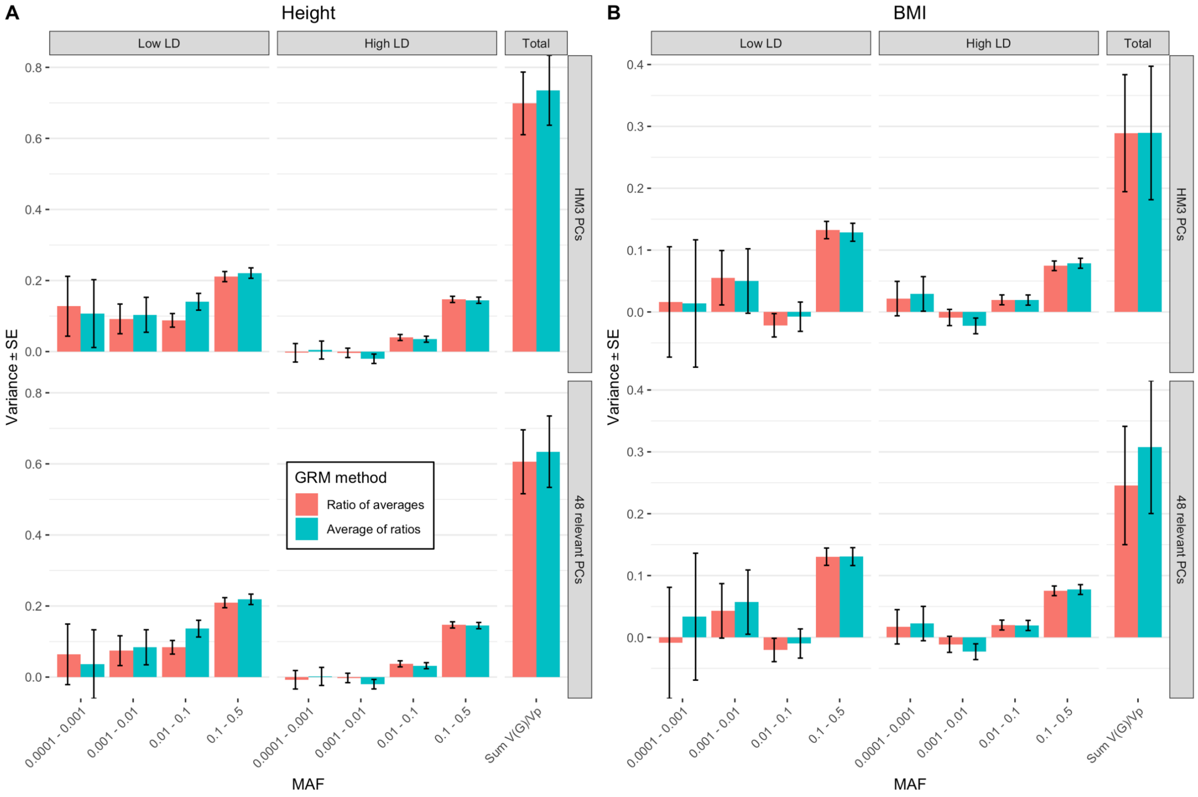


*Supplementary Figure 22: GREML-LDMS using WGS (~33.7M variants) estimates using either the average over loci of ratios (blue) or the ratio over averages of loci (red) methods to calculate the GRM values. (A) Estimates for height fitting 8 bins: 0.70 - 0.74 (SE 0.09 - 0.10) fitting 20 HM3 PCs and 0.61 – 0.63 (SE 0.09 – 0.10) fitting 48 PCs. (B) Estimates for BMI fitting 8 bins: 0.29 - 0.29 (SE 0.09 - 0.11) fitting 20 HM3 PCs and 0.25 – 0.31 (SE 0.10 – 0.11) fitting 48 PCs.*

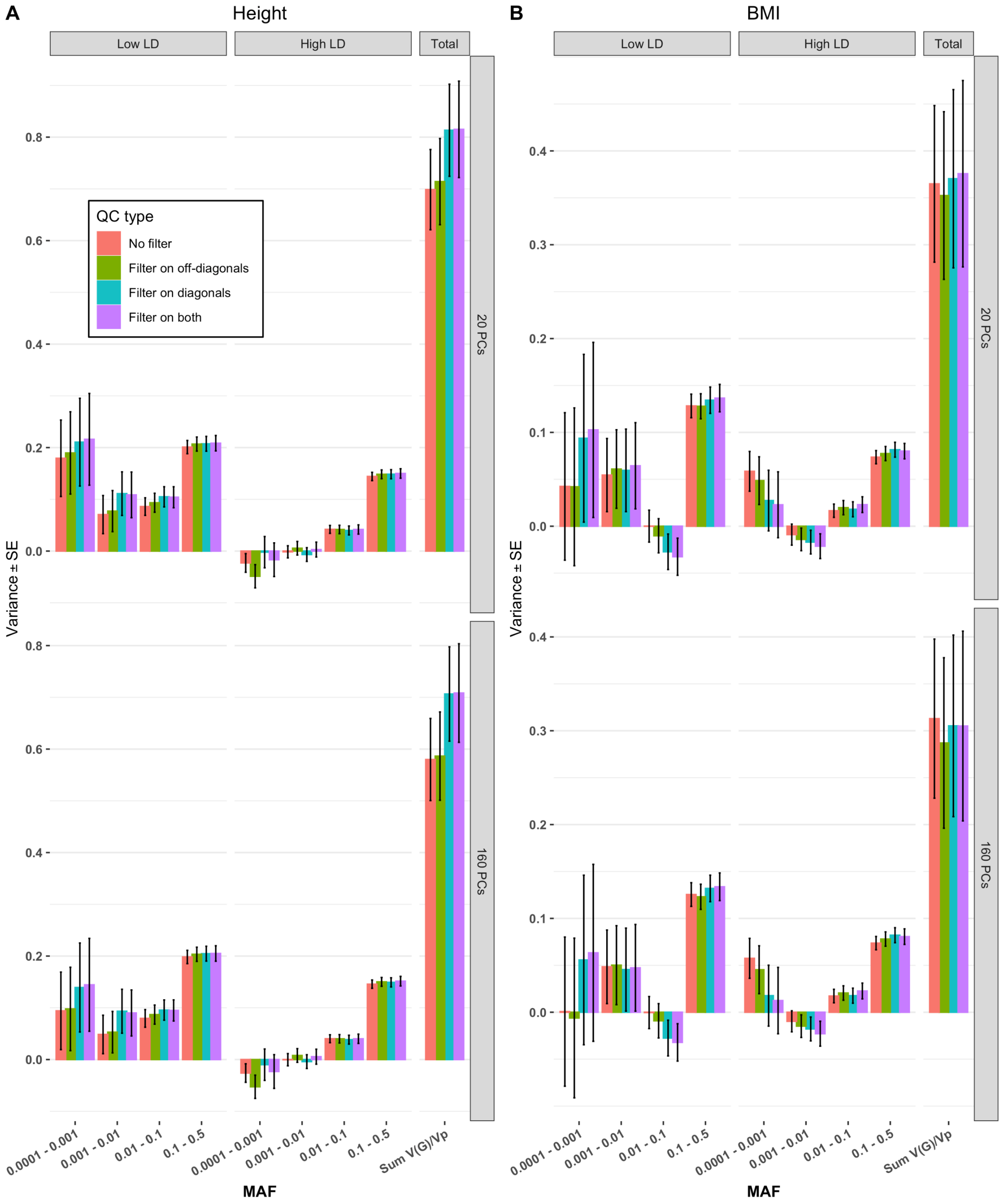


*Supplementary Figure 23: GREML-LDMS estimates by removing samples from the full GRM of Europeans (before QC on sample heterozygosity). Samples were removed either based on their off-diagonal values (>0.1) across all GRMs (all residual relatedness coming from the rare variants GRMs) or their diagonal values (<0.7 or >1.3) or both (removing samples based on the diagonals first then off-diagonals). (A) GREML-LDMS estimates for height fitting either 20 HM3 PCs or 160 PCs computed from all WGS bins. Estimates without any filtering 0.58 – 0.70 (SE 0.08), estimates when filtering on off-diagonals 0.59 – 0.71 (SE 0.08 – 0.09), estimates when filtering on diagonals 0.71 – 0.81 (SE 0.09), estimates when filtering on both 0.71 – 0.82 (SE 0.09 – 0.10). (B) GREML-LDMS estimates for BMI without filtering 0.31 – 0.36 (SE 0.08), estimates when filtering on off-diagonals 0.29 – 0.35 (SE 0.09), estimates when filtering on diagonals 0.31 – 0.37 (SE 0.09 – 0.10), estimates when filtering on both 0.31 – 0.38 (SE 0.10).*

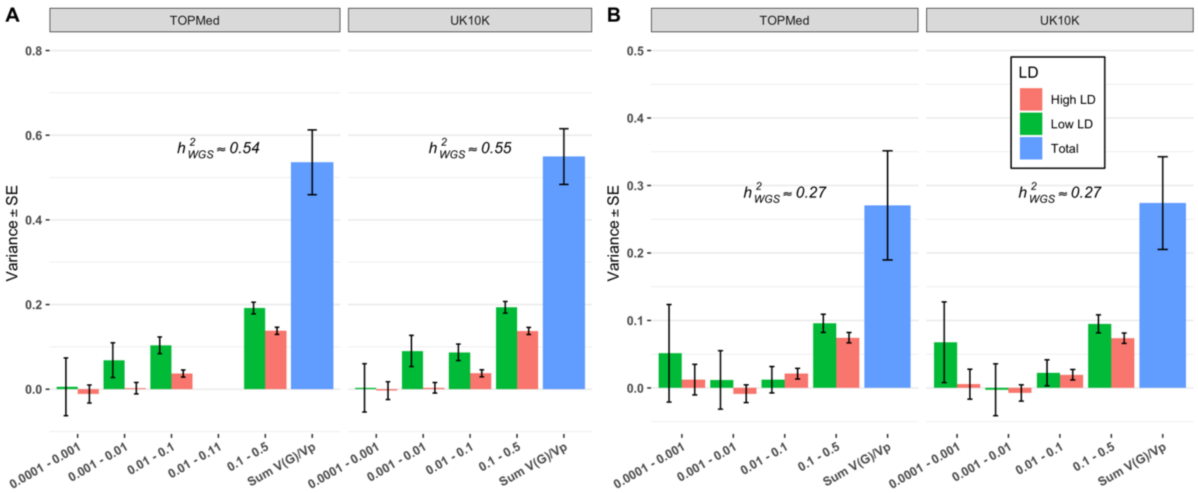


*Supplementary Figure 24: GREML-LDMS estimates from variants in common (~20.0M variants) between TOPMed and UK10K datasets stratified in 8 bins according to variants MAF and LD properties using either TOPMed or UK10K LD and MAF reference, corrected for 160 PCs from WGS independent variants. (A) Estimates of $h^2_{WGS}$ for height (~0.54 – 0.55 (SE 0.07 – 0.08)) or (B) BMI (~0.27 (SE 0.07 – 0.08)) are similar and independent of the LD and MAF reference. The number of variants in each of the 7 MAF bins (twice the number in each LD bin) is, from the lowest to highest MAF bins, 9.1.M, 4.5M, 3.3M and 3.1M, respectively.*

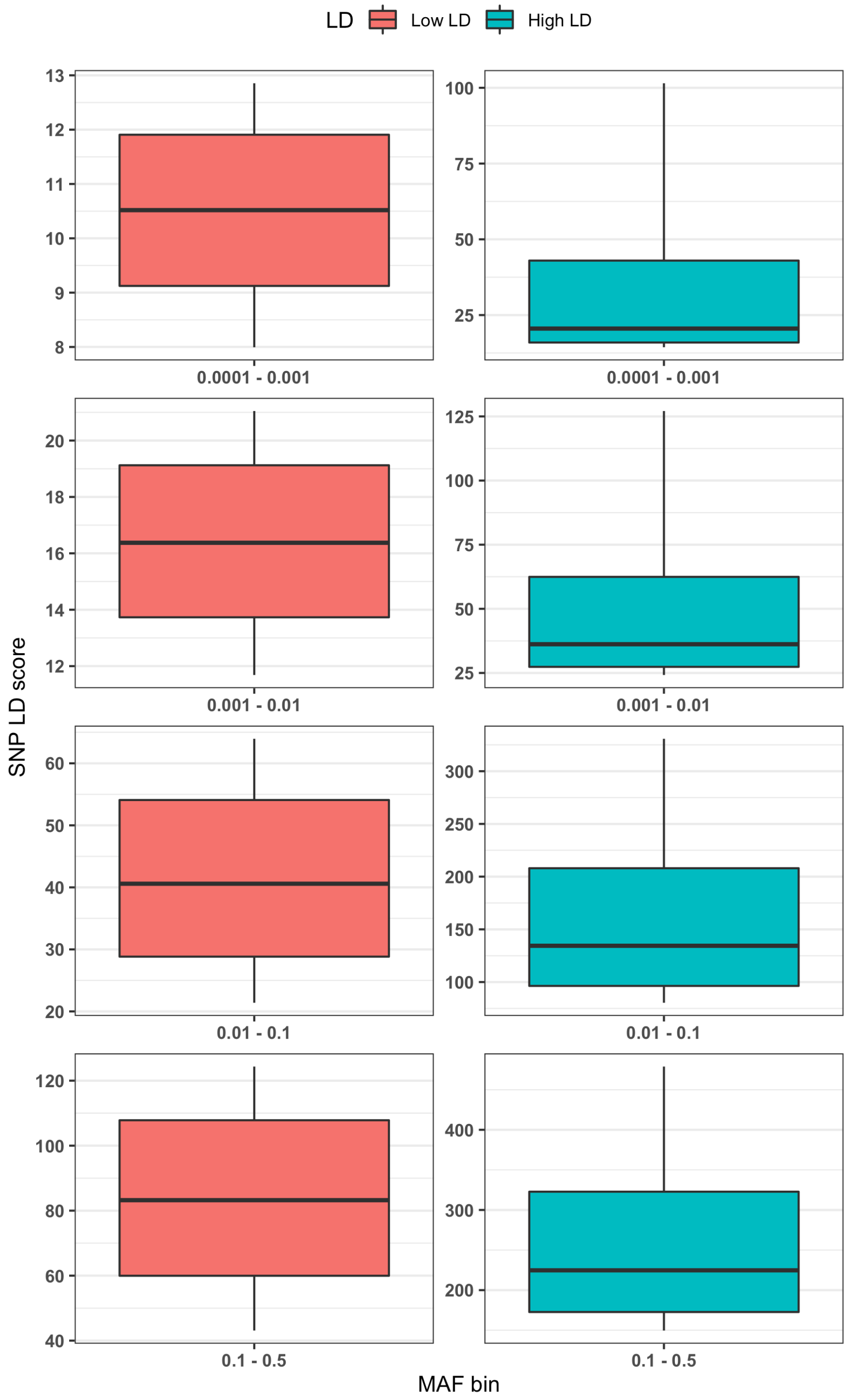

*Supplementary Figure 25: Distribution of SNP LD score for each MAF/LD bin for N=25,465 samples. For more clarity, boxplot outliers are not shown here. Most of the extreme outliers are from rare SNPs in very high LD bins.*

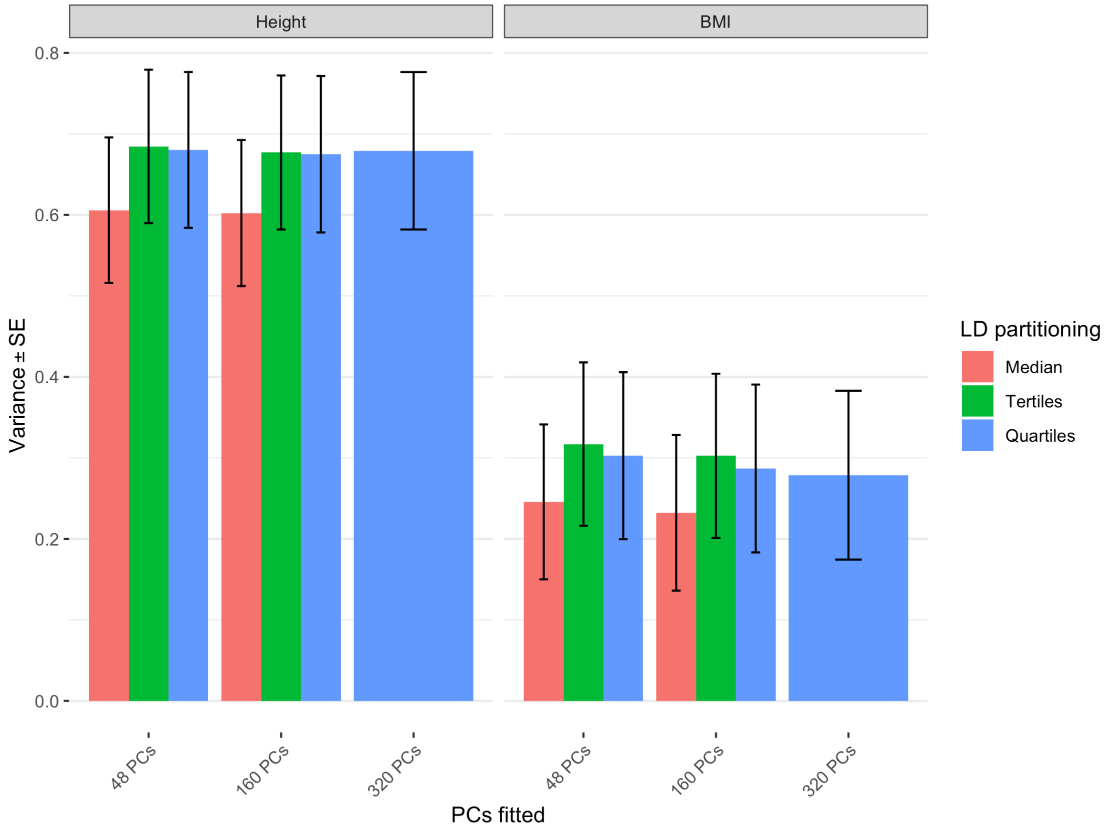


*Supplementary Figure 26: Summary of the impact of different LD grouping strategies on total heritability estimates for N=25,465 samples using 2, 3 or 4 LD grouping for each MAF bin correcting by 48/160/320 PCs computed from WGS independent SNPs. Estimates for height 0.60 – 0.68 (SE 0.09 – 0.10), estimates for BMI 0.23 – 0.32 (SE 0.10).*

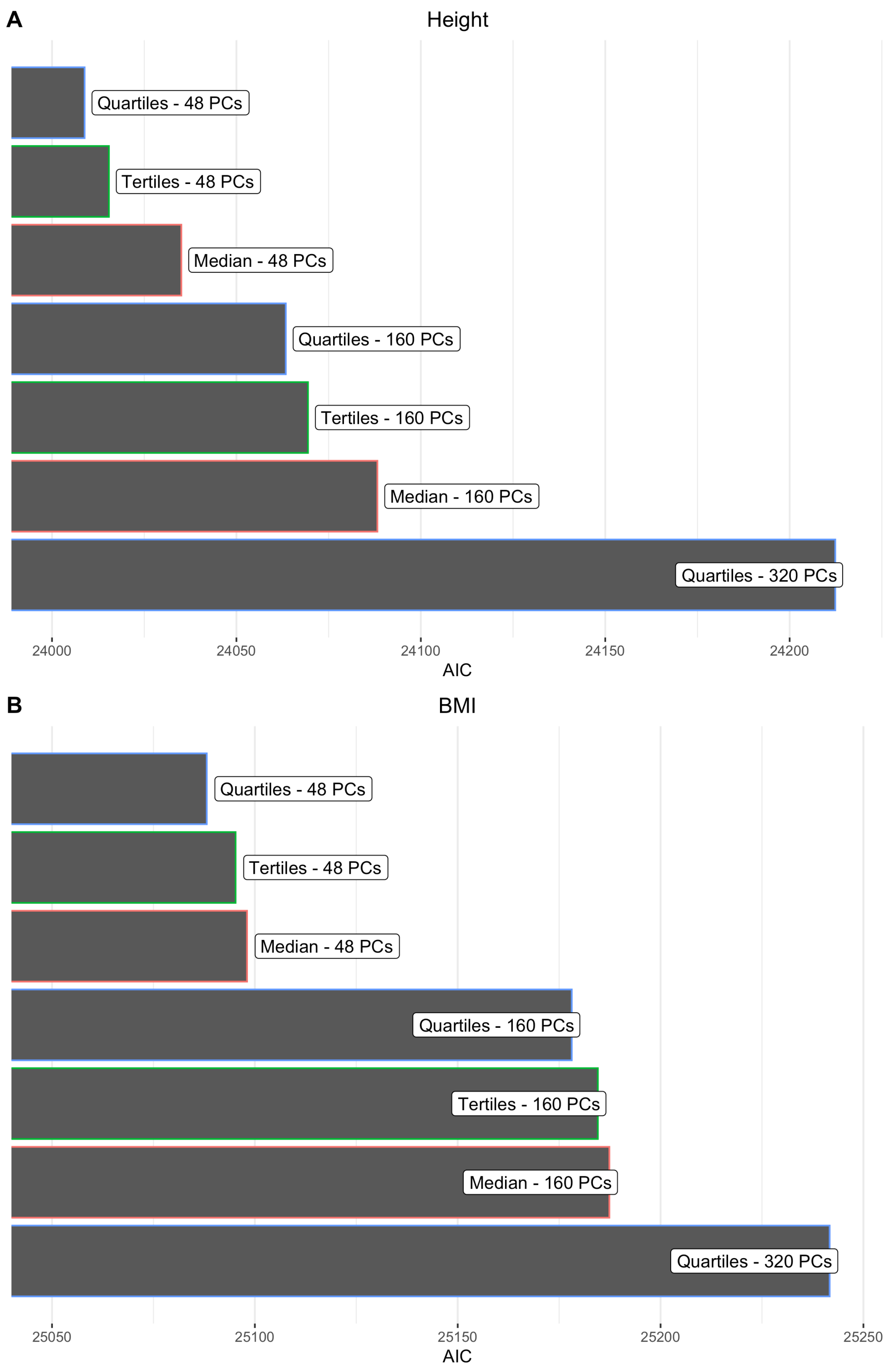


*Supplementary Figure 27: AIC fitting different models, with either 2, 3 or 4 LD bins and 40, 160 or 320 PCs for height (A) and BMI (B). The lower AIC indicates a better fitting model.*

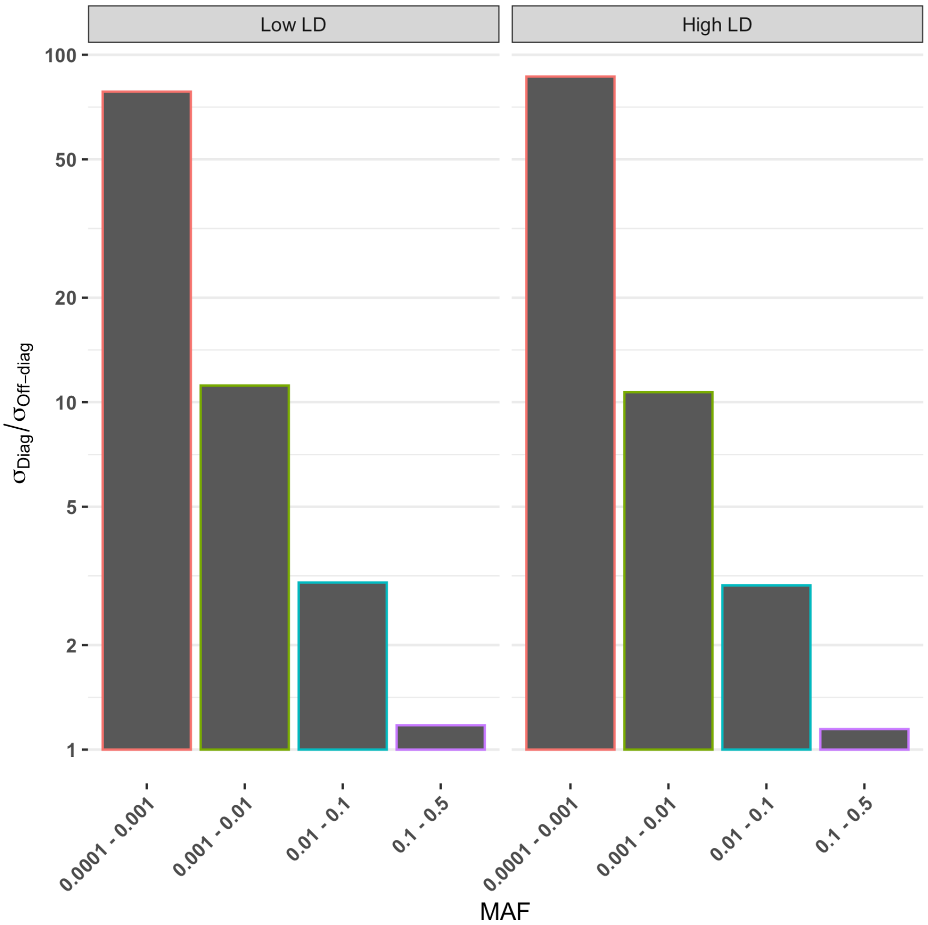


*Supplementary Figure 28: Ratio of GRM diagonal elements variance over GRM off-diagonal elements variance, for each MAF and LD bin using N=25,465 unrelated Europeans samples. GRMs were computed using the ratio of average method.*

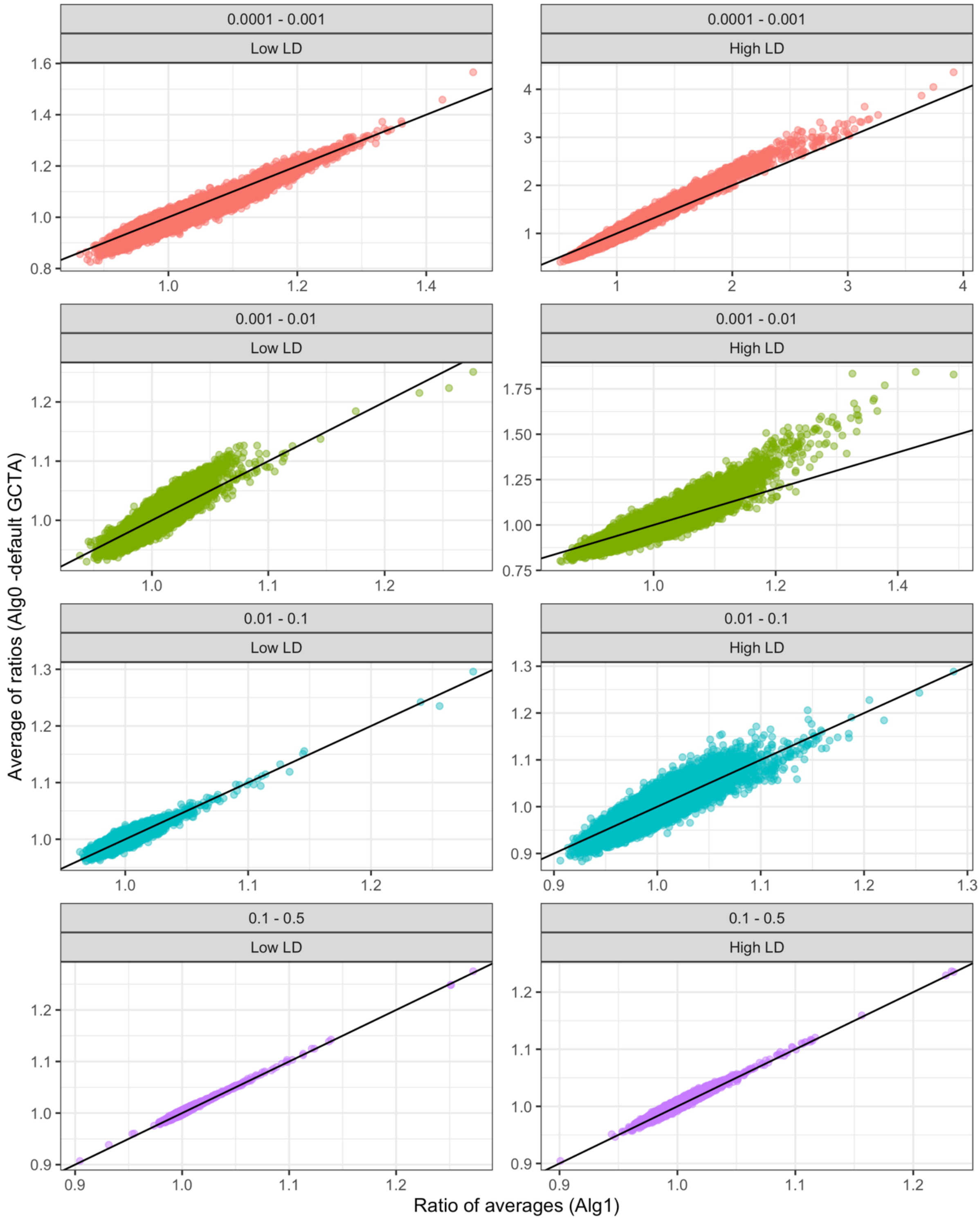


*Supplementary Figure 29: Difference in diagonal elements for the ratio of averages and average of ratios GRM estimators. Diagonal for each MAF/LD bins are plotted using 28,755 unrelated samples of European ancestry.*

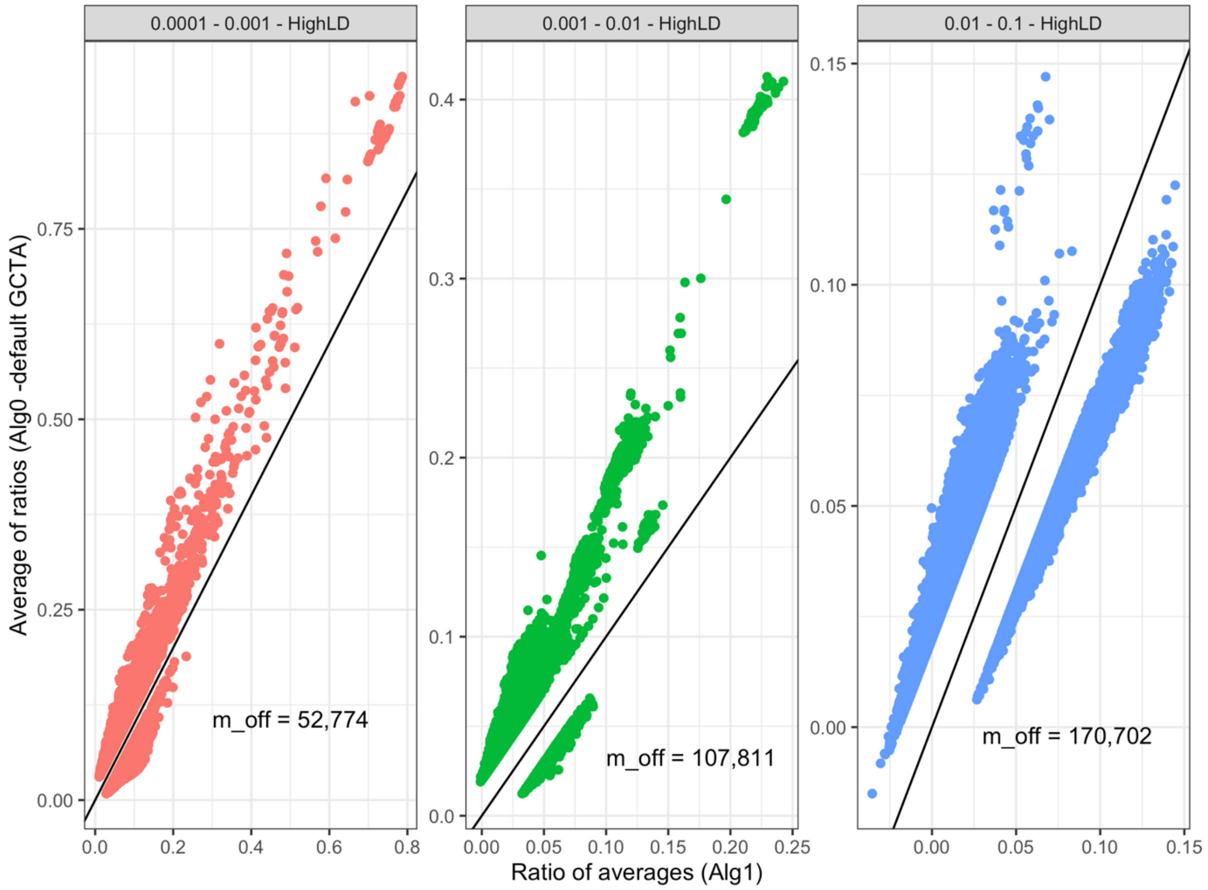


*Supplementary Figure 30: Impact of the GRM estimator on off-diagonal elements using 28,755 unrelated samples of European ancestry, only pairs with a relatedness difference larger than 2% are shown. Total number of diverging pairs is indicated within each window. The total number of pairs in the GRM ~ 413M. Low LD bins and common ones (0.1 < MAF < 0.5) did not present any difference larger than 2%.*

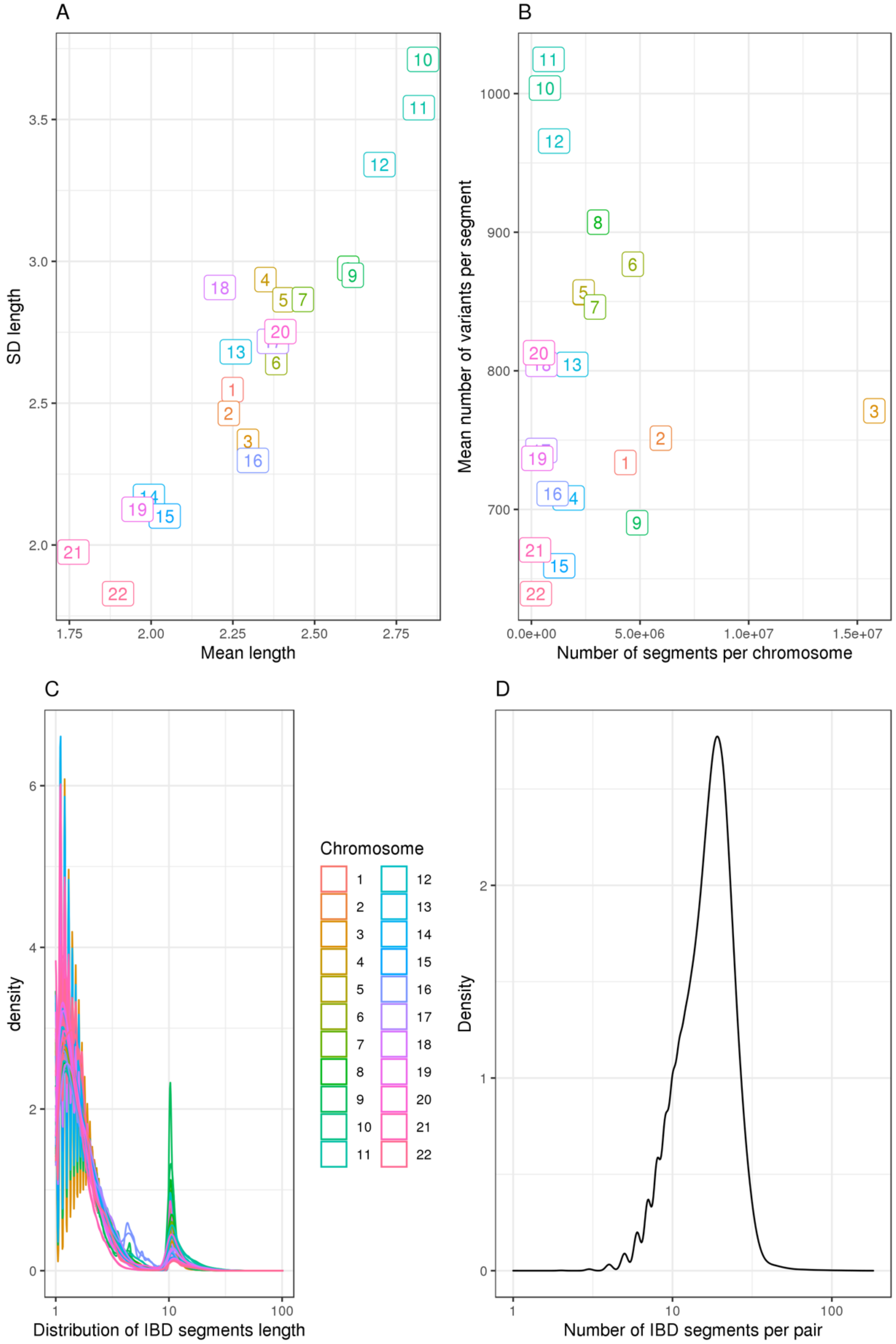


*Supplementary Figure 31: Information on IBD segments distribution among N=28,755 unrelated Europeans. (A) Mean segment length per chromosome and length SD. (B) Number of segments and mean number of variants per segment. (C) Distribution (log10 scale) of the length of IBD segments per chromosome. (D) Distribution of the number of segments per unique sample pair sharing a segment IBD (log10 scale).*

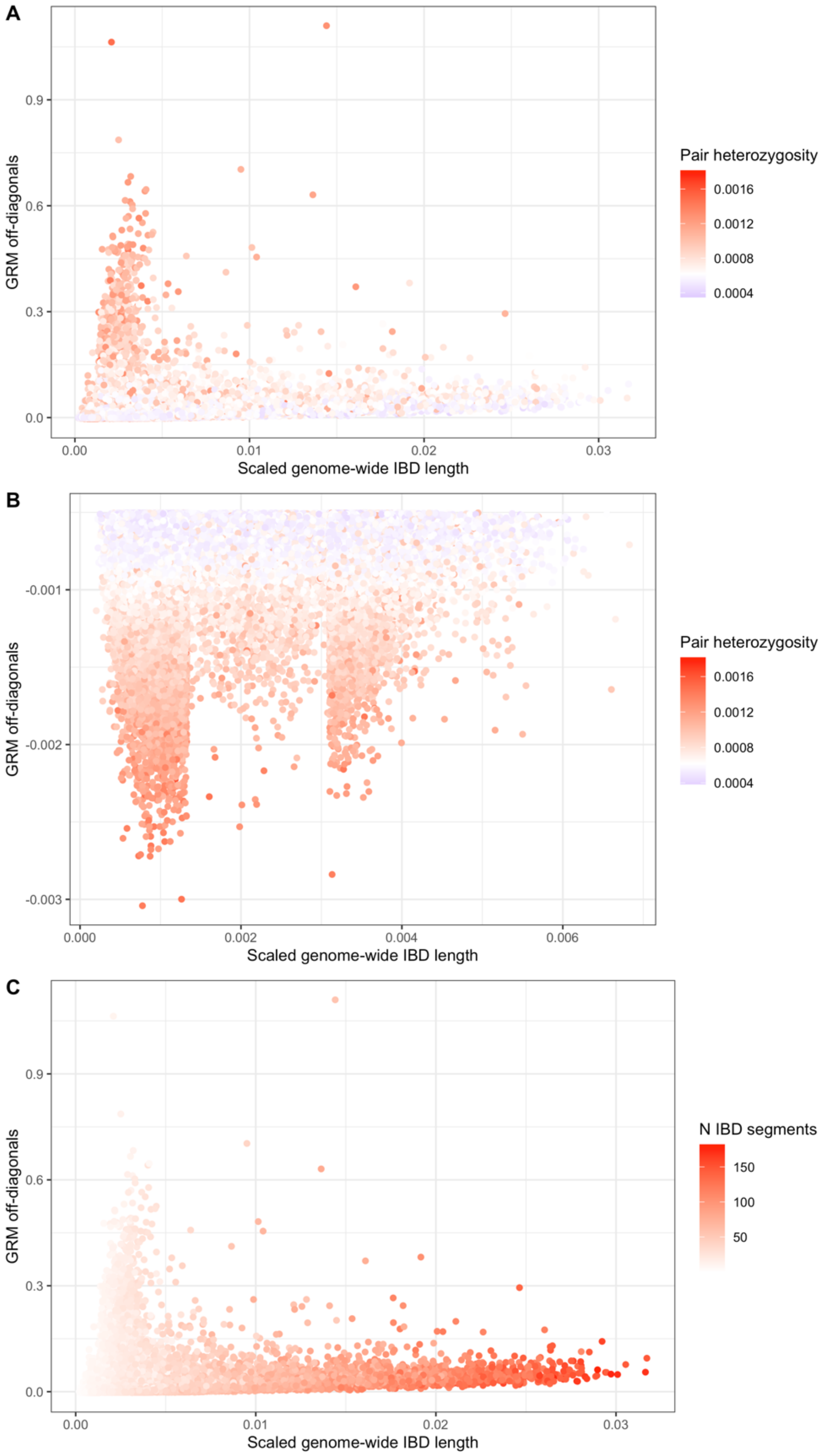


*Supplementary Figure 32: Investigation on the relationship between genome-wide length shared IBD and GRM off-diagonal elements for variants 0.0001 < MAF < 0.001 in high LD. (A) Each sample pair is coloured by their bin specific mean individual heterozygosity rate. (B) Similar to (A), but showing only pairs with low relatedness value. (C) Each sample pair is coloured by the number segments shared IBD.*

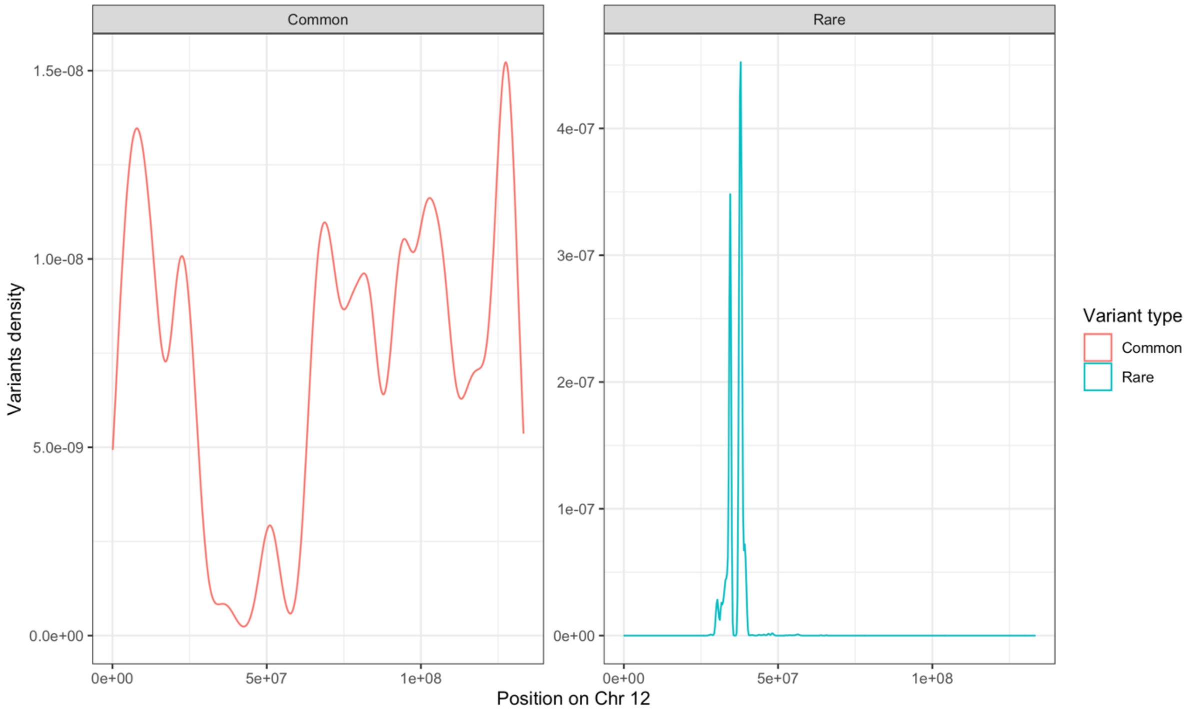


*Supplementary Figure 33: Variant density for a pair of samples showing extreme relatedness in the off-diagonal of the GRM from variants of 0.0001 < MAF < 0.001 in high LD. Common variants (0.1 < MAF < 0.5) distribution on chromosome 12 (left panel) filtering out pair-specific heterozygotes variants. Rare variants distribution (right panel) with a MAC 2 threshold for the pair considered. Note that over 99% of shared heterozygotes rare variants were located on chromosome 12. The lack of oppositive homozygotes variants in common SNPs, at the same location of the pair specific rare heterozygotes variants suggests a region shared IBD.*

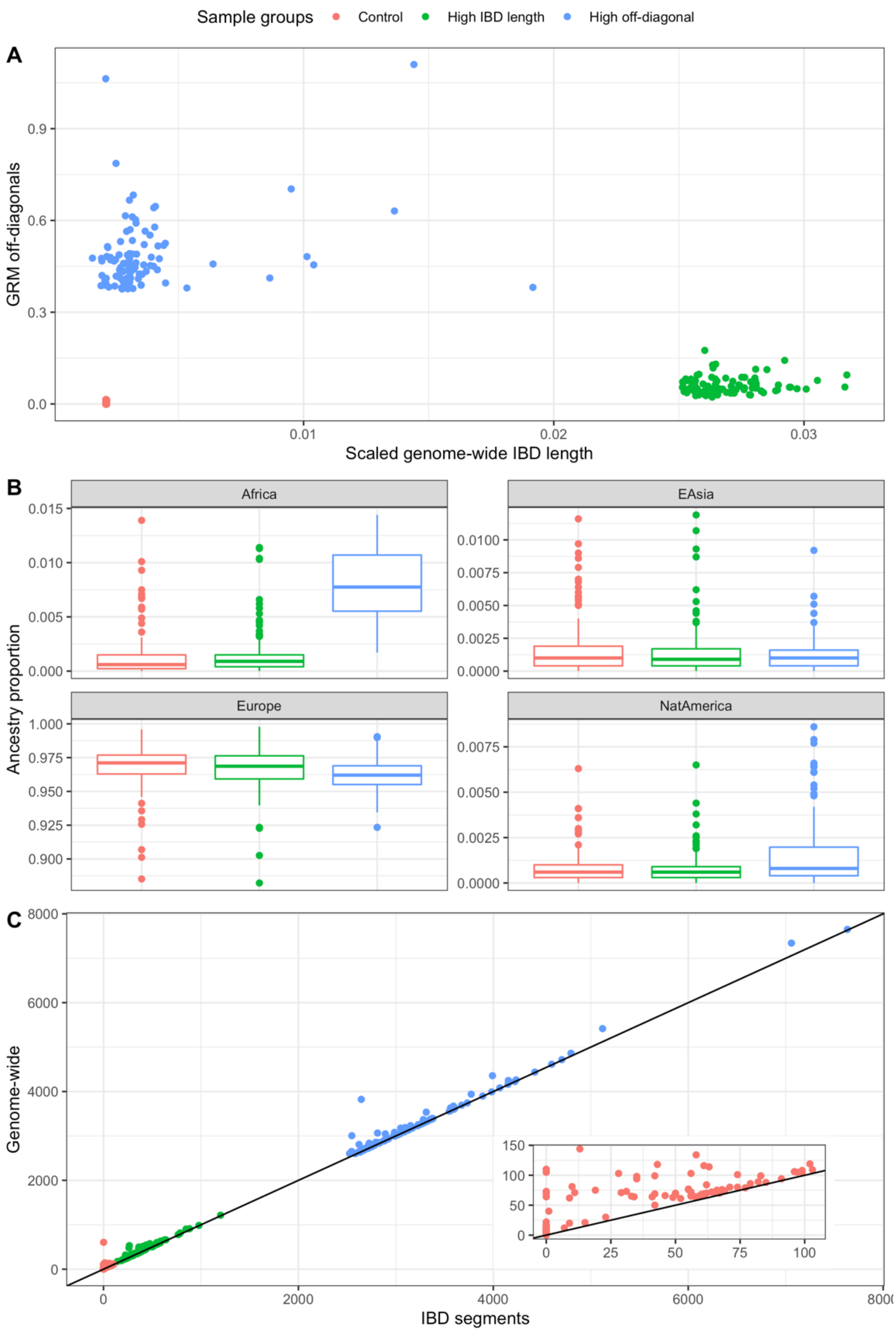


*Supplementary Figure 34: Investigating the characteristics of extreme GRM pairs. (A) Top 100 pairs with either the largest proportion of genome shared IBD (green), the largest off-diagonal values for rare variants in high LD (blue) or 100 pairs selected near the median (red). Most of the high off-diagonal values do not present large segments shared IBD. (B) Ancestry proportion of samples within each group. The highly related samples show an increase of genome-wide African ancestry compared to the two other groups. (C) Origin of the pair-specific heterozygotes variants considering only the very rare variants in high LD, either coming from the segments shared IBD by each pair or coming from the shared SNPs genome-wide. Embedded plot shows the control pairs only.*

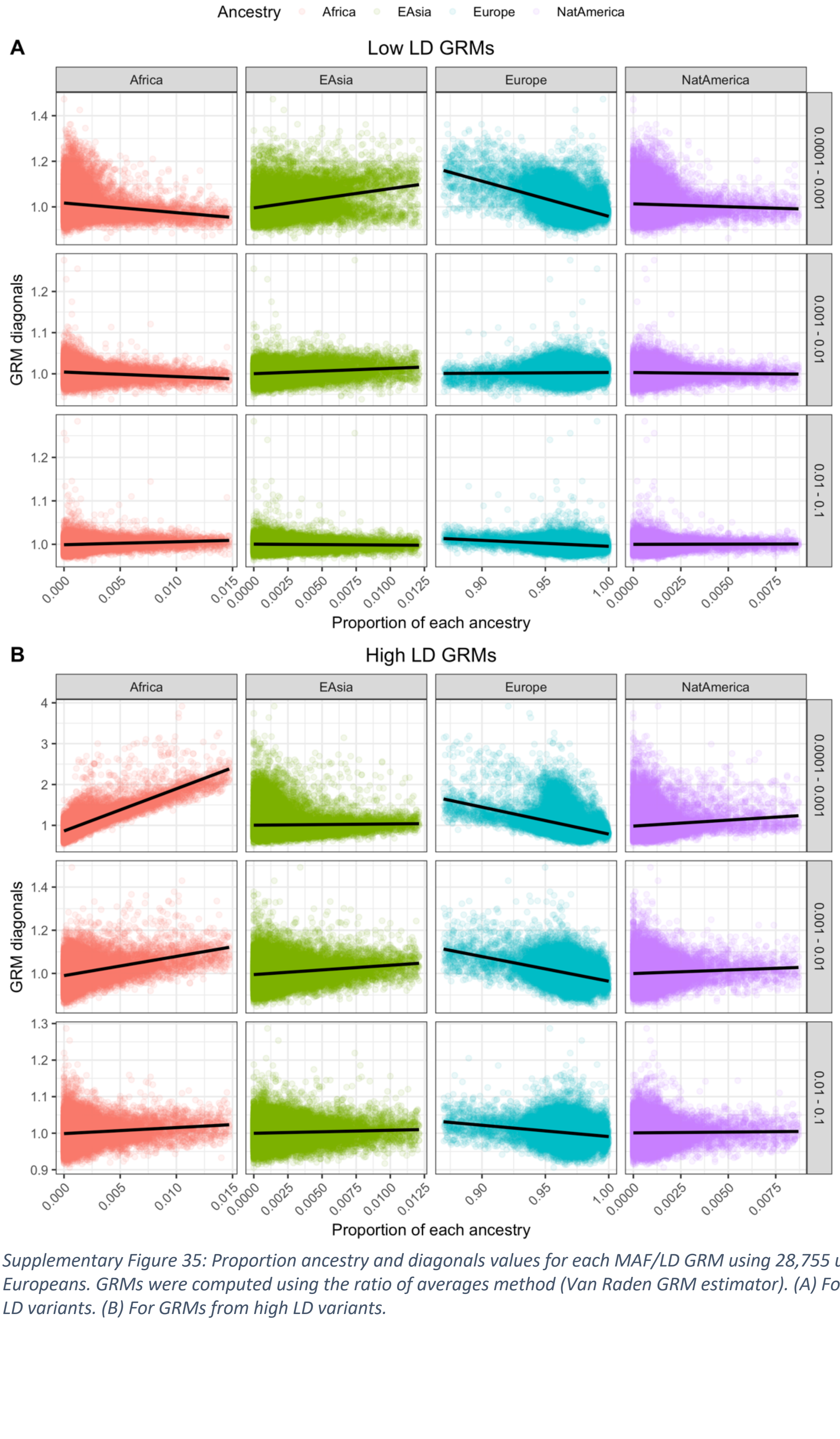


*Supplementary Figure 35: Proportion ancestry and diagonals values for each MAF/LD GRM using 28,755 unrelated Europeans. GRMs were computed using the ratio of averages method (Van Raden GRM estimator). (A) For GRMs from low LD variants. (B) For GRMs from high LD variants.*
